# Learning interpretable cellular and gene signature embeddings from single-cell transcriptomic data

**DOI:** 10.1101/2021.01.13.426593

**Authors:** Yifan Zhao, Huiyu Cai, Zuobai Zhang, Jian Tang, Yue Li

## Abstract

The advent of single-cell RNA sequencing (scRNA-seq) technologies has revolutionized transcriptomic studies. However, large-scale integrative analysis of scRNA-seq data remains a challenge largely due to unwanted batch effects and the limited transferabilty, interpretability, and scalability of the existing computational methods. We present single-cell Embedded Topic Model (scETM). Our key contribution is the utilization of a transferable neural-network-based encoder while having an interpretable linear decoder via a matrix tri-factorization. In particular, scETM simultaneously learns an encoder network to infer cell type mixture and a set of highly interpretable gene embeddings, topic embeddings, and batch effect linear intercepts from multiple scRNA-seq datasets. scETM is scalable to over 10^6^ cells and confers remarkable cross-tissue and cross-species zero-shot transfer-learning performance. Using gene set enrichment analysis, we find that scETM-learned topics are enriched in biologically meaningful and disease-related pathways. Lastly, scETM enables the incorporation of known gene sets into the gene embeddings, thereby directly learning the associations between pathways and topics via the topic embeddings.

## Background

Advances in high-throughput sequencing technologies [1] provide an unprecedented opportunity to profile the individual cells’ transcriptomes across various biological and pathological conditions, and have spurred the creation of several atlas projects [2–5]. Emerged as a key application of single-cell RNA sequencing (scRNA-seq) data, unsupervised clustering allows for cell-type identification in a data-driven manner. Flexible, scalable, and interpretable computational methods are crucial for unleashing the full potential from the wealth of single-cell datasets and translating the transcription profiles into biological insights. Despite considerable progress made on clustering method development for scRNA-seq data analysis [6–16], several challenges remain.

First, compared to bulk RNA-seq, scRNA-seq data commonly exhibit higher noise levels and drop-out rates, where the data only captures a small fraction of a cell’s transcriptome [17]. Changes in gene expression due to experimental design, often referred to as batch effects [18], can have a large impact on clustering [12, 18–20]. If not properly addressed, these technical artefacts may mask the true biological signals in cell clustering.

Second, the partitioning of the cell population alone is insufficient to produce biological interpretation. The annotations of the cell clusters require extensive manual literature search in practice and the annotation quality may be dependent on users’ domain knowledge [20]. Therefore, an interpretable and flexible model is needed. In the current work, we consider model interpretability as whether the model parameters can be directly used to associate the input features with latent factors or target outcomes. Latent topic models are a popular approach in mining genomic and healthcare data [21–23] and are increasingly being used in the scRNA-seq literature [24]. Specifically, in topic modeling, we infer the topic distribution for both the samples and genomic features by decomposing the samples-by-features matrix into samples-by-topics and topics-by-features matrices, which also be viewed as a probabilistic non-negative factorization (NMF) [25]. Importantly, the top genes under each latent topic can reveal the gene signatures for specific cellular programs, which may be shared across cell types or exclusive to a particular cell type. Traditionally, the latter are detected via differential expression analysis at individual gene levels, which has limited statistical power in scRNA-seq data analysis because of the sparse gene counts, small number of unique biological samples, and the burdens of multiple testings.

Third, model transferrability is an important consideration. We consider a model as *transferable* if the learned knowledge manifested as the model parameters could benefit future data modeling. In the context of scRNA-seq data analysis, it translates to learning feature representations from one or more large-scale reference datasets and applying the learned representations to a target dataset [26, 27]. If the model is not further trained on the target dataset, the learning setting called *zero-shot transfer learning*. A model that can successfully separate cells of distinct cell types that are not present in the reference dataset implies that the model has learned some meaningful abstraction of the cellular programs from the reference dataset such that it can generalize to annotating new cell types of different kinds. An analogy would be that someone who has learned how to distinguish triangles from rectangles may also be able to distinguish squares from circles. As the number and size of scRNA-seq datasets continue to increase, there is an increasingly high demand for efficient exploitation and knowledge transfer from the existing reference datasets.

Several recent methods have attempted to address these challenges. Seurat [7] uses canonical correlation analysis to project cells onto a common embedding, then identifies, filters, scores, and weights anchor cell pairs between batches to perform data integration. Harmony [28] iterates between maximum diversity clustering and a linear batch correction based on the mixture-of-experts model. Scanorama [10] performs all-to-all dataset matching by querying nearest neighbors of a cell among all remaining batches, after which it merges the batches with a Gaussian kernel to form a single cell panorama. These methods rely on feature (gene) selection and/or dimensionality reduction methods; otherwise they can not scale to compendium-scale reference [2] or cohort-level single-cell transcriptome data [29] or are sensitive to the noise inherent to scRNA-seq count data. They are also non-transferable, meaning that the knowledge learned from one dataset cannot be easily transferred through model parameter sharing to benefit the modeling of another dataset. NMF approaches such as UNCURL [30] works only with one scRNA-seq dataset. LIGER [9] uses integrative NMF to jointly factorize multiple scRNA-seq matrices across conditions using genes as the common axis, linking cells from different conditions by a common set of latent factors also known as metagenes. LIGER is weakly transferable in the sense that the global metagenes-by-genes matrix can be recycled as initial parameters when modeling a new target dataset, whereas the cells-by-metagenes and the final metagenes-by-genes must be further computed and updated by iterative numerical optimization to fit the target dataset.

Deep learning approaches, especially autoencoders, have demonstrated promising performance in scRNA-seq data modeling. scAlign [15] and MARS [31] encode cells with non-linear embeddings using autoencoders, which is naturally transferable across datasets. While scAlign minimizes the distance between the pairwise cell similarity at the embedding and original space, MARS looks for latent landmarks from known cell types to infer cell types of unknown cells. Variational autoencoders (VAE) [32] is an efficient Bayesian framework for approximating intractable posterior distribution using proposed distribution parameterized by neural networks. Several recent studies have tailored the original VAE framework towards modeling single-cell data. Single-cell variational inference (scVI) [6] models library size and takes into account batch effect in generating cell embeddings. scVAE-GM [11] changed the prior distribution of the latent variables in the VAE from Gaussian to Gaussian mixture model, adding a categorical latent variable that clusters cells. Lotfollahi *et al*. developed a VAE model called scGen to infer the expression difference due to perturbation conditions by latent space interpolation [26]. A key drawback for these VAE models is the lack of interpretability – post hoc analyses are needed to decipher the learned model parameters and distill biological meaning from the learned network parameters. To improve interpretability, Svensson *et al.* (2020) developed a linear decoded VAE (hereafter referred to as scVI-LD) as a part of the scVI software [14].

In this paper, we present *single-cell Embedded Topic Model* (scETM), a generative topic model that facilitates integrative analysis of large-scale single-cell transcriptomic data. Our key contribution is the utilization of a transferable neural-network-based encoder while having an interpretable linear decoder via a matrix tri-factorization. The scETM simultaneously learns the encoder network parameters and a set of highly interpretable gene embeddings, topic embeddings, and batch-effect linear intercepts from scRNA-seq data. The flexibility and expressiveness of the encoder network enable scETM to model extremely large scRNA-seq datasets without the need of feature selection or dimension reduction. By tri-factorizing cells-genes matrix into cells-by-topics, topics-by-embeddings, and embeddings-by-genes, we are able to incorporate existing pathway information into gene embeddings during the model training to further improve interpretability. This is a salient feature compared to related methods such as scVI-LD. It allows scETM to simultaneously discover interpretable cellular signatures and gene markers while integrating scRNA-seq data across conditions, subjects and/or experimental studies.

We demonstrate that scETM offers state-of-the-art performance in clustering cells into known cell types across a diverse range of datasets with desirable runtime and memory requirements. We also demonstrate scETM’s capability of effective knowledge transfer between different sequencing technologies, between different tissues, and between different species. We then use scETM to discover biologically meaningful gene expression signatures indicative of known cell types and pathophysiological conditions. We analyze scETM-inferred topics and show that several topics are enriched in cell-type-specific or disease-related pathways. Finally, we directly incorporate known pathway-gene relationships (pathway gene set databases) into scETM in the form of gene embeddings, and use the learned pathway-topic embedding to show the pathway-informed scETM (p-scETM)’s capability of learning biologically meaningful information.

## Results

### scETM model overview

We developed scETM to model scRNA-seq data across experiments or studies, which we term as batches (**Fig. 1a** and **Supp. Fig. S1a**). Adapted from the Embedded Topic Model (ETM) [33], scETM inherits the benefits of topic models, and is effective for handling large and heavy-tailed distribution of word frequency. In the context of scRNA-seq data analysis, each sampled single-cell transcriptome is provided as a vector of normalized gene counts to a two-layer fully-connected neural network (i.e., encoder; see detailed architecture in **Supp. Fig. S1c**), which infers the topic mixing proportions of the cell. The trained encoder on a reference scRNA-seq data can be used to infer topic mixture of unseen scRNA-seq data collected from different tissues or species (**Fig. 1b**).

**Figure 1:**
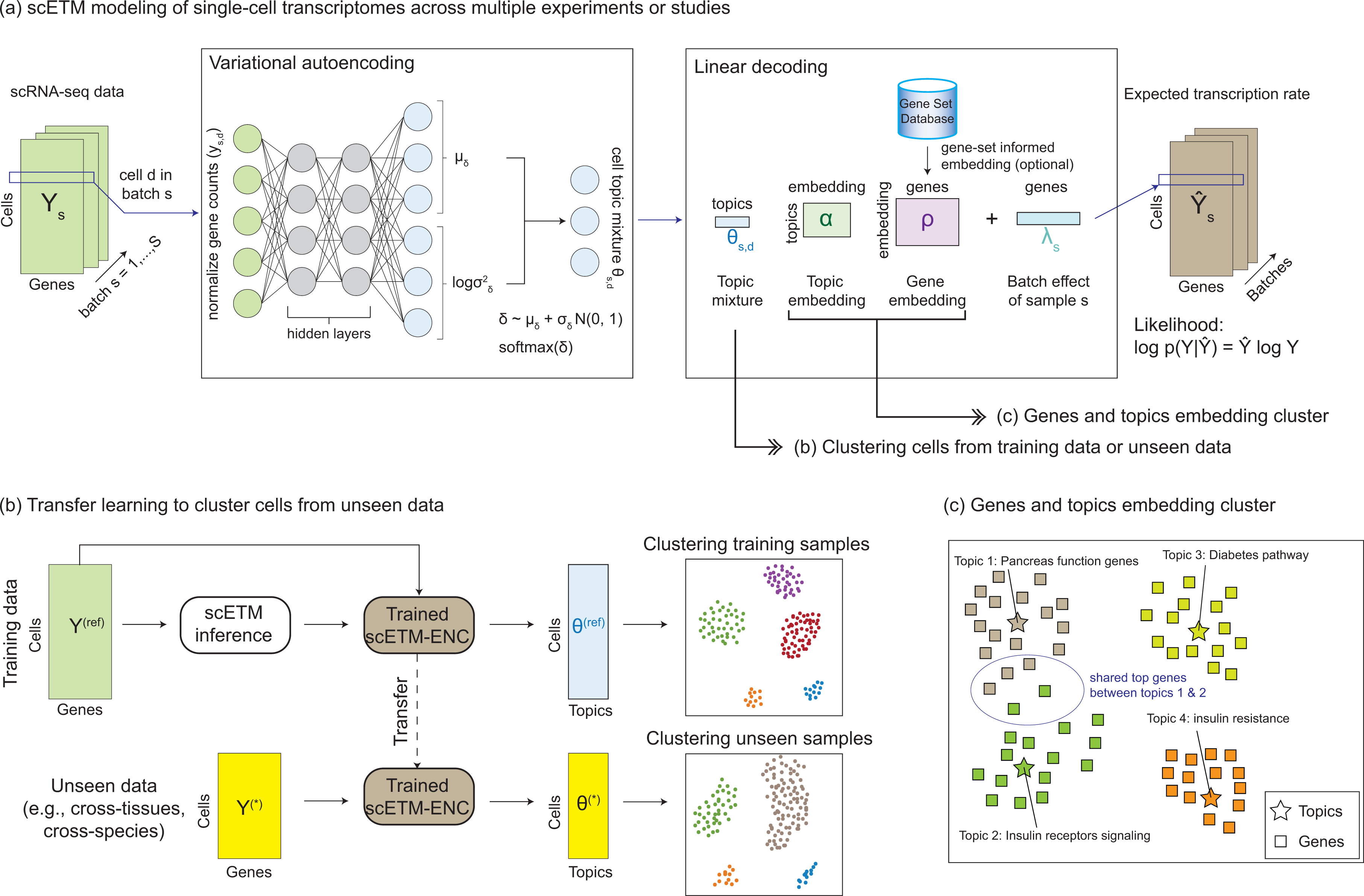
scETM model overview. **(a)** scETM training. Given as input the scRNA-seq data matrices across multiple experiments or studies (i.e., batches), scETM models the single-cell transcriptomes using an embedded topic modeling approach. Each scRNA-seq profile serves as an input to a variational autoencoder (VAE) as the normalized gene counts. The encoder network produces a stochastic sample of the latent topic mixture (**θ***_s,d_* for batch *s* = 1*, … , S* and cell *d* = 1*, … , N_s_*), which can be used for clustering cells (see panel b). The linear decoder learns topic embedding and gene embedding, which can be used to analyze cellular programs via enrichment analyses (see panel c). **(b)** Workflow used to perform zero-shot transfer learning. The trained scETM-encoder on a reference scRNA-seq dataset is used to infer the cell topic mixture **θ***^∗^* from an unseen scRNA-seq dataset *without training* them. The resulting cell mixtures are then visualized via UMAP visualization and evaluated by standard unsupervised clustering metrics using the groundtruth cell types. **(c)** Exploring gene embeddings and topic embeddings. Because the genes and topics share the same embedding space, we can explore their connections via UMAP visualization or annotate each topic via enrichment analyses using known pathways.

For interpretability, we use a linear decoder with the gene and topic embeddings as the learnable parameters. Specifically, we factorize the cells-by-genes count matrix into a cells-by-topics matrix **θ** (inferred by the encoder), topics-by-embedding **α**, and embedding-by-genes **ρ** matrices (**Supp. Fig. S1b**). This tri-factorization design allows for exploring the relations among cells, genes, and topics in a highly interpretable way. To account for biases across conditions or subjects, we introduce an optional batch correction parameter **λ**, which acts as a linear intercept term in the categorical softmax function to alleviate the burden of modeling batch effects from the encoder to let it focus on inferring biologically meaningful cell topic mixture **θ***_d_*. The encoder and embedding learning is performed by an amortized variational inference algorithm to maximize the evidence lower bound (ELBO) of the marginal categorical likelihood of the scRNA-seq counts [32]. Compared to scVI-LD [14], the linear decoder component that learns a common embeddings for both topics and genes offers more flexibility and interpretability and overall better performance as we demonstrate next (**Fig. 1c**). Details of the scETM algorithm and implementation are described in **Methods**.

### Data Integration

We benchmarked scETM, along with 7 state-of-the-art single-cell clustering or integrative analysis methods – scVI [6], scVI-LD [14], Seurat v3 [7], scVAE-GM [11], Scanorama [10], Harmony [28] and LIGER [9], on 6 published datasets, namely Mouse Pancreatic Islet (MP) [34], Human Pancreatic Islet (HP) [7], *Tabula Muris* (TM) [3], Alzheimer’s Disease dataset (AD) [35], Major Depressive Disorder dataset (MDD) [29], and Mouse Retina (MR) [36, 37] (**Supplementary Methods**). Across all datasets, scETM stably delivered competitive results especially among the transferable and interpretable models, while others methods fluctuate across different datasets in terms of Adjusted Rand Index (ARI) and Normalized Mutual Information (NMI) (**Table** 1; **Supp. Table S1**). Overall, Harmony and Seurat have slightly higher ARIs than scETM, with trade-offs of model transferrability, interpretability, and/or scalability, which we investigate in the following sections.

**Table 1:** Model properties and unsupervised clustering performance on data integration tasks. The clustering performance is measured by Adjusted Rand Index (ARI) between ground truth cell types and Leiden [75] clusters. scETM performances with or without the linear batch correction (*scETM, scETM λ*) are both reported. scETM+adv is scETM plus adversarial network loss to further correct batch effects. Batch variables include strain, sequencing technologies (“Tech.”) and individuals (“Ind.”). NA is reported for models that did not converge. Experimental details are described in the **Methods** section. We ran Seurat on all datasets with integration turned on whenever applicable.

|  | Transferable | Interpretable | MP | HP | TM | AD | MDD | MR |
| --- | --- | --- | --- | --- | --- | --- | --- | --- |
| Harmony |  |  | 0.969 | 0.955 | 0.705 | 0.994 | 0.784 | 0.763 |
| Scanorama |  |  | 0.915 | 0.859 | 0.542 | 0.997 | 0.587 | 0.780 |
| Seurat |  |  | 0.944 | 0.968 | 0.676 | 0.991 | 0.550 | 0.781 |
| scVAE-GM | ✓ |  | 0.805 | NA | NA | 0.997 | 0.563 | 0.778 |
| scVI | ✓ |  | 0.932 | 0.759 | 0.670 | 0.991 | 0.541 | 0.783 |
| LIGER | ✓ | ✓ | 0.914 | 0.911 | 0.591 | 0.894 | 0.704 | 0.714 |
| scVI-LD | ✓ | ✓ | 0.875 | 0.656 | 0.608 | 0.989 | 0.666 | 0.718 |
| scETM | ✓ | ✓ | 0.946 | 0.943 | 0.761 | 0.996 | 0.717 | 0.859 |
| scETM-λ | ✓ | ✓ | 0.851 | 0.474 | 0.629 | 0.996 | 0.719 | 0.773 |
| scETM+adv | ✓ | ✓ | 0.944 | 0.946 | 0.704 | 0.993 | 0.717 | 0.772 |
| Batch Effect |  |  | Strain | Tech. | Tech. | Ind. | Ind. | Studies |

We further experimented the same scETM without the batch correction term, namely scETM-*λ*. Compared to the *λ*-ablated model, the full scETM model confers higher ARI in 3 out of the 5 datasets (**Table 1**) and higher NMI in 4 out of the 5 datasets (**Supp. Table S1**). Improvement over the Human Pancreas (HP) dataset is remarkably high, implying an effective correction of the confounder due to the scRNA-seq technology differences. We observe no improvement in the AD dataset in terms of both ARI and NMI and small improvement in MDD only in terms of NMI. This implies a lesser concern of batch effects from only the individual brain sample donors, with all data being collected by the same technology in a single study.

We also evaluated the batch mixing aspect of scETM and other methods using k-nearest-neighbor Batch-Effect Test (kBET) [18] (**Table** 2) and examined to what extent scETM’s batch mixing performance can be improved by introducing an adversarial loss term to scETM (**Methods**) [38]. Briefly, we used a discriminator network (a two-layer feed-forward network) to predict batch labels using the cell topic mixture embeddings generated by the encoder network, and directed the encoder network to fool the discriminator. We observe notable improvement on kBET with similar ARI and NMI scores (**Table** 1 and **Supp. Table** S1, row “scETM+adv”) at the cost of up to 50% more running time. This shows the *expandability* of scETM. For the subsequent analyses, we opted to use the results from scETM (without the adversarial loss but with the linear batch correction *λ*) because of its simpler design, scalability, comparable ARI scores, and less aggressive batch correction (see below).

**Table 2:** Batch correction performance on data integration tasks. The batch correction performance is measured by kBET [18]. More details are described in Table 1 caption and **Methods**.

|  | MP | HP | TM | AD | MDD | MR |
| --- | --- | --- | --- | --- | --- | --- |
| Harmony | 0.390 | 0.342 | 0.148 | 0.518 | 0.440 | 0.063 |
| Scanorama | 0.439 | 0.155 | 0.001 | 0.286 | 0.202 | 0.063 |
| Seurat | 0.538 | 0.381 | 0.281 | 0.195 | 0.085 | 0.053 |
| scVAE-GM | 0.233 | NA | NA | 0.073 | 0.051 | 0.033 |
| scVI | 0.516 | 0.140 | 0.056 | 0.434 | 0.313 | 0.073 |
| LIGER | 0.602 | 0.660 | 0.374 | 0.771 | 0.716 | 0.176 |
| scVI-LD | 0.148 | 0.034 | 0.069 | 0.476 | 0.277 | 0.022 |
| scETM+adv | 0.546 | 0.443 | 0.144 | 0.627 | 0.570 | 0.141 |
| scETM | 0.270 | 0.163 | 0.096 | 0.278 | 0.228 | 0.066 |
| scETM$-\lambda$ | 0.217 | 0.000 | 0.000 | 0.122 | 0.066 | 0.017 |
| Batch Effect | Strain | Technology | Technology | Individual | Individual | Studies |

scETM is also robust to architectural and hyperparameter changes, requiring very few or no architecture adaptation or hyperparameter tuning efforts when applied to unseen datasets (**Supp. Table S2**). As a result, we used the same architecture and hyperparameters for all datasets in **Table 1**. We also performed a comprehensive ablation analysis to validate our model choices. The ablation experiment demonstrates the necessity of the key model components, such as the batch effect correction factors *λ* and the batch normalization technique used in training the encoder. Normalizing gene expression scRNA-seq counts as the input to the encoder also improves the performance (**Supp. Table S3**).

Clustering agreement metrics are not the only metrics for evaluating scRNA-seq methods, and are not available to unannotated datasets. Therefore, we also evaluated the negative log-likelihood (NLL) on held-out samples, which is a principled way for model selection without labels. We computed the held-out (10%) NLL. We found that scETM is robust to different architectures in terms of the NLL (**Supp. Table S2**). We also found that ARI and NLL are modestly negatively correlated on the TM dataset (**Supp. Fig. S2**), implying an agreement between the two metrics although this might not be always the case since it highly depends on the cell type labels and the data quality.

To further verify the clustering performance and validate our evaluation metrics, we visualized the cell embeddings using Uniform Manifold Approximation and Projection (UMAP) [39] for some of the datasets (**Supp. Fig. S3,S4,S5,S6,S7,S8**, **Fig. 2**). Together, these results support that scETM effectively captures cell-type-specific information, while accounting for artefacts arising from individual or technological variations.

**Figure 2:**
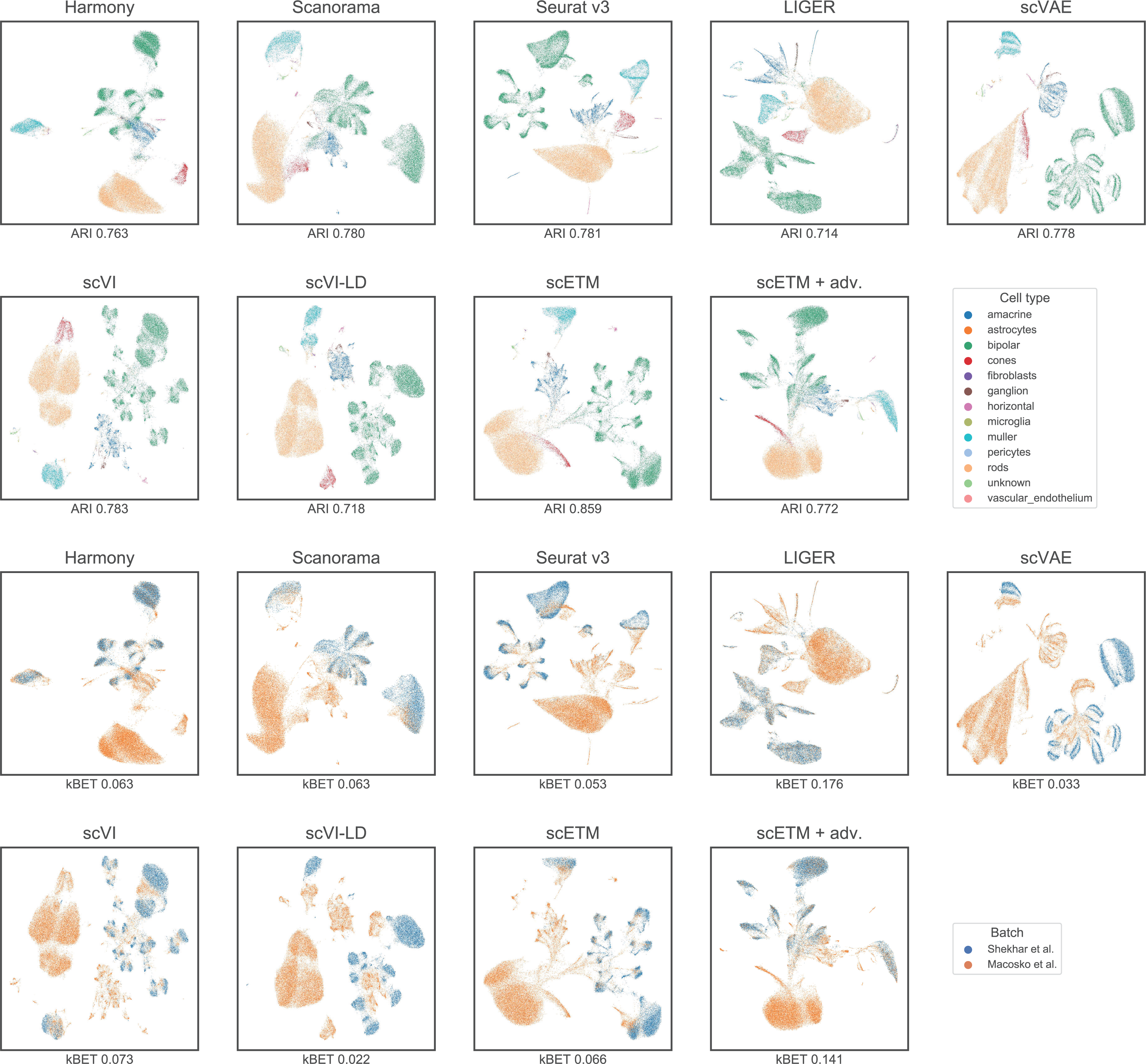
Integration and batch correction on the Mouse Retina dataset. Each panel shows the Mouse Retina cell clusters using UMAP based on the cell embeddings obtained by each of the 9 methods. The cells are colored by cell types in the first two rows and by batches, which are the two source studies, in the last two rows. The ARI and kBET scores of each method are shown below each plot. UMAP visualization for the other 5 datasets are illustrated in **Supp. Fig. S3,S4,S5,S6,S7**.

### Batch overcorrection analysis

Some methods may risk over-correcting batch effects and fail to capture some aspect of biological variations. In the above analysis, we observe that some methods such as LIGER conferred competitive kBET but low ARI, suggesting potential overcorrection of batch effects. To experiment the extent of batch overcorrection by each method, we conducted two experiments using two datasets, namely the Human Pancreas (HP) dataset [7] and the Mouse Retina (MR) dataset [36, 37].

For the HP data, we manually removed *beta* cells from all 5 batches except for batch CelSeq2, resulting in the cell type distributions shown in **Supp. Table S4**. We expect that, if a method is guilty of batch effect overcorrection, it would assign beta cells to other non-beta cell clusters in the latent space by forcing the alignment of different batches. Consequently, such methods would have poor clustering scores. We evaluated all of the methods on this dataset using 3 metrics: ARI, kBET, and average silhouette width (ASW) [40]. Briefly, higher ASW indicates larger distances between cell types and lower distance within cell type in the cell embedding space (**Methods**). We measured the overall ASW as well as the ASW for only the beta cells (i.e., ASW-beta). The results are summarized in **Supp. Table S5**. We observed that scETM struck a good balance between discriminating cell types (ARI: 0.9298; ASW: 0.3525; ASW-beta: 0.5370) and integrating different batches (kBET: 0.1247). Adding the adversarial loss to the scETM (i.e., scETM+adv) increased kBET from 0.1247 to 0.3445 (while maintaining ARI above 0.92) but greatly compromised ASW-beta (0.0045), suggesting a more aggressive over-correction. Similarly, LIGER performed the best in kBET (0.5978) but conferred a much lower ARI (0.8476) and low ASW-beta (0.0912), indicating a more severe overcorrection of the batch effects (i.e., mixing beta cells with other cells).

We then visualized the clustering by UMAP to examine how the beta cells are assigned to different clusters (**Supp. Fig. S9**). We found that beta cells (colored in red) were clustered separately by scETM from other cell types. In contrast, beta cells were mixed up with other cell types by methods including Harmony and LIGER, which overcorrected the batch effects when integrating the five batches. Visually, we also observe that scETM+adv method moves the beta cell cluster closer to the alpha cell cluster, confirming a higher level of overcorrection compared to scETM.

The MR dataset is a collection of two independent studies on mouse retina [36, 37]. Here we consider the two source studies as two batches, hereafter referred to as the Macosko batch and the Shekhar batch. Many cell types are uniquely present in the Macosko batch (**Supp. Table S6**). There is also a large difference in the cell proportion between the two batches. In particular, rods is only 0.35% in Shekhar but 65% in Macosko. In this scenario, we expect that methods that overcorrect the batch effects would tend to mix rods with cells of other cell types from the Shekhar batch, resulting in low ARI and high kBET. On the contrary, a desirable integration method would strike a balance between the ARI (or ASW) and kBET on this combined dataset. Therefore, this setup imposes a great challenge on the integration methods.

Overall, scETM achieved the highest ARI (0.859), reasonable ASW (0.2873), and modest kBET (0.0656), indicating its ability to capture the true biology from the data without over-correcting the batch effects (**Supp. Table S7**). In contrast, LIGER is more aggressive in its batch correction, resulting in the highest kBET score of 0.176 but lowest ARI score of 0.714. We further investigated the extent of improving kBET while maintaining a high ARI score with scETM+adv. Indeed, scETM+adv conferred an increased kBET of 0.1410 and a reasonably high ARI score of 0.7720. Visualizing the clustering of each method using UMAP (**Fig. 2**) confirms the quantitative clustering results.

Incidentally, we also notice that scETM is not sensitive to the 669 doublelets or contaminants, all of which were from the Shekhar batch (**Supp. Table S8**). In contrast, if we did not filter out the doublets/contaminants, the performance of LIGER and Seurat degrades drastically possibly due to batch over-correction or failing to integrate the same cell types from different batches together.

### Scalability

A key advantage of scETM is its high scalability and efficiency. We demonstrated this by comparing the run time, memory usage, and clustering performance of the state-of-the-art models using their recommended pipelines when integrating a merged dataset consisting of cells from the MDD and AD datasets (**Methods**). Because of the simple model design and efficient implementation (e.g., sparse matrix representation, multi-threaded data retrieval, etc; **Discussion**), scETM achieved the shortest run time among all deep-learning based models (**Fig. 3a**). Specifically, on the largest dataset (148,247 cells), scETM ran 3-4 times faster than scVI and scVI-LD, and over 10 times faster than scVAE-GM. We note that the run time largely depends on the implementation rather than the network architectures and loss functions in these deep learning methods. Harmony and Scanorama were the only methods faster than scETM, yet they both operate on a hundred principal components at most. Although for comparison purpose we used the top 3000 most variable genes for all of the methods, scETM can easily scale to all of the genes, which is more desirable because the resulting model can generalize to other datasets.

**Figure 3:**
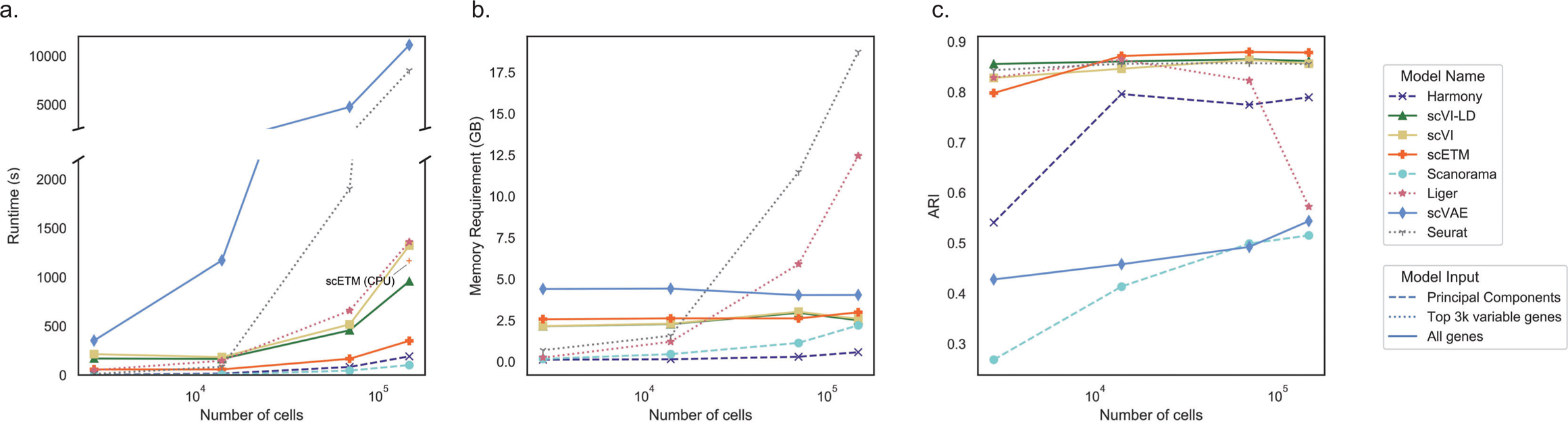
Benchmark of the efficiency and scalability of the seven scRNA-seq clustering algorithms. The line styles in the plot indicate model inputs. The number of genes was fixed to 3000 in this experiment. We increased the number of cells randomly sampled from the combined AD and MDD dataset and evaluated the performance of each method in terms of: **(a)** runtime, **(b)** memory usage, and **(c)** Adjusted Rand Index (ARI). The run time of scETM on CPU is annotated on the left panel.

Because of the amortized stochastic variational inference [32, 41, 42], scETM in principle takes linear run-time and constant memory with respect to the sample size per training epoch. The use of multi-threaded data loader to streamline the random minibatch retrieval and loading further speed up the training process in practice. In contrast, the memory requirement of Seurat increases rapidly with the number of cells, due to the vast numbers of plausible anchor cell pairs in the two brain datasets (**Fig. 3b**). In terms of clustering accuracy, scETM consistently confers competitive performance, whereas Harmony and Scanorama perform unstably as dataset sizes vary (**Fig. 3c**). UMAP visual inspection of scVAE embeddings suggests that scVAE likely suffers from under-correction of batch effects (**Supp. Fig. S8**). The sudden drop of LIGER’s clustering performance in the largest benchmark dataset may be due to batch overcorrection.

Although it has been widely accepted by the deep learning community that computing using Graphical Processing Units (GPUs) results in *∼*10*×* speedup over computing using CPU, the adoption of GPUs in the computational biology community is beginning to catch up. In our non-exhaustive experiment on the Mouse Pancreas dataset, training scETM for 1000 steps on the 6-core Core i7 10750H CPU requires 650 seconds (**Fig. 3a**), while with an Nvidia RTX 2070 laptop GPU it only takes 50 seconds - a 13*×* speedup over the CPU computer.

### Transfer learning across single-cell datasets

A prominent feature of scETM is that its parameters, hence the knowledge of modeling scRNA-seq data, are transferable across datasets. Specifically, as part of the scETM, the encoder trained on a reference scRNA-seq dataset can be applied to infer cell topic mixture of a target scRNA-seq dataset (**Fig. 1b**), regardless of whether the two datasets share the same cell types. As an example, we trained an scETM model on the *Tabula Muris* FACS dataset (TM (FACS)) (which is a subset of the TM dataset) from a multi-organ mouse single-cell atlas, and evaluated it using the MP data, which only contains mouse pancreatic islet cells. Although the two datasets were obtained using different sequencing technologies in two independent studies, the model yielded an encouragingly high ARI score of 0.941, considering that a model directly trained on MP achieves ARI 0.946. Interestingly, in the UMAP plot, the TM (FACS)- pretrained model placed B cells, T cells and macrophages far away from other clusters and separated B cells and T cells from macrophages, which is not observed in the model directly trained on MP (**Supp. Fig. S3,S4,S5,S6,S7**; **Supp. Table S9**). We repeated the same experiment 3 times with different random seeds and observed consistently that B and T cells are close to each other and distant from macrophages (**Supp. Fig. S10**). We also experimented transfer learning by first training scETM on TM (FACS) *with pancreas removed* and then applied to MP dataset. The performance decreased but is still reasonably good (**Supp. Table S9**), demontrasting scETM’s ability to transfer knowledge across tissues.

Encouraged by the above results, we then performed a comprehensive set of cross-tissue and cross-species transfer learning analysis with 6 tasks (**Methods**): (1) Transfer between the TM (FACS) and the MP dataset (including MP*→*TM (FACS)); (2) Transfer between the Human Pancreas (HP) dataset and the Mouse Pancreas (MP) dataset; (3) Transfer between the Human primary motor cortex (M1C) (HumM1C) dataset and the Mouse primary motor area (MusMOp) dataset both obtained from the Allen Brain Map data portal [43]. We chose to transfer between the human M1C and mouse MOp because of the high number of shared cell types between the brain regions of the two species. The batches for HumM1C are the two post-mortem human brain M1 specimens and the two mice for MusMOp. Note that in these transfer learning tasks (*A → B*) we only corrected batch effects during the training on the source data *A* but not during the transfer to the target data *B*.

As a comparison, we evaluated and visualized the clustering results in all 6 transfer learning tasks using scETM, scVI-LD, and scVI (**Fig. 4**; **Supp. Table S10** and S11). Overall, scETM achieved the highest ARI across all tasks and competitive kBET scores. In particular, scETM trained on TM (FACS) on heterogeneous tissues clustered much better the MP cells (ARI: 0.941; kBET: 0.339) than scVI (ARI: 0.484; kBET: 0.257) and scVI-LD (ARI: 0.398; kBET: 0.256). Remarkably, scETM trained only on the MP dataset can cluster reasonably well the much larger TM single-cell data, which were collected from diverse primary tissues including pancreas. This implies that scETM does not merely learn cell-type-specific signatures but also the underlying transcriptional programs that are generalizable to unseen tissues.

**Figure 4:**
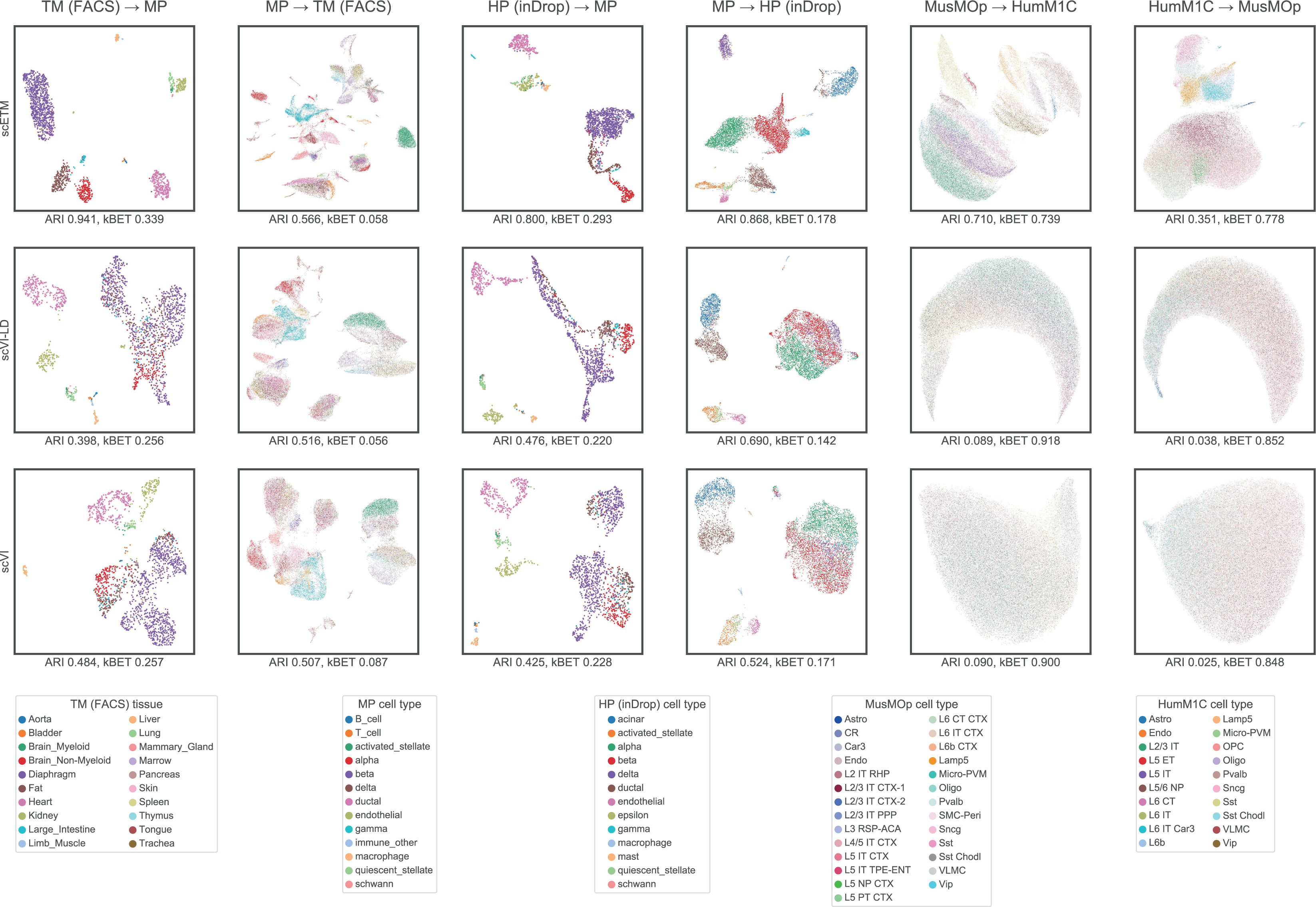
Cross-tissue and cross-species zero-shot transfer learning. Each panel displays the UMAP visualization of cells by training on dataset A and the applied to dataset B (i.e., A B). In total, we performed 6 transfer learning tasks: TM (FACS) MP, HP (inDrop) MP, MusMOp HumM1C. For each task, we evaluated scETM, scVI-LD, or scVI, which are the rows in the above figure. The cells are colored by tissues or cell types as indicated in the legend. The corresponding ARI and kBET are indicated below each panel. For MP TM (FACS), we evaluated the ARI based on the 92 cell types although we colored the cells by the 20 tissues of origin instead of their cell types because of the large number of cell types. Abbreviations: TM (FACS): Tabula Muris sequenced with Fluorescence-Activated Single Cell Sorting; MP: Mouse Pancreas; HP (inDrop): Human Pancreas sequenced with InDrop technology; HumM1C: human primary motor cortex (from Allen Brain map); MusMOp: Mouse primary motor area (from Allen Brain map).

In cross-species transfer learning between HP and MP, scETM captured better the conserved pancreas functions compared to scVI and scVI-LD (**Fig. 4**; **Supp. Table S10**). On the other hand, cross-species transfer between MusMOp and HumM1C is a much more challenging task due to the evolutionarily divergent functions of the brains between the two species.

Nonetheless, scETM conferred a much higher ARI of 0.696 for the MusMOp*→*HumM1C transfer and ARI of 0.167 for the HumM1C*→*MusMOp transfer. In contrast, scVI-LD and scVI did not work well on these tasks with ARI scores lower than 0.1. Since they cannot separate cells by cell types, all cells are mixed together, leading to a high kBET score. The improvements achieved by scETM over scVI(-LD) are possibly attributable to the simpler linear batch correction on the source data, jointly learning topic and gene embedding, and the topic modeling formalism, which together lead to an encoder network that is better at capturing the transferable cellular programs.

### Pathway enrichment analysis of scETM topics

We next investigated whether the scETM-inferred topics are biologically relevant in terms of known gene pathways in human. One approach would be to arbitrarily choose a number of top genes under each topic and test for pathway enrichment using hypergeometric tests. This approach works well when there are asymptotic p-values at the individual gene level. In our case, each gene is characterized by the topic scores, making it difficult to systematically choose the number of top genes per topic. To this end, we resorted to Gene Set Enrichment Analysis (GSEA) [44]. Briefly, we calculated the maximum running sum of the enrichment scores with respect to a query gene set by going down the gene list that is sorted in the decreasing order by a given topic distribution **β***_k_* (**Methods**). For each dataset, we trained a scETM with 100 topics.

For the HP dataset, each topic detected many significantly enriched pathways with Benjamini-Hochberg False Discover Rate (FDR) *<* 0.01 (**Fig. 5a**). Many of them are relevant to pancreas functions, including insulin processing (**Fig. 5b**), insulin receptor recycling, insulin glucose pathway, pancreatic cancer, etc (**Supp. Table S12**). Because scETM jointly learns both the gene embeddings and topic embeddings, we can visualize both the genes and topics in the same embedding space via UMAP (**Fig. 5c**). Indeed, we observe a strong co-localization of the genes in Insulin Processing pathway and the corresponding enriched topic (i.e., Topic 54).

**Figure 5:**
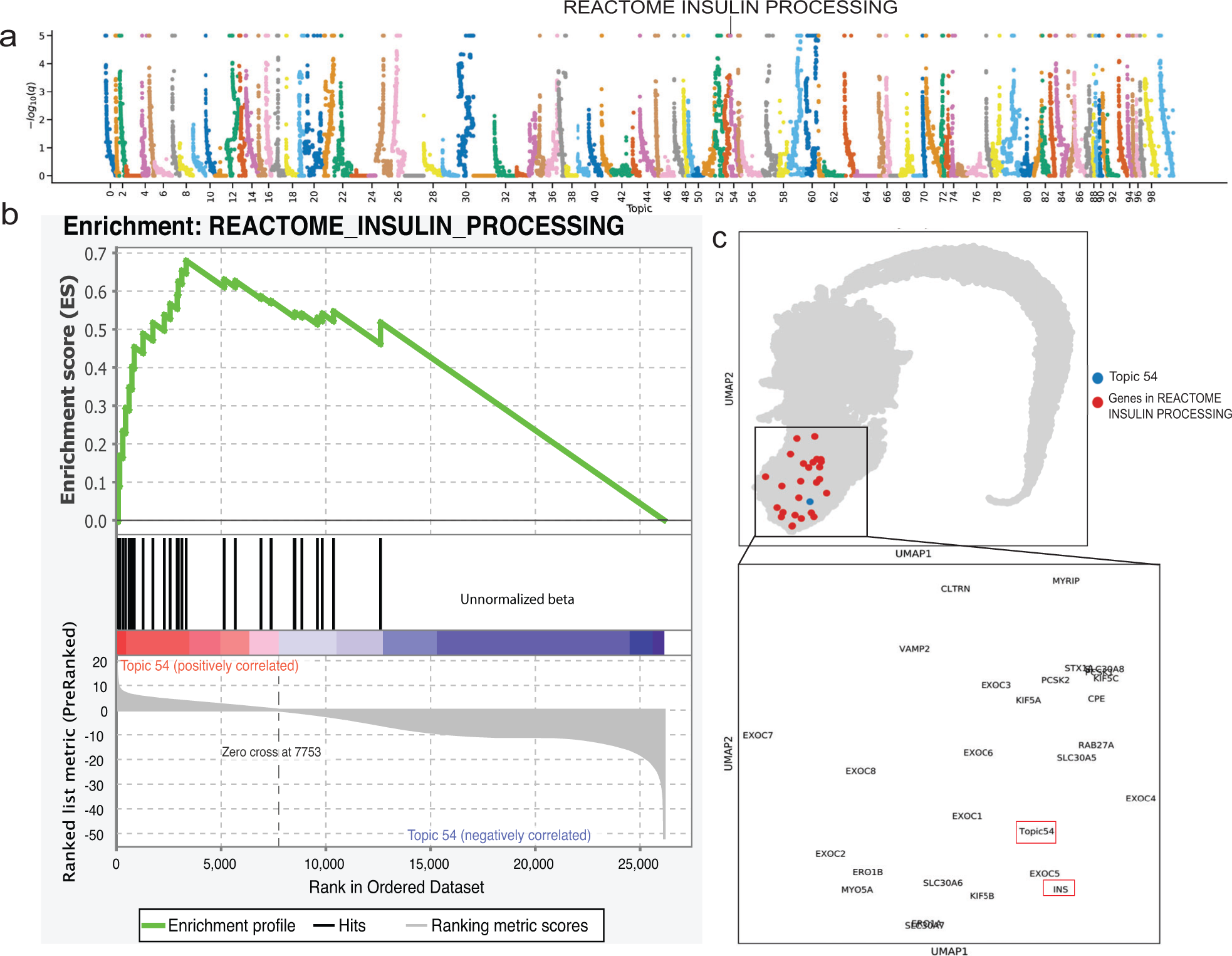
Gene set enrichment analysis of the Human Pancreas dataset. **(a)** Manhattan plot of the GSEA results on the 100 scETM topics learned from the HP dataset. The x-axis is the topic and the y-axis is the -log q-value from the permutation test corrected for multiple testings by the BH-method. The dots correspond to the tested GSEA gene sets. The maximum -log q-value is capped at 5. **(b)** Leading edge analysis of Insulin Processing (IP) pathway using Topic 54. The genes are ranked by the topic score (i.e., the unnormalized topic mixture beta) under Topic 54. The running sum enrichment score was calculated by GSEA. The black bars in the middle indicate the genes that are found in the IP pathway. Topic 54 is significantly enriched in Insulin Processing pathway (GSEA permutation test BH-adjusted q-value = 0). **(c)** UMAP visualization of the gene and topic embeddings learned from the HP dataset. Genes in IP are colored in red and Topic 54 in blue. The inset box displays a magnified view of the cluster zooming into the IP pathway genes (including *Insulin* (*INS*)) that are near Topic 54 in the embedding space.

For the AD dataset, we found topics enriched for Reactome Amyloid Fiber Formation, KEGG AD, and Deregulated CDK5 triggers multiple neurodegenerative in AD (**Supp. Fig. S11**; **Supp. Table S13**). For MDD dataset, we found enrichment for Substance/Drug Induced Depressive Disorder (**Supp. Fig. S12**, S13; **Supp. Table S14**). The full GSEA enrichment results for all 3 datasets are listed in **Supp. Table S18**. As a comparison, we also performed GSEA over the 100 gene loadings learned by scVI-LD (matching the 100 topics in the scETM) on these 3 datasets but found fewer relevant distinct gene sets (**Supp. Table S12**,S13,**Supp. Table S14**) or weaker statistical enrichments by GSEA (**Supp. Fig. S14**).

### Differential scETM topics in disease conditions and cell types

We sought to discover scETM topics that are condition-specific or cell-type specific. Starting with the AD dataset, we found that the scETM-learned topics are highly selective of cell-type marker genes (**Fig. 6a**) and highly discriminative of cell types (**Fig. 6b**). To detect disease signatures, we separated the cells into the ones derived from the 24 AD subjects and the ones from the 24 control subjects. We then performed permutation tests to evaluate whether the two cell groups exhibit significant differences in terms of their topic expression (**Methods**). Topic 12 and 58 are differentially expressed in the AD cells and control cells (**Fig. 6c**, d; permutation test p-value = 1e-5). Interestingly, topic 58 is also highly enriched for mitochondrial genes. Indeed, it is known that *β*-amyloids selectively build up in the mitochondria in the cells of AD-affected brains [45]. For the MDD dataset, topics 1, 52, 68, 70, 86 exhibit differential expressions between the suicidal group and the healthy group (**Supp. Fig. S18c**) and interesting neurological pathway enrichments (**Supp. Table S14**).

**Figure 6:**
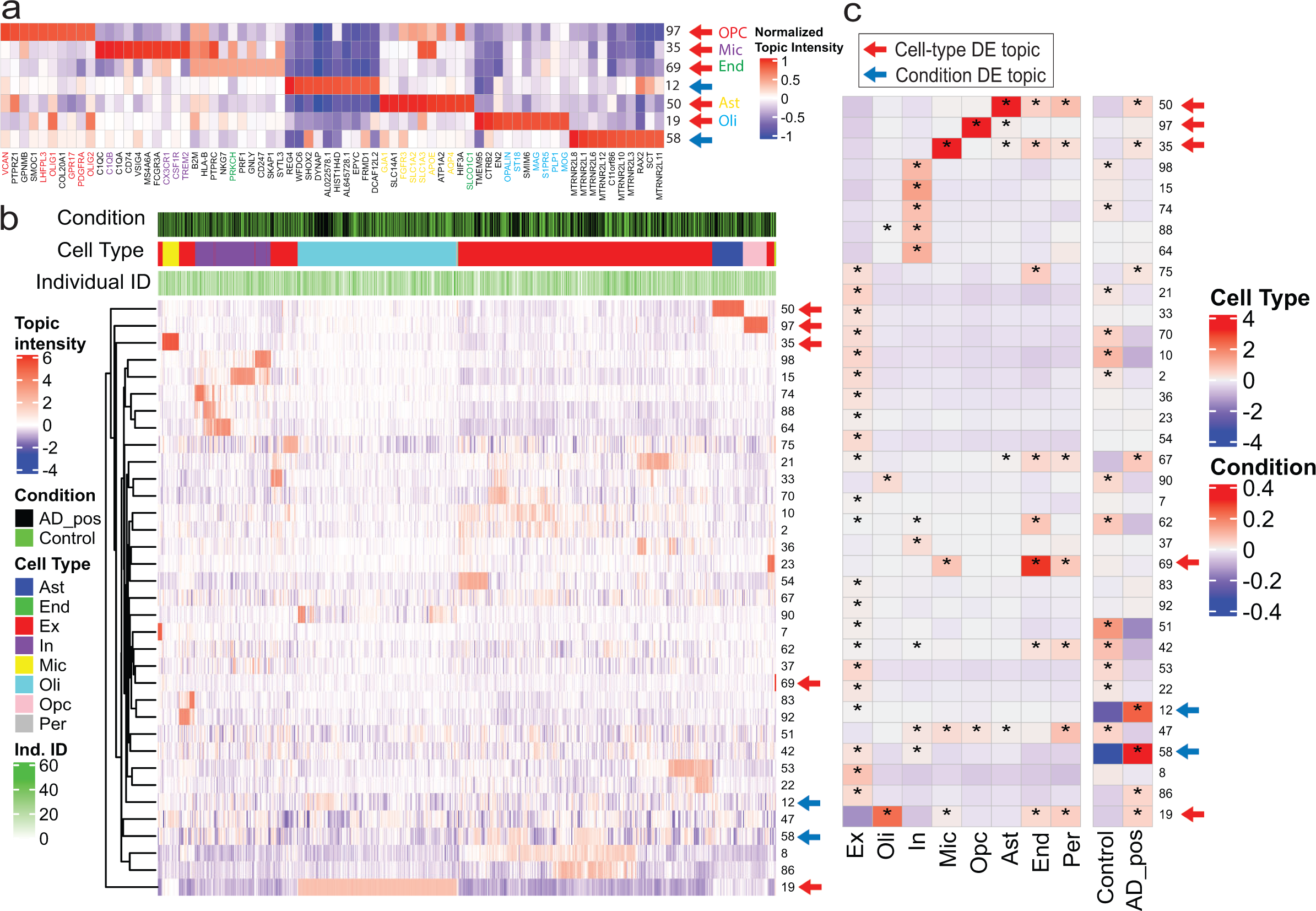
scETM topic embeddings of the Alzheimer’s Disease snRNA-seq dataset. **(a)** Gene-topic heatmap. The top genes which are known as cell-type marker genes based on PanglaoDB are highlighted. For visualization purposes, we divided the topic values by the maximum absolute value within the same topic. Only the differential topics with respect to cell-type or AD were shown. **(b)** Topics intensity of cells (n=10,000) sub-sampled from the AD dataset. Topic intensities shown here are the Gaussian mean before applying softmax. Only the select topics with the sum of absolute values greater than 1500 across all sampled cells are shown. The three color bars show disease conditions, cell types, and batch identifiers (i.e., subject IDs). **(c)** Differential expression analysis of topics across the 8 cell types and 2 clinical conditions. Colors indicate mean differences between cell groups with and without a certain label (cell-type or condition). Asterisks indicate Bonferroni-corrected empirical p-value *<* 0.05 for 100,000 permutation tests of up-regulated topics in each cell-type and disease labels.

We also identified several cell-type-specific scETM topics from the HP, AD, and MDD datasets. In HP, topics or metagenes 20, 45, 99 are up-regulated in acinar cells, topic 12 up-regulated in macrophage, topic 52 up-regulated in delta, and topics 30 and 37 are up-regulated in more than one cell types, including endothelial, stellate and others (**Supp. Fig. S15**). In AD, as shown by both the cell topic mixture heatmap and the differential expression analysis (**Fig. 6b**), topics 19, 35, 50, 69, 97 are up-regulated in oligodendrocytes, micro/macroglia, astrocytes, endothelial cells, and oligodendrocyte progenitor cells (OPCs) respectively (permutation test p-value = 1e-5; **Fig. 6b**, **Supp. Fig. S16**). Interestingly, two subpopulations of cells from the oligodendrocytes (Oli) and excitatory (Ex) exhibit high expression of topics 12 and 58, respectively, and are primarily AD cells (**Supp. Fig. S17**). Among them, there is also a strong enrichment for the female subjects, which is consistent with the original finding [35].

For MDD, topics 1, 20, 59, 64 and 72 are up-regulated in astrocytes, oligodendrocytes, micro/macroglia, endothelial cells, and OPCs, respectively (**Supp. Fig. S18c**). This is consistent with the heatmap pattern (**Supp. Fig. S18b**). Several topics are dominated by long non-coding RNAs (lincRNAs) (**Fig. S18a**). While previous studies have suggested that lincRNAs can be cell-type-specific [46], it remains difficult to interpret them [47]. We further experimented the enrichment using only the protein coding genes, but did not find significantly more marker genes among the top 10 genes per topic (**Supp. Fig. S19**).

### Pathway-informed scETM topics

To further improve topic interpretability, we incorporated the known pathway information to guide the learning of the topic embeddings (**Fig. 7a**). We denoted this scETM variant as the pathway-informed scETM or *p-scETM*. In particular, we fixed the gene embedding **ρ** to a pathways-by-genes matrix obtained from pathDIP4 database [48, 49]. We then learned only the topics embedding **α**, which provides direct associations between topics and pathways (**Methods**).

**Figure 7:**
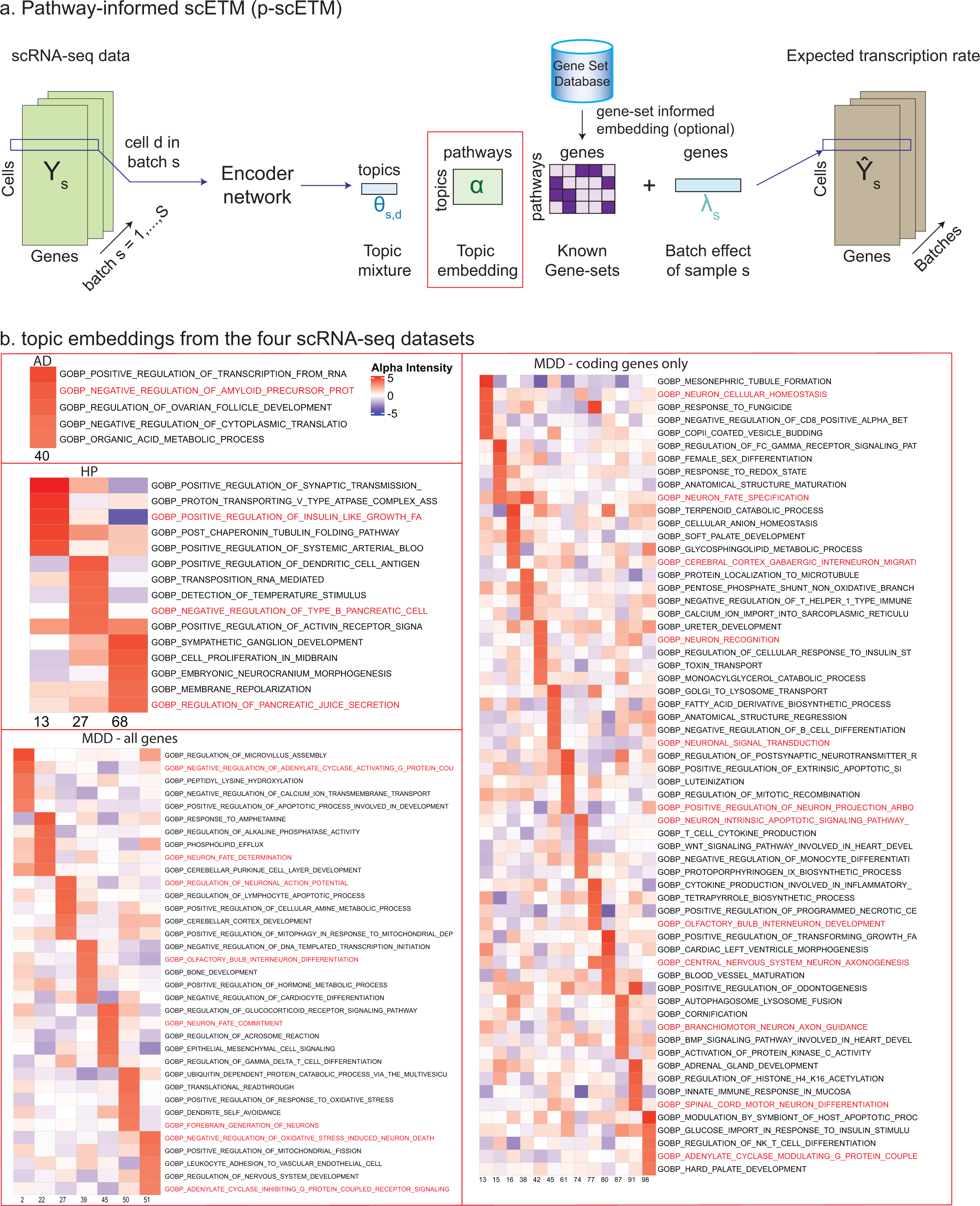
Pathway-topics embeddings learned by the pathway-informed scETM (p-scETM). **(a)** p-scETM overview. Pathways information as pathways-by-genes are provided as the gene embedding in the linear decoder. The learned topic embedding is the direct association between the topics and pathways. **(b)** The pathway-topics heatmap of top 5 pathways in selected topics. Here the pathways are the Gene Ontology - Biological Processes (GO-BP) terms. For the HP dataset, GO-BP terms whose names include the keywords “insulin” or “pancreatic” were highlighted. For the AD dataset, GO-BP terms whose names include the keywords “amyloid” or “alzheimer” were highlighted. For *MDD - all genes* and *MDD - coding genes only*, GO-BP terms whose names include the keywords “neuron” or “G-protein” were highlighted.

We tested p-scETM on the HP, AD and MDD datasets. Without compromising the clustering performance (**Supp. Table S15**), p-scETM learned functionally meaningful topic embeddings (**Fig. S20**; **Supp. Table S16**,S17). In the HP topic embeddings, we found Insulin Signaling, Nutrient Digestion and Metabolism to be the top pathways among several topics (**Fig. S20a**). In the MDD topic embeddings, the top pathway associated with Topic 40, Beta-2 Adrenergic Receptor Signaling, was also enriched in a MDD genome-wide association studies [50]. In the AD topic embeddings, we found the association between Topic 9 and Alzheimer Disease-Amyloid Secretase pathway.

To further demonstrate the utility of p-scETM, we also used 7481 Gene Ontology Biological Process (GO-BP) terms [51, 52] as the fixed gene embedding, which learns the topics-by-GOs topic embedding from each dataset. Under each topic, we selected the top 5 high-scoring GO- BP terms to examine their relevance to the target tissue or disease (**Fig. 7b**; **Supp. Table S19**). For the HP dataset, Negative Regulation of Type B Pancreatic Cell (GO:2000675) and Regulation of Pancreatic Juice Secretion (GO:0090186) are among the top GO-BP terms for Topics 27 and 68, respectively. For the AD dataset, Amyloid Precursor Protein Biosynthetic Process

(GO:0042983) is among the top 5 GO-BP terms under Topic 40. For the MDD dataset, similar top GO-BP terms were found among topics learned using all of the genes and using only the coding gene. Many topics exhibit high embedding scores for neuronal functions including Neuronal Signal Transduction (GO:0023041), Central Nervous System Projection Neuron Axonogenesis (GO:0021952), and Branchiomotor Neuron Axon Guidance (GO:0021785). Interestingly, Topic 98 in *MDD - coding genes only* and Topics 22, 51 in *MDD - all genes* involve Adenylate Cyclase Modulating G-protein Coupled Receptors (GPCRs) Signaling (GO:0007188), which is the target of several recently-developed antidepressant drugs [53].

## Discussion

As scRNA-seq technologies become increasingly affordable and accessible, large-scale datasets have emerged. This challenges traditional statistical approaches and calls for transferable, scalable, and interpretable representation learning methods to mine the latent biological knowledge from the vast amount of scRNA-seq data. To address these challenges, we developed scETM and demonstrated its state-of-the-art performance on several unsupervised learning tasks across diverse scRNA-seq datasets. scETM demonstrates excellent capabilities of batch effect correction and knowledge transfer across datasets.

In terms of integrating multiple scRNA-seq data from different technologies, experimental batches, or studies, we introduce a simple batch-effect bias term to correct for non-biological effects. This in general improves the cell clustering and topic quality. When using the original ETM [33], we observed that ubiquitously expressed genes such as *MALAT1* tended to appear among the top genes in several topics. Our scETM corrects the background gene expressions by the gene-dependent and batch-dependent intercepts. As a result, the ubiquitously expressed genes do not dominate all topics from scETM. We also introduced a more aggressive batch correction strategy by adversarial network loss, which shows improved kBET with small trade-off for the ARI in most datasets.

In terms of scalability, although scETM is similar to other existing VAE models in terms of theoretical time and space complexity, we emphasize that implementation is also very important, especially for deep learning models. For example, scVAE-GM [11] is much slower and more memory consuming than scVI [6], while they are very similar VAE models. One of the main speedups provided by scETM comes from our implementation of a multi-threaded data loader for minibatches of cells, which does not need to be re-initialized at every training epoch as the standard PyTorch DataLoader. Compared to scVI and scVI-LD, the normalized counts in both the encoder input and the reconstruction loss used by scETM remove the need to infer the cell-specific library size variable, and the simpler categorical likelihood choice also helps reduce the computational time.

In terms of transferability, many existing integration methods require running on both reference and query datasets to perform post hoc analyses such as joint clustering and label transfer [7, 9, 10, 28]. In contrast, our method enables a direct or zero-shot knowledge transfer of the pretrained encoder network parameters learned from a reference dataset in annotating a new target dataset without further training. We demonstrated this important aspect in cross-technology, cross-tissue, and cross-species applications, for which we achieved superior performance compared to the state-of-the-art methods.

In terms of interpretability, our quantitative experiments showed that scETM identified more relevant pathways than scVI-LD. Qualitative experiments also show that scETM topics preserve cell functional and cell-type-specific biological signals implicated in the single-cell transcriptomes. By seamlessly incorporating the known pathway information in the gene embedding, p-s cETM finds biologically and pathologically important pathways by directly learning the association between the topics with the pathways via the topic embedding. Recently proposed by [54], single-cell Hierarchical Poisson Factor (scHPF) model applies hierarchical Poisson factorization to discover interpretable gene expression signatures in an attempt to address the interpretability challenge. However, compared to our model, scHPF lacks the flexibility in learning the gene embedding and incorporating existing pathway knowledge, and is not designed to account for batch effects. Moreover, scETM has the benefits of both flexibility in the neural network encoder and the interpretability in the linear decoder.

As future work, we will extend scETM in several directions. To further improve batch correction, as our current model only considers a single categorical batch variable, we can extend it to correct for multiple categorical batch variables. For a small number of categorical batch variables, we may use several sets of batch intercept terms to model them. For hierarchical batch variables, we may use a tree of batch intercept terms. For numerical batch effects such as subject age, one way is to convert them into categorical batch variables by numerical ranges. When the number of batch variables become larger, we consider three strategies. First, we can add the batch variables as the covariates in the linear regression on the gene expression and fit the linear coefficients each corresponding to a sample-dependent batch variable. Second, we can factorize the batches-by-genes into batches-by-factors and factors-by-genes. Learning the two matrices will be similar to the scETM algorithm. Third, we can extend our current scETM+adv to correct for both categorical and continuous batch variables with a discriminator network, which predicts batch effects using the encoder-generated cell topic mixture [38].

To further improve data integration, we can extend scETM to a multi-omic integration method, which can integrate scRNA-seq plus other omics such as protein expression measured in the same cells as scRNA-seq [55] or scATAC-seq measured in different cells but the same biological system [7]. In these applications, multi-modality over different omics will need to be considered to capture the intrinsic technical and biological variance of each omic while borrowing information among them.

To further improve interpretability, the original ETM used pretrained word embedding from word2vec [56] on a larger reference corpus such as Wikipedia to improve topic quality on modeling the target documents [33]. Similarly, although we demonstrated the use of existing pathway information in p-scETM, we can also pretrain our gene embeddings on PubMed articles, gene regulatory network, protein-protein interactions, or Gene Ontology graph using either gene2vec [57] or more general graph embedding approaches [58, 59] [59]. We expect that the gene embedding pretrained from these (structured) knowledge graphs will further improve the efficiency and interpretability of scETM.

Together, scETM serves as a unified and highly scalable framework for integrative analysis of large-scale single-cell transcriptomes across multiple datasets. Compared to existing methods, scETM offers consistently competitive performance in data integration, transfer learning, scalability, and interpretability. The simple Bayesian model design in scETM also provides a highly expandable framework for future developments.

## Methods

### scETM data generative process

To model scRNA-seq data distribution, we take a topic-modeling approach [60]. In our framework, each cell is considered as a “document”, each scRNA-seq read (or UMI) as a “token” in the document, and the gene that gives rise to the read (or UMI) is considered as a “word” from the vocabulary of size *V* . We as ed as a mixture of latent cell types, which are commonly referred to as the latent topics. The original LDA model [60] defines a fixed set of *K* independent Dirichlet distributions ***β*** over a vocabulary of size *V* . Following the ETM model [33], here we decompose the unnormalized topic distribution 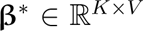 into the topic embedding 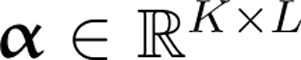 and gene embedding 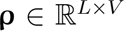, where *L* denotes the size of the embedding space. Therefore, the unnormalized probability of a gene belonging to a topic is proportional to the dot product between the topic embedding matrix and the gene embedding matrix. Formally, the data generating process of each scRNA-seq profile *d* is:

1. Draw a latent cell type proportion **θ***_d_* for a cell *d* from logistic normal **θ***_d_ ∼ LN* (0, **I**): 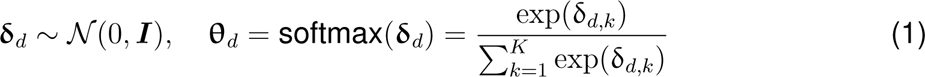
2. For each gene read (or UMI) *w_i,d_* in cell *d*, draw gene *g* from a categorical distribution cat(r_d_): 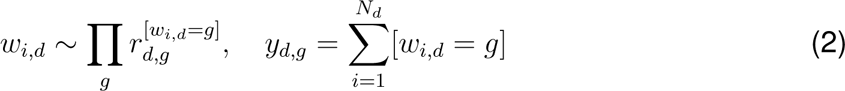 Here *N_d_* is the library size of cell *d*, *w_i,d_* is the index of the gene that gives rise to the *i^th^* read (or UMI) in cell *d* (i.e., [*w_i,d_* = *g*]), and *y_d,g_* is the total counts of gene *g* in cell *d*. The transcription rate *r_d,g_* is parameterized as follows: 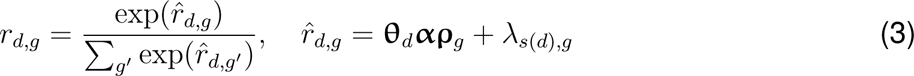 Here **θ***_d_* is the 1 *× K* cell topic mixture for cell *d*, **α** is the global *K × L* cell topic embedding, **ρ***_g_* is a *L ×* 1 gene-specific embedding, and *λ_s_*_(_*_d_*_)_*_,g_* is the batch-dependent and gene-specific scalar effect, where *s*(*d*) indicates the batch index for cell *d*. Notably, to model the sparsity of gene expression in each cell (i.e., only a small fraction of the genes have non-zero expression), we use the softmax function to normalize the transcription rate over all of the genes.

### scETM model inference

In scETM, we treat the latent cell topic mixture **δ***_d_* for each cell *d* as the only latent variable. We treat the topic embedding **α**, the gene-specific transcriptomic embedding **ρ**, and the batch-effect **λ** as point estimates. Let **Y** be the *D × V* gene expression matrix for *D* cells and *V* genes. The posterior distribution of the latent variables *p*(**δ***|***Y**) is intractable. Hence, we took a variational inference approach using a proposed distribution *q*(**δ***_d_*) to approximate the true posterior. Specifically, we define the following proposed distribution: *q*(**δ** *|* **y**) = *_d_ q*(δ*_d_|***y***_d_*), where *q*(**δ***_d_|***y***_d_*) = **µ***_d_* + diag(**σ***_d_*)*N* (0*, I*) and [**µ***_d_,* log **σ**^2^] = NNET(**y***_d_*; **W***_θ_*). Here **y**Y*_d_* is the normalized counts for each gene as the raw count of the gene divided by the total counts in cell *d*. The function NNET(**v**; **W**) is a two-layer feed-forward neural network used to estimate the sufficient statistics of the proposed distribution for the cell topic mixture **δ***_d_*.

To learn the above variational parameters **W***_θ_*, we optimize the evidence lower bound (ELBO) of the log likelihood, which is equivalent to minimizing the Kullback-Leibler (KL) divergence between the true posterior and the proposed distribution: ELBO = E*_q_* [log *p*(**Y***|*Θ)]*−*KL [*q*(Θ*|***Y**)*||p*(Θ)].

The Bayesian learning is carried out by maximizing the reconstruction likelihood with regularization in the form of KL divergence of the proposed distribution (*q*(**δ***_d_|***y***_d_*) = *N* (**µ***_d_,* diag(**σ***_d_*))) from the prior (*p*(**δ***_d_*) = *N* (0, **I**)). For computational efficiency, we optimize ELBO with respect to the variational parameters by amortized variational inference [32, 41, 42]. Specifically, we draw a sample of the latent variables from *q*(**δ** *|* **y**) for a minibatch of cells from reparameterized Gaussian proposed distribution *q*(**δ** *|* **y**) [32], which has the mean and variance determined by the NNET functions. We then use those draws as the noisy estimates of the variational expectation for the ELBO. The optimization is then carried out by back-propagating the gradients into the encoder weights and the topic and gene embeddings.

### Details of training scETM

We chose the encoder for inferring the cell topic mixture to be a 2-layer neural network, with hidden sizes of 128, ReLU activations [61], 1D batch normalization [62], and 0.1 dropout rate between layers. We set the gene embedding dimension to 400, and the number of topics to 50. We optimize our model with Adam Optimizer and a 0.005 learning rate. To prevent over-regularization, we start with zero weight penalty on the KL divergence and linearly increase the weight of the KL divergence in the ELBO loss to 10*^−^*^7^ during the first <u>^1^</u> epochs. With a minibatch size of 2000, scETM typically needs 5k-20k training steps to converge. We show that our model is robust to changes in the above hyperparameters (**Supp. Table** S2). During the evaluation, we used the variational mean of the unnormalized topic mixture **µ***_d_* in *q*(**δ***_d_|***y***_d_*) = *N* (**µ***_d_,* diag(**σ***_d_*)) as the scETM cell topic mixture for cell *d*.

### scETM+adv: adversarial loss for further batch correction

In the scETM+adv variant, we added a discriminator network (a two-layer fully-connected network) to predict batch labels using the unnormalized cell topic mixture embedding **δ** generated by the encoder network. This discriminator helps batch correction in an adversarial fashion.

Specifically, in each training iteration, we first update the scETM parameters once by maximizing the ELBO plus the batch prediction cross-entropy loss from the discriminator, with a hyperparameter controlling the weight of the latter term. It should be noted that by maximizing the prediction loss the encoder network learns to produce batch agnostic cell embeddings. Then we update the discriminator network eight times by trying to minimize the cross-entropy loss in predicting the batch labels.

### scETM software

We implemented scETM using the PyTorch library [63]. Our initial implementation was based on the ETM code from GitHub (adjidieng/ETM) by [33]. Since then, we completely revamped the code to substantially improve the scalability and to integrate it into the Python ecosystem. In particular, we packaged and released our code on PyPI so one can easily install the package by entering pip install scETM in the terminal. The package is integrated with scanpy [16] and tensorboard [64]. Users can view the cell, gene and topic embeddings interactively via tensorboard. For example, one can easily train a scETM as follows: from scETM import scETM, UnsupervisedTrainer model = scETM(adata.n_vars, adata.obs.batch_indices.nunique()) trainer = UnsupervisedTrainer(model, adata) trainer.train(save_model_ckpt = False) model.get_all_embeddings_and_nll(adata)

The above code snippet will instantiate an scETM model object, train the model, infer the unnormalized cell topics mixture of adata and store them in adata.obsm[‘delta’]. We can also access the gene and topic embeddings via adata.varm[‘rho’] and adata.uns[‘alpha’].

### Transfer learning with scETM

When transferring from a reference dataset to a target dataset, we operate on the genes common to both datasets. For cross-species transfer, the orthologous genes based on the Mouse Genome Informatics database [65, 66] are considered as common genes. The trained scETM encoder can be directly applied to an unseen target dataset, as long as the genes in the target dataset are aligned to the genes in the reference dataset. In the main text, for example, we trained scETM on a reference dataset and evaluated the scETM-encoder on a target dataset in 6 transfer learning tasks (**Fig. 4**).

### Pathway enrichment analysis

To assess whether a topic is *enriched* in any known pathway, one common way is to test for Over Representation Analysis (ORA) [67]. However, ORA requires choosing a subset of genes (e.g., from differential expression analysis). While we could choose the top genes scored by each topic, it requires some arbitrary threshold to select those genes. To avoid thresholding genes, we employed Gene Set Enrichment Analysis (GSEA) [44]. GSEA calculates a running sum of enrichment scores (ES) by going down the gene list that is sorted in the decreasing order by their association statistic with a phenotype.

In our context, we treated the gene scores under each topic from the genes-by-topics matrix (i.e., **β**) as the association statistic. The ES for a gene set S is the maximum difference between P_hit(S,i) and P_miss(S,i), where P_hit(S,i) is the fraction of genes in S weighted by their topic scores up to gene index i in the sorted list and P_miss(S,i) is the fraction of genes *not in S* weighted by their topic scores up to gene index i in the sorted list. The enrichment p-value for each gene set is computed by permutation tests by randomly shuffling the gene symbols on the sorted list (while keeping the gene topic scores in the decreasing order) 1000 times to compute the null distribution of the ES for each gene set and each topic. The empirical p-value was calculated as (N’+1)/(N+1), where N’ is the number of permutation trials in which ES is greater than the observed ES, and N is the total number of trials (i.e., 1000). We then corrected the p-values for multiple testing using Benjamini-Hochberg (BH) method [68].

For AD and HP datasets, we used the MSigDB Canonical Pathways gene sets [69] as the gene set database in GSEA; and for MDD, we used PsyGeNET database [70] in order to find psychiatric disease-specific associations. We also run GSEA for scVI-LD gene loadings for comparison. The detailed pathway enrichment statistic can be found in **Supp. Table S18**.

### Differential analysis of topic expression

We aimed to identify topics that are differentially associated with known cell type labels or disease conditions. For topic k and cell label j (i.e., cell type or disease condition), we first calculated the difference of the average topic activities between the cells with label j and the cells without label j. For each permutation trial, we randomly shuffled the label assignments among cells and recalculated the difference of average topic activities from the resulting permutation. The empirical p-value was calculated as (N’+1)/(N+1), where N’ is the number of permutation trials in which the difference is greater than the observed difference, and N is the total number of trials. To account for multiple hypotheses, we applied Bonferroni correction by multiplying the p-value by the product of the topic number and the number of labels. We performed N=100,000 permutations.

We determined a topic to be differentially expressed (DE) if the Bonferroni corrected q-value is lower than 0.01 and the mean difference is greater than 2 for cell-type DE topcis or 0.2 for disease DE topics. **Supp. Table S20** summarizes the number of DE topics we identified for each cell type and disease conditions from the AD and MDD data. We use the PanglaoDB database [71] to find the overlap between top genes of cell-type-specific DE topics and known cell type markers.

### Incorporation of pathway knowledge into the gene embeddings in p-scETM

We downloaded the pathDIP4 pathway database from [49], and the Gene Ontology (Biological Processes) (GO-BP) dataset from MSigDB v7.2 Release [69]. Pathway gene sets or GO-BP terms containing fewer than five genes were removed. We represented the pathway knowledge as a pathways-by-genes **ρ** matrix, where *ρ_ij_* = 1 if gene set *i* contains gene *j*, and *ρ_ij_* = 0 otherwise. We standardized each column (i.e., gene) of this matrix for numerical stability. During training the p-scETM, we fixed the gene embedding matrix **ρ** to the pathways-by-genes matrix.

### Clustering performance benchmark and visualization

We assessed the performance of each method by three metrics: Adjusted Rand Index (ARI) [72], Normalized Mutual Information (NMI) and k-nearest-neighbor Batch-Effect Test (kBET) [18]. ARI and NMI are widely-used representatives of two families of clustering agreement measures, pair-counting and information theoretic measures, respectively. A high ARI or NMI indicates a high degree of agreement for a given clustering result against the ground-truth cell type labels. We calculated ARI and NMI using the Python library scikit-learn [73].

kBET measures how well mixed the batches are based on the local batch label distribution in randomly sampled nearest-neighbor cells compared against the global batch label distribution. Average silhouette width (ASW) [40] indicates clustering quality using cell type labels. Silhouette width (SW) of a cell *i* is the distance of cell *i* from all of the cells within the same cluster subtracted by the distance of cell i from cells in a nearest but different cluster, normalized by the maximum of these two values. ASW is the averaged SW over all the cells in a dataset and larger values indicate better clustering quality. Therefore, larger ASW indicates the higher distances between cell types and lower distance within cell type in the cell embedding space. We choose the distance function to be the euclidean distance. We adapted the Pegasus implementation [74] for kBET calculation, and set the *k* to 15.

All embedding plots were generated using the Python scanpy package [16]. We use UMAP to reduce the dimension of the embeddings to 2 for visualization, and Leiden [75] to cluster cells by their cell embeddings produced by each method in comparison. During the clustering, we tried multiple resolution values and reported the result with the highest ARI for each method.

For reproducibility, the evaluation and the plotting steps were implemented in a single evaluate function in the scETM package, which takes in an AnnData object with cell embeddings and returns a Figure object for the ARI, NMI, kBET and embedding plot. For consistency, we used this function to evaluate all methods, including those written in R, where we used the reticulate package [76] to call our evaluate function.

We ran all methods under their recommended pipeline settings (**Supplementary Methods**), and we use batch correction option whenever applicable to account for batch effects. All results are obtained on a compute cluster with Intel Gold 6148 Skylake CPUs and Nvidia V100 GPUs. We limit each experiment to use 8 CPU cores, 192 GB RAM and 1 GPU.

### Efficiency and scalability benchmark of the existing methods

To create a benchmark dataset for evaluating the run time of each method, we merged MDD and AD, keeping the genes that appear in both datasets. We then selected the 3000 most variable genes using scanpy’s highly_variable_genes(n_top_genes=3000, flavor=‘seurat_v3’) function, and randomly sampled 28,000, 14,000, 70,000 and 148,247 (all) cells to create our benchmark datasets. The memory requirements reported in **Fig. 3** were obtained by reading the rss attribute of the named tuple returned by calling Process().memory_info() from the psutil Python package [77]. For methods based on R, we use the reticulate package [76] to call the above Python function for consistency. We used the same settings (RAM size, number of GPUs, etc) as described in the **Clustering performance benchmark and visualization** section throughout the experiments.

## Data Availability

The datasets analysed during the current study are from publicly available repositories or data portals. The acquisition and quality control steps for all datasets are included in the supplementary information.

## Code Availability

scETM source codes as well as the benchmarking workflows have been deposited at the GitHub repository (https://www.github.com/hui2000ji/scETM) and available in the zip file accompanied with the paper.

## Ethics approval and consent to participate

Not applicable.

## Consent for publication

Not applicable.

## Authors’ contributions

YL and JT conceived of the study. YZ, HC, YL analyzed and interpreted the data, wrote the manuscript, and wrote the code for scETM. HC optimized and completed the final scETM code. ZZ ran some initial experiments. YL and JT supervised the project. All authors approved the final manuscript.

## S1 Supplementary Methods

### S1.1 Data processing

All of the single-cell datasets used in this study are from publicly available repositories or data portals. We describe below the acquisition and quality control (QC) for each of the datasets used in the current work.

### S1.1.1 Human pancreatic islet

We obtained the human pancreatic islet dataset and the ground truth cell type labels from Satija Lab at the following link: https://satijalab.org/seurat/v3.0/integration.html (accessed 1 Dec 2020), originally deposited by Stuart *et al*. [7]. This dataset is a compilation of scRNA-seq data from five studies which can be accessed using the following Gene Expression Omnibus (GEO) accession numbers: GSE81076 (CelSeq), GSE85241 (CelSeq2), GSE86469 (Fluidigm C1), E-MTAB-5061 (SMART-Seq2), and GSE84133 (inDrop). A QC step was conducted by [7], and no additional QC was performed. In our benchmarking experiment, we use the different scRNA-seq technologies as the batch variable.

### S1.1.2 Mouse pancreatic islet

We obtained the mouse pancreatic islet data and ground truth cell type labels from GSE84133 (inDrops) without conducting additional QC. There are 1,886 mouse cells from two mice of different strains, ICR and C57BL/6 [34]. The cell counts from the two trains are of approximately equal proportions. In our benchmarking experiment, we treated the mouse strain as the batch variable because of the different genetic backgrounds.

### S1.1.3 Major Depressive Disorder (MDD)

We obtained the 10X Genomics-based MDD snRNA-seq dataset with ground truth cell type labels from GSE144136. A strict QC step was conducted in the original empirical study by [29], where cells with fewer than 110 detected genes were removed. The top 0.5% of cells based on the total number of UMI (unique molecular identifiers) detected in each cell were also excluded because they are likely to be multiplets rather than single nuclei. No additional QC was performed. The MDD dataset consists of 78,886 cells from the dorsolateral prefrontal cortex of 34 male participants. The participants in the control group (n=17) who died due to natural cause and case group (n=17) who died by suicide were matched for age (18–87 years), postmortem interval (12–93h) and brain pH (6–7.01) [29]. The number of cells from each donor is approximately the same.

### S1.1.4 Alzheimer’s Disease (AD)

We obtained the droplet-based AD snRNA-seq data and the corresponding ground truth cell type labels from Synapse (https://www.synapse.org/#!Synapse:syn18485175) under the doi 10.7303/syn18485175, and the metadata from https://www.synapse.org/#!Synapse: syn3157322. A strict QC step based on UMI counts and mitochondrial ratio values was conducted in the original empirical study by Mathys *et al.* [35]. The AD dataset consists of 70,634 cells from the prefrontal cortex of 48 individuals, both male and female, in the Religious Order Study (ROS) or the Rush Memory and Aging Project (MAP), two longitudinal cohort studies of aging and dementia. The cases group consists of 24 individuals with high levels of *β*-amyloid and other pathological hallmarks of AD, and the control group consists of 24 individuals who have no or very low *β*-amyloid or other pathologies.

Study data were provided by the Rush Alzheimer’s Disease Center, Rush University Medical Center, Chicago. Data collection was supported through funding by NIA grants P30AG10161 (ROS), R01AG15819 (ROSMAP; genomics and RNAseq), R01AG17917 (MAP), R01AG30146, R01AG36042 (5hC methylation, ATACseq), RC2AG036547 (H3K9Ac), R01AG36836 (RNAseq),

R01AG48015 (monocyte RNAseq) RF1AG57473 (single nucleus RNAseq), U01AG32984 (genomic and whole exome sequencing), U01AG46152 (ROSMAP AMP-AD, targeted proteomics), U01AG46161(TMT proteomics), U01AG61356 (whole genome sequencing, targeted proteomics, ROSMAP AMP-AD), the Illinois Department of Public Health (ROSMAP), and the Translational Genomics Research Institute (genomic). Additional phenotypic data can be requested at www.radc.rush.edu.

### S1.1.5 Tabula Muris

We obtained the *Tabula Muris* dataset with ground truth cell type labels from FigShare (https://figshare.com/projects/Tabula_Muris_Transcriptomic_characterization_of_20_organs_and_tissues_from_Mus_musculus_at_single_cell_resolution/27733) for the Version 2 release [3]. This dataset includes mouse single-cell transcriptome data sequenced by two tech-nologies: microfluidic droplet-based, and fluorescence-activated cell sorting (FACS)-based. A QC cutoff was applied in the original empirical study where only cells with at least 500 genes and 50,000 reads/ 1000 UMI are kept. The droplet subset includes data for 422,803 droplets, 55,656 of which passed the QC cutoff. The FACS subset, denoted as “TM (FACS)” in the paper, contains data for 53,760 cells, 44,879 of which passed the QC cutoff.

### S1.1.6 Allen Brain Atlas

We downloaded two brain datasets from Allen Brain Atlas [78] (accessed 03/21/2021). The human primary motor cortex dataset (HumM1C) includes single-nucleus transcriptomes from 76,533 nuclei from the primary motor cortex (M1C) of 2 post-mortem human brain specimens. In total, 127 transcriptomic cell types are present in this dataset. The sample processing follows the 10x Genomics pipeline. To generate the ground-truth cell labels, the default 10x Cell Ranger v3 pipeline was used except substituting the curated genome annotation used for SMART-seq v4 quantification. The mouse brain dataset includes single-cell transcriptomes from more than 20 areas of mouse cortex and the hippocampus, which has 1,093,785 cells in total. Samples were collected from male and female mice around 8 week-old, from panneuronal transgenic lines. The cell transcriptomes were sequenced using 10x Genomics. We used the subclass labels in the metadata used as ground truth cell type label in the current study. We removed the cells and nuclei with subclass label “outlier”. To encourage better transfer from HumM1C, we subset the mouse brain dataset by keeping the 124,953 cells with region_label “MOp”, and obtain the MusMOp dataset.

### S1.1.7 Mouse Retina

We obtained the MR dataset from https://hemberg-lab.github.io/scRNA.seq.datasets/ mouse/retina/, which is a collection of mouse retina scRNA-seq datasets from two independent studies, namely Macosko *et al*. [36] with 44808 samples, and Shekhar *et al*. [37] with 27499 samples. Both subsets were sequenced using Drop-seq. We kept the genes shared by the two subsets, and filtered out the 669 samples labeled as “Doublets/Contaminants” in the Shekhar batch. The merged dataset contains 71638 cells and 12333 genes.

### S1.2 Experimental details of other scRNA-seq methods

Neural-network based models, including scVI, scVI-LD, scVAE-GM and scETM typically need at least 5000 gradient updates to converge. When running on small datasets, the total number of gradient updates per epoch may be very small (sometimes as low as 1). In these cases, we increase the number of epochs *T* to ensure the model goes through at least 12000 gradient updates, i.e. *T* = max(800, 12000 *<u>^B^</u>*), where *N* is the number of cells in the dataset and *B* is the mini-batch size.

### S1.2.1 Seurat v3

We downloaded Seurat v3 (version 3.1.5) from CRAN [8]. We followed the steps outlined by the integration workflow (https://satijalab.org/seurat/v3.2/integration.html) which includes NormalizeData, FindVariableFeatures, FindIntegrationAnchors, and IntegrateData. To make the comparisons more equitable, we set the min.features=0 to avoid exclusion of cells. All other parameters were set as default. We noted that, with batch integration turned on, Seurat reports error in the integration step due to the high number of anchors arising from the 48 individuals (batch variable in AD), which is a known implementation issue with the standard Seurat v3 integration workflow [79]. We therefore turned off the batch integration for AD in the benchmarking experiments (see **Clustering performance benchmark and visualization** and **Efficiency and scalability benchmark of the existing methods**) and followed the steps described in the Guided Clustering Tutorial (https://satijalab.org/seurat/v3.2/pbmc3k_ tutorial.html).

### S1.2.2 Scanorama

We downloaded the source code from GitHub brianhe/scanorama. We used the integrate_scanpy function for dataset integration and batch correction as suggested by the guided tutorial. All parameters were set as default. The algorithm performs a PCA on the stacked datasets and uses 100 PCs for downstream computation.

### S1.2.3 Harmony

We downloaded the source code from GitHub slowkow/harmonypy suggested by the primary repository immunogenomics/harmony and followed the preprocessing (normalization and top variable gene selection) described in the publication and the integration steps in the provided tutorial. We used the run_harmony function to obtain the corrected PCA embeddings and used 50 PCs as input. All other parameters were set as default.

### S1.2.4 LIGER

We used the official implementation provided on the website of the LIGER package. For the convenience of implementation, we followed the usage tutorial using Seurat Wrapper to process the raw data and then ran LIGER with default parameters.

### S1.2.5 scVI/scVI-LD

We downloaded the implementation from the Github repository YosefLab/scVI. We used the default model, which has one layer for both the encoder and decoder (for scVI-LD the decoder is a latent dimensions-by-genes matrix), 128 hidden units, 10 latent dimensions and ZINB distribution for modeling the data. We chose 10*^−^*^3^ as the learning rate and trained on each unprocessed dataset for 400 epochs, following the provided tutorials. We change the training batch size to 2000 for faster training. We obtained the cell embeddings via the get_latent method. We also compared the performance of scVI(-LD) with 10 and 100 latent dimensions on the five benchmark scRNA-seq datasets (**Supp. Table S21**). We found that in most cases, scVI(-LD) with 10 latent dimensions performs better than scVI(-LD) with 100 latent dimensions, justifying the hyperparameter choices made by the scVI(-LD) authors. When evaluating the interpretability aspect of scVI-LD, we use 100 latent dimensions for fair comparison with scETM.

### S1.2.6 scVAE

We downloaded the implementation from GitHub repository scvae/scvae. We set the hidden units to be (256, 128) for the encoder. The decoder is symmetric to the encoder. Latent dimension was set to 128 to match scETM. We chose 10*^−^*^4^ as the learning rate and NB distribution for modeling the data following the authors’ recommendation. We trained on each unprocessed dataset for 400 epochs with batch size of 250, including a 200-epoch warm-up for the KL divergence loss. In the scalability benchmark, we disabled the time-consuming per-epoch check-points to match other methods. The model did not converge on the Human Pancreatic Islet dataset, where the ELBO went to infinity. It failed to extract meaningful information from the *Tabula Muris* dataset, resulting in an ARI of 0.0.

## Supplementary Figures

**Figure S1:**
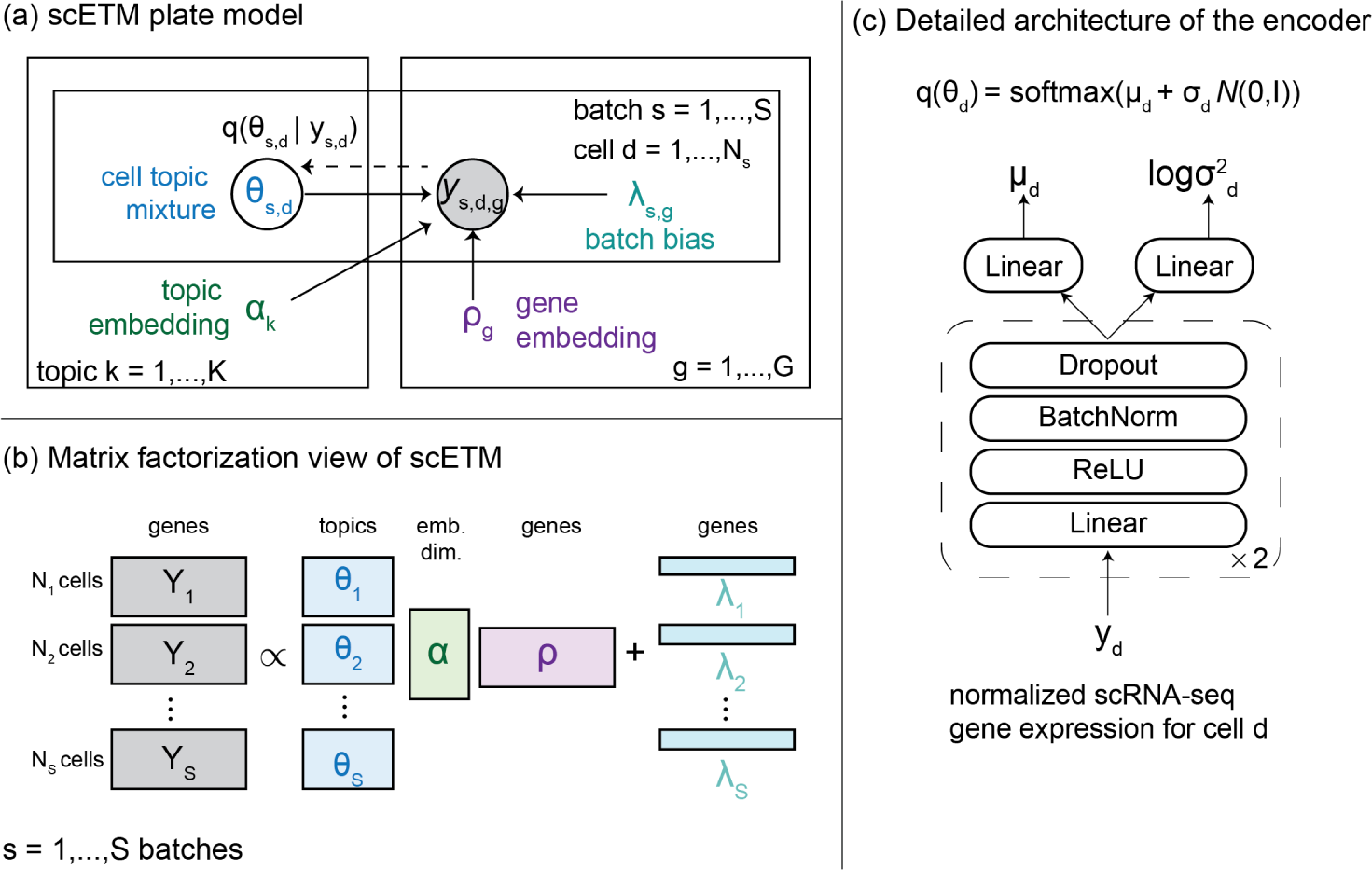
scETM model details. (a) The plate model for scETM. We model the scRNA-profile count matrix *y_d,g_* in cell *d* and gene *g* across *S* batches by a multinomial distribution with the rate parameterized by cell topic mixture *θ*, topic embedding *α*, gene embedding *ρ*, and batch effects *λ*. (b) Matrix factorization view of scETM. (c) Encoder architecture for inferring the cell topic mixture *θ*.

**Figure S2:**
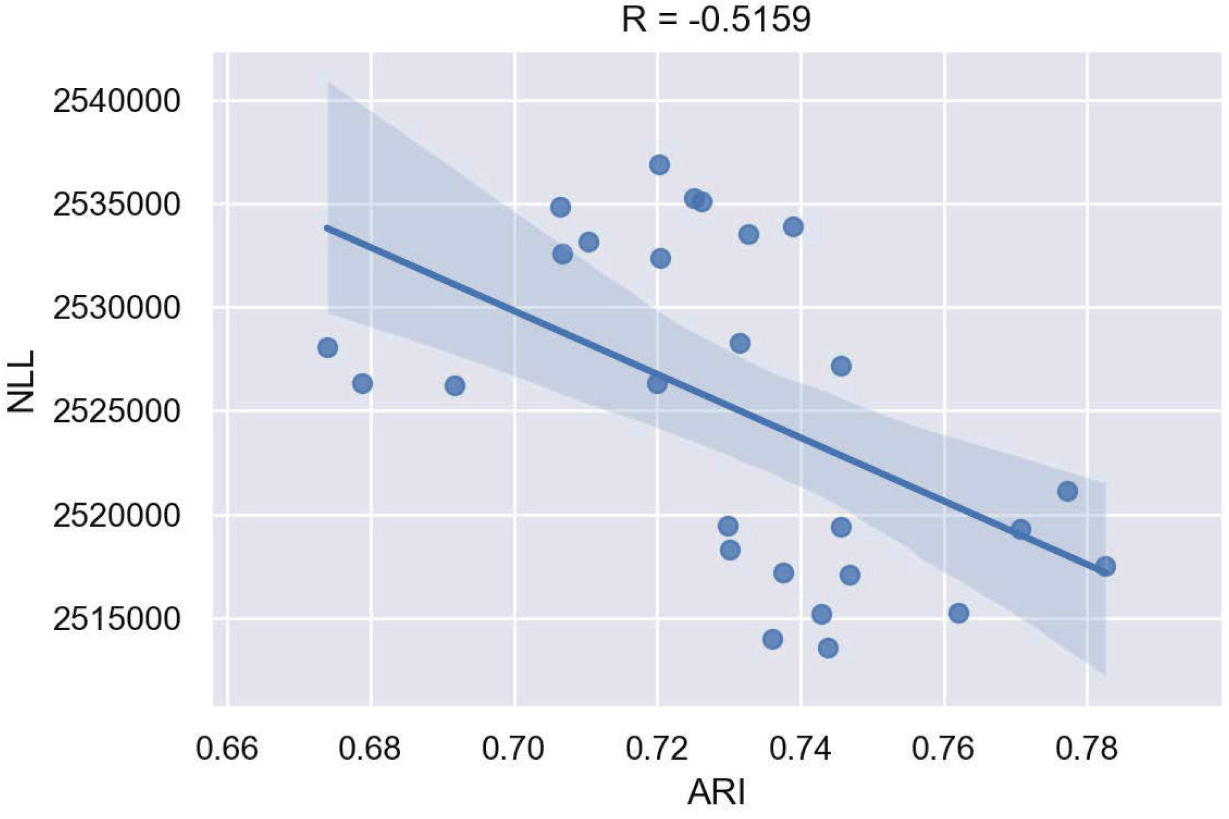
Relationship between average negative log-likelihood (NLL) and adjusted Rand Index (ARI) on the TM dataset. Each point denotes the performance of a trained scETM instance with a specific hyperparameter configuration (e.g., encoder hidden size, topic number, embedding dimensions, etc), averaged over three runs with different random seeds.

**Figure S3:**
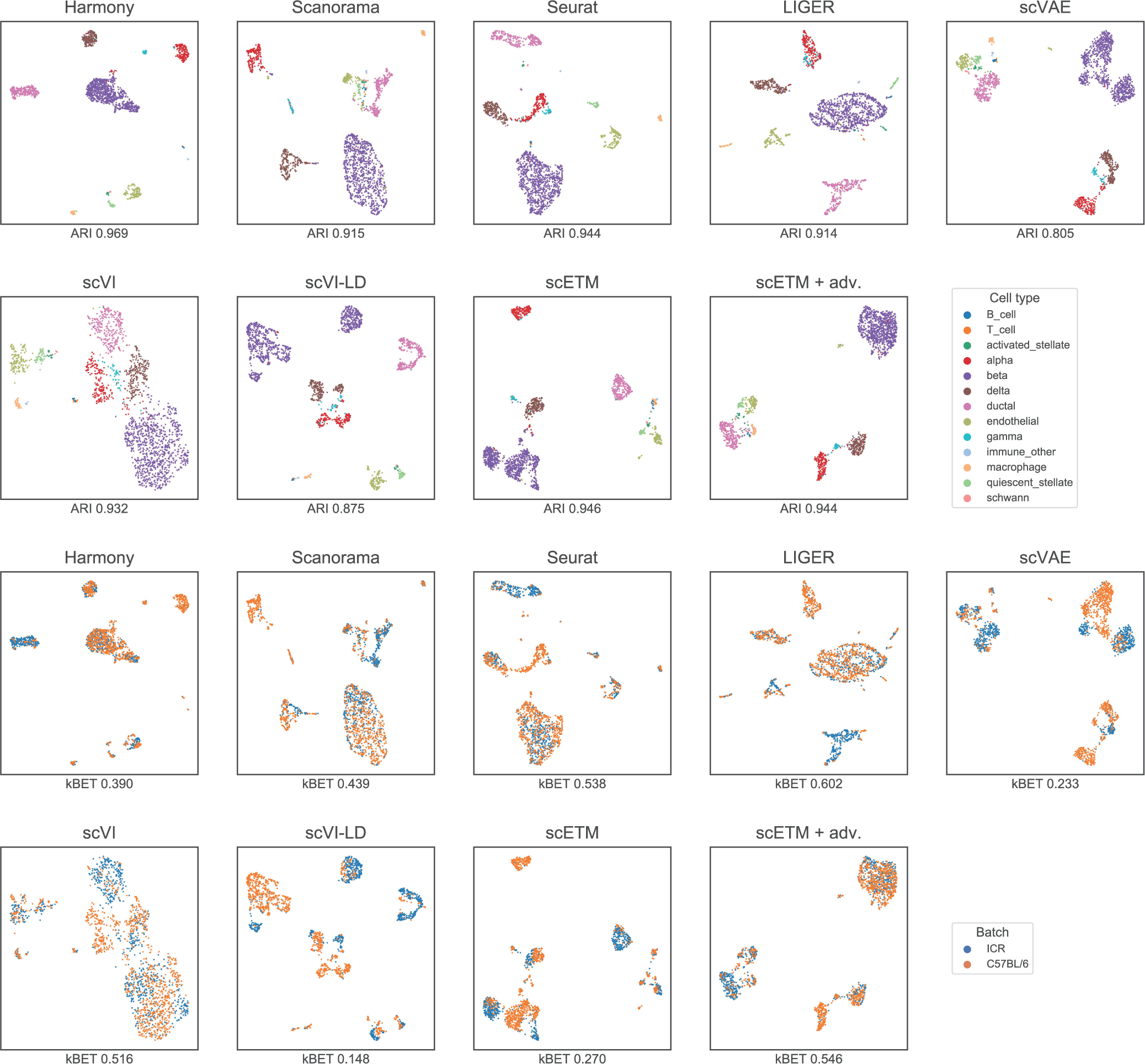
Integration and batch correction on the Mouse Pancreas (MP) dataset. Each panel shows the MP cell clusters using UMAP based on the cell embeddings obtained by each of the 9 methods. The cells are colored by cell types in the first two rows and by batches, which are the two mouse strains, in the last two rows.

**Figure S4:**
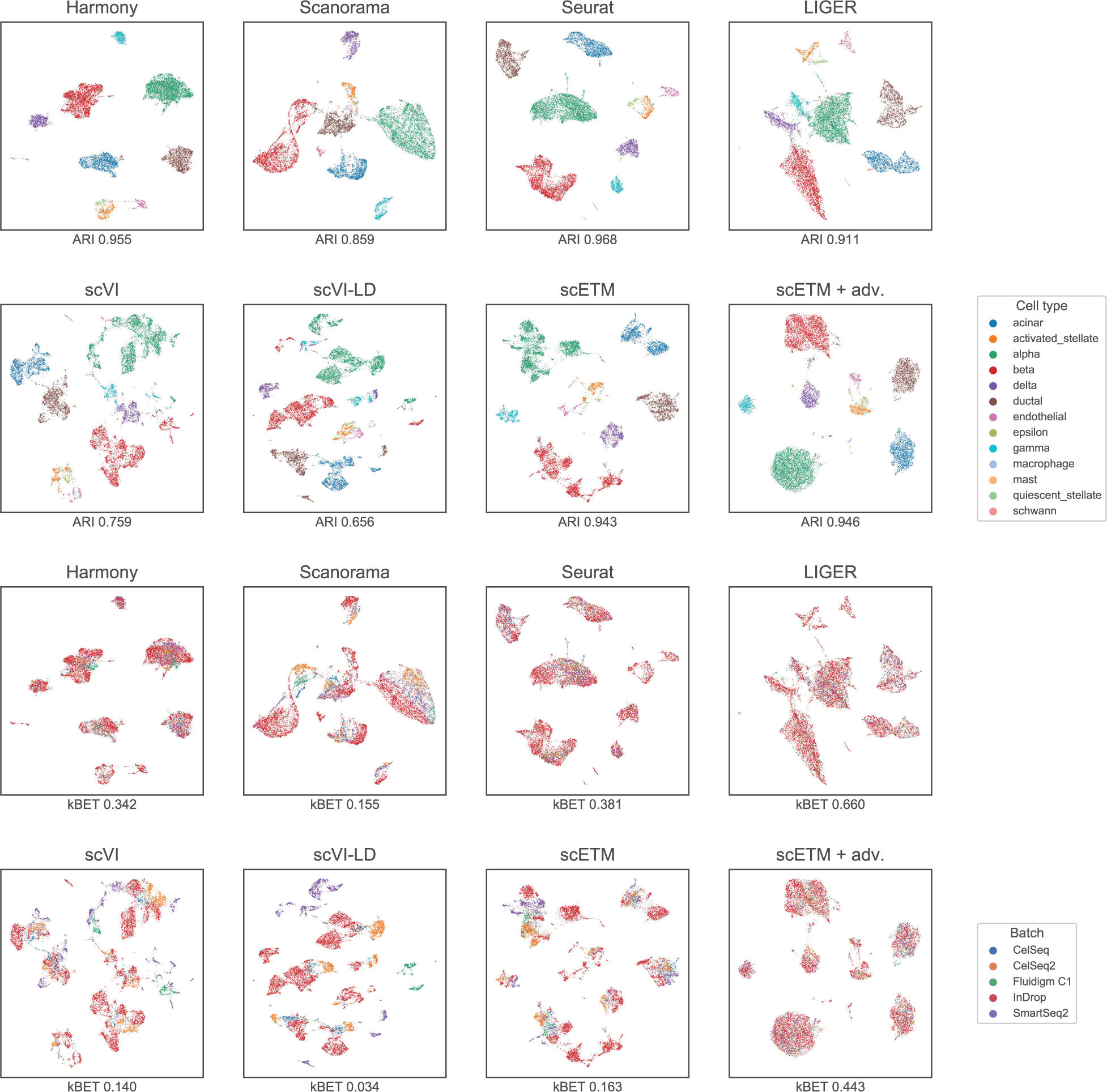
Integration and batch correction on the Human Pancreas (HP) dataset. Each panel shows the HP cell clusters using UMAP based on the cell embeddings obtained by each of the 9 methods. The cells are colored by cell types in the first two rows and by batches, which are the five sequencing technologies, in the last two rows.

**Figure S5:**
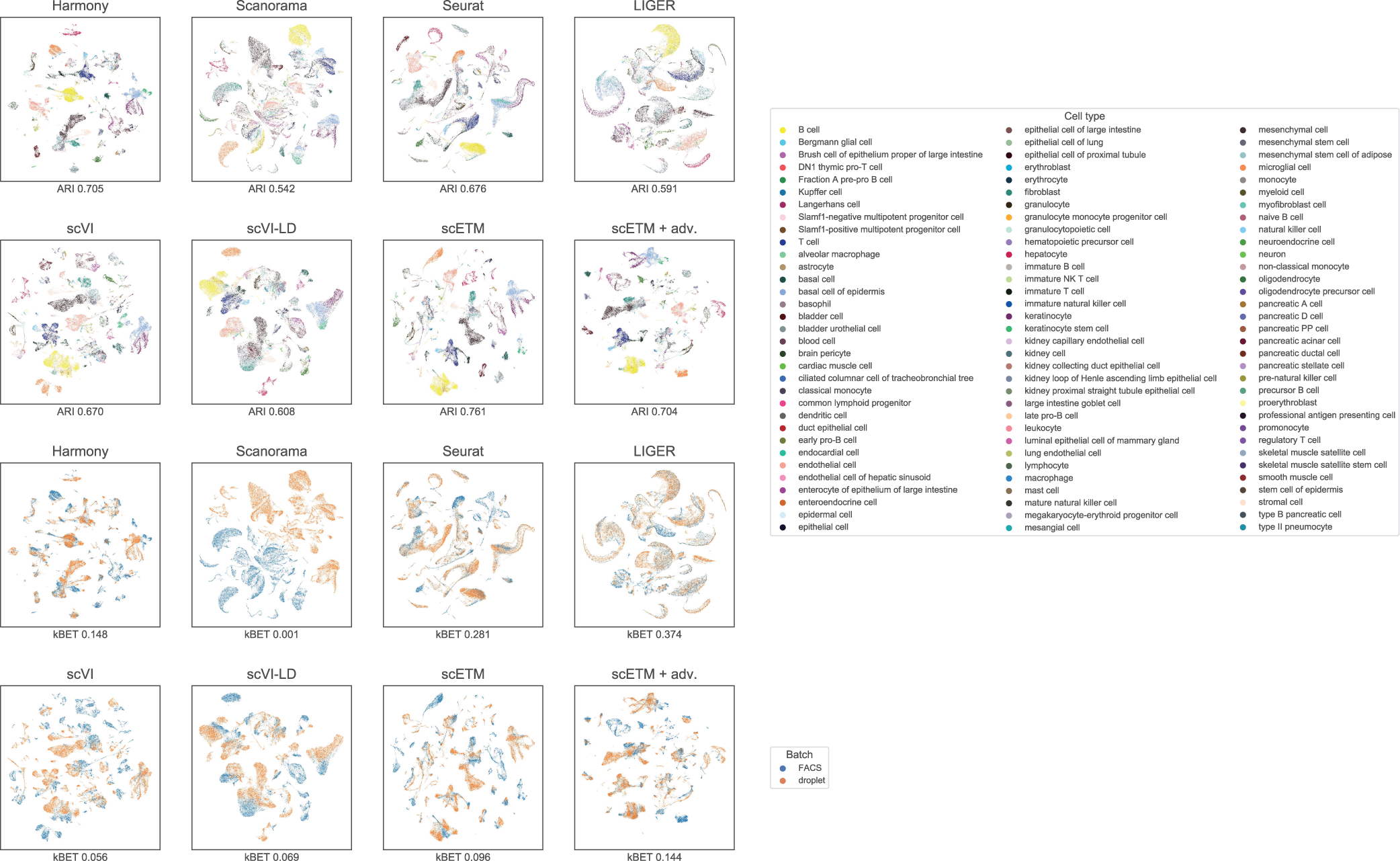
Integration and batch correction on the Tabula Muris (TM) dataset. Each panel shows the TM cell clusters using UMAP based on the cell embeddings obtained by each of the 9 methods. The cells are colored by cell types in the first two rows and by batches, which are the two sequencing technologies, in the last two rows.

**Figure S6:**
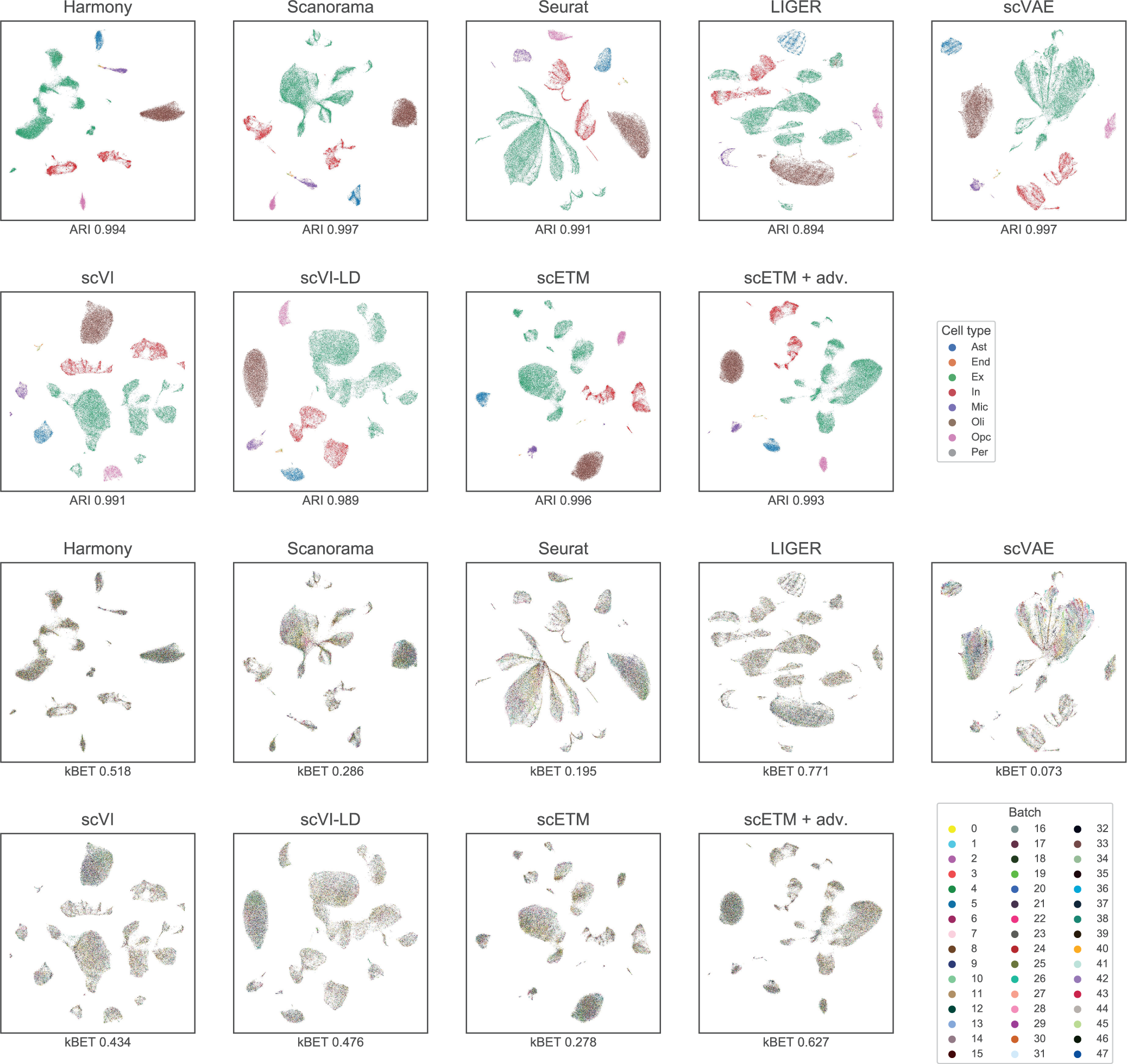
Integration and batch correction on the Alzheimer’s Disease (AD) dataset. Each panel shows the prefrontal cortex (PFC) cell clusters using UMAP based on the cell em- beddings obtained by each of the 9 methods. The cells are colored by cell types in the first two rows and by batches, which are the 48 donors, in the last two rows.

**Figure S7:**
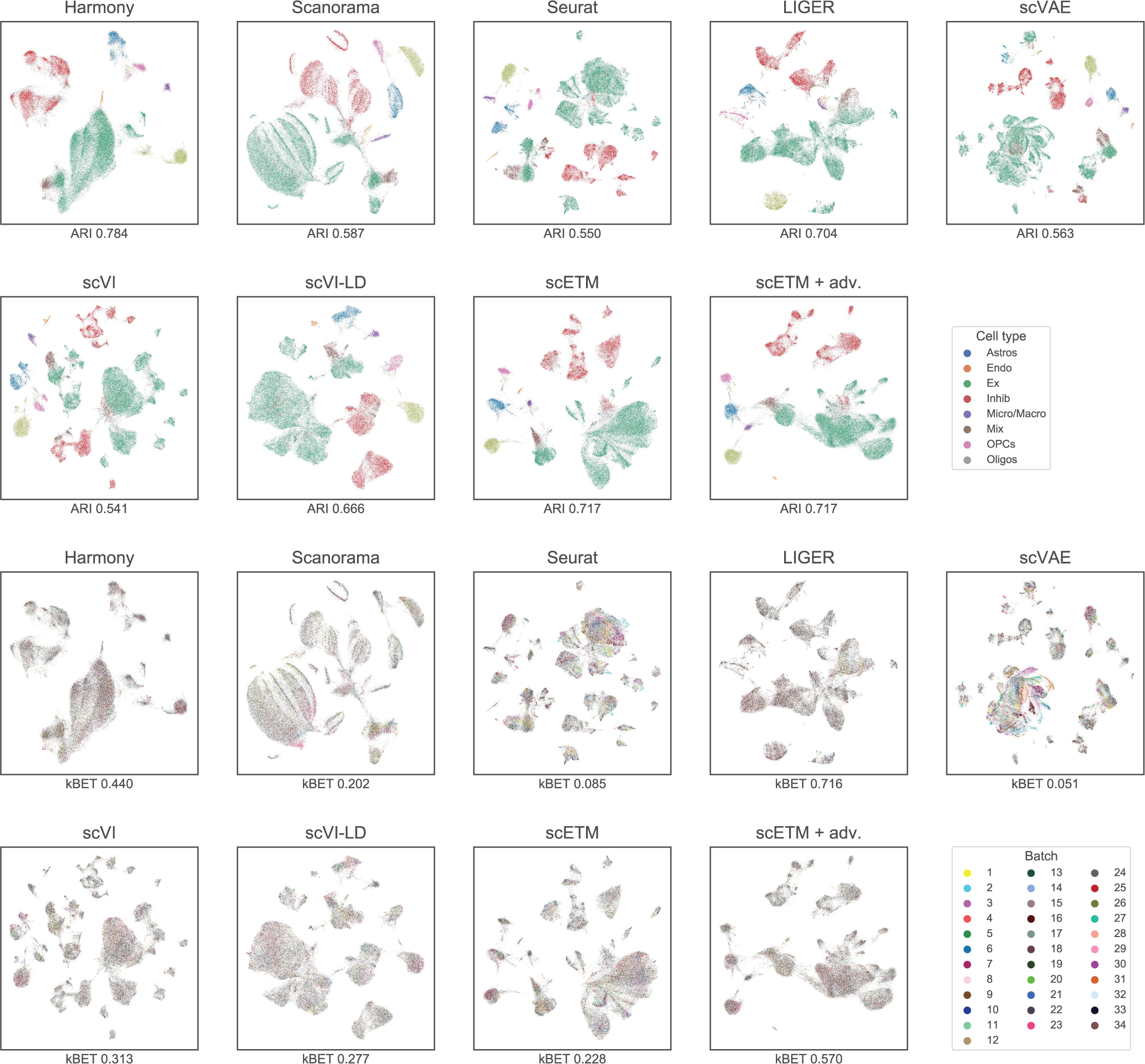
Integration and batch correction on the Major Depressive Disorder (MDD) dataset. Each panel shows the prefrontal cortex (PFC) cell clusters from the MDD or healthy subjects using UMAP based on the cell embeddings obtained by each of the 9 methods. The cells are colored by cell types in the first two rows and by batches, which are the 34 donors, in the last two rows.

**Figure S8:**
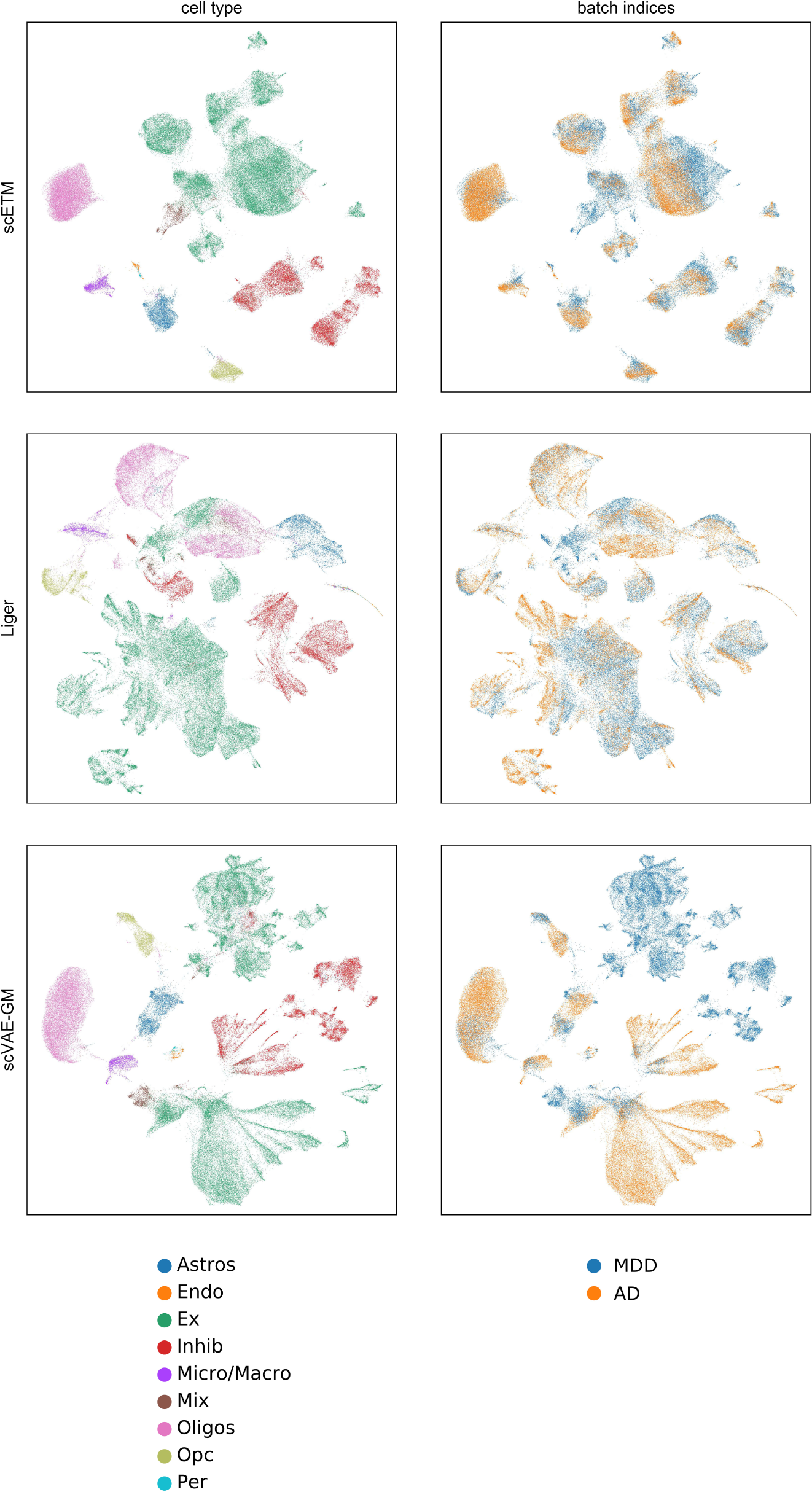
UMAP visualization of scETM, LIGER and scVAE-GM cell embeddings on the combined MDD and AD benchmark dataset, which includes in total 148247 cells and 3000 genes.

**Figure S9:**
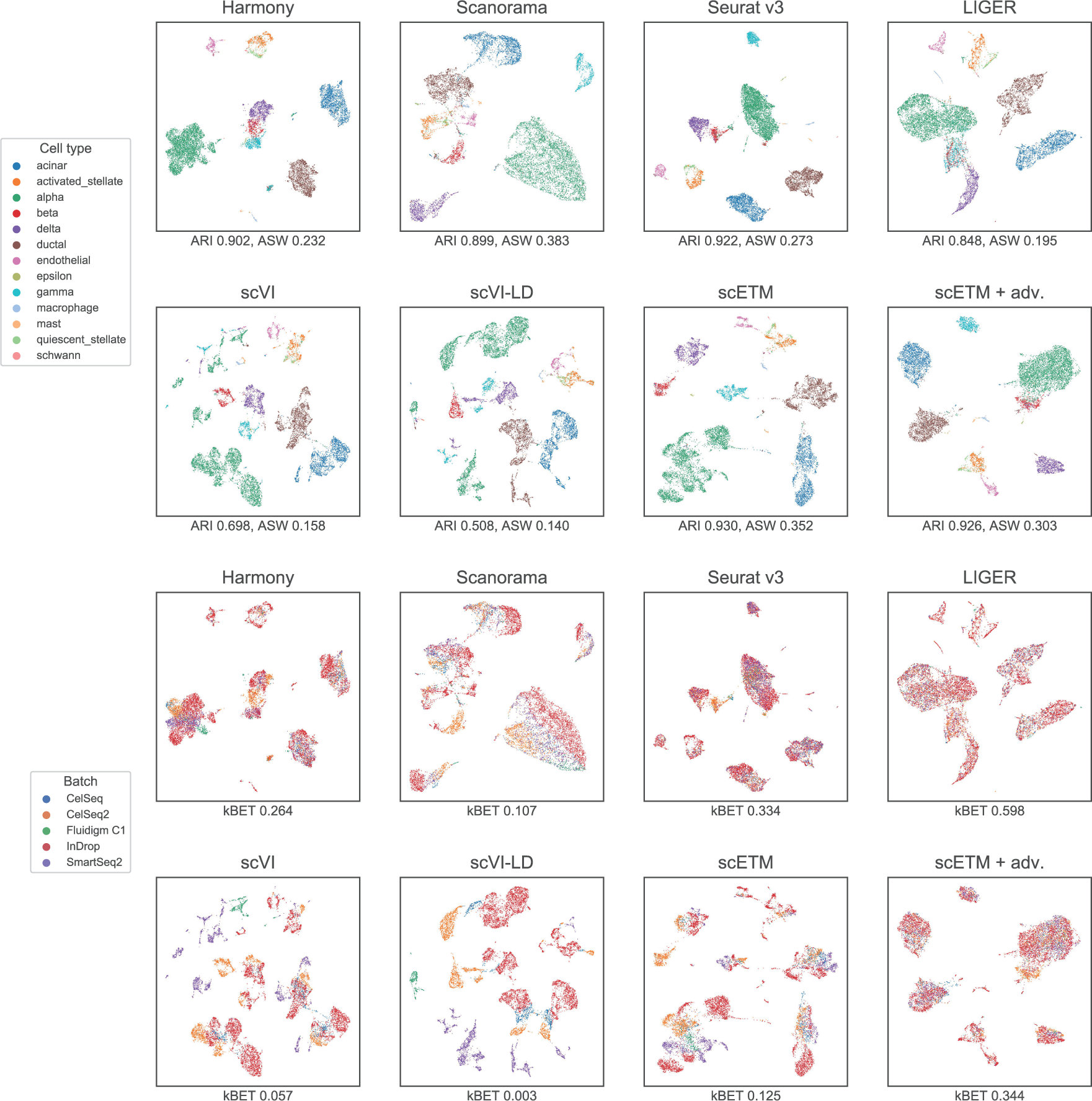
Integration and batch correction on the HP–beta dataset. Each panel shows the HP–beta cell clusters using UMAP based on the cell embeddings obtained by each of the 8 methods (scVAE did not converge on this dataset). The cells are colored by cell types in the first two rows and by batches, which are the five sequencing technologies, in the last two rows. The ARI and kBET scores of each method are shown below each plot.

**Figure S10:**
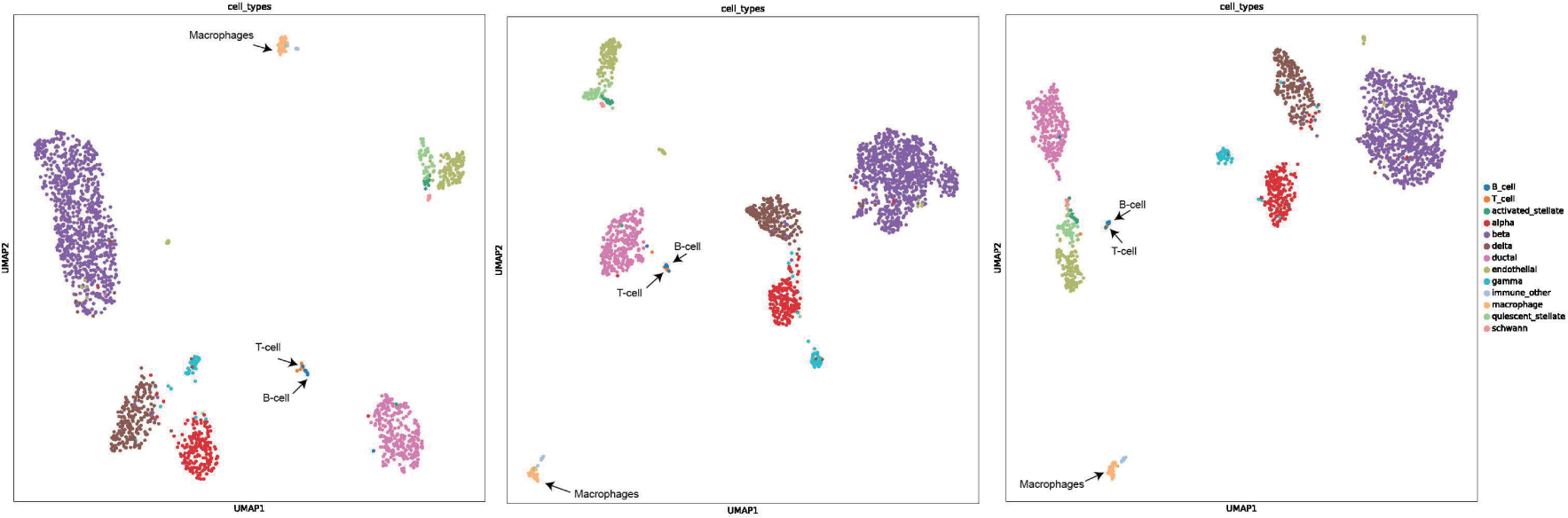
Reproducibility of the separation of T-cells and B-cells from macrophages. We trained scETM on TM (FACS) and applied it to the MP data. To assess the reproducibility, we repeated the experiments 3 times using different random seeds to initialize the model.

**Figure S11:**
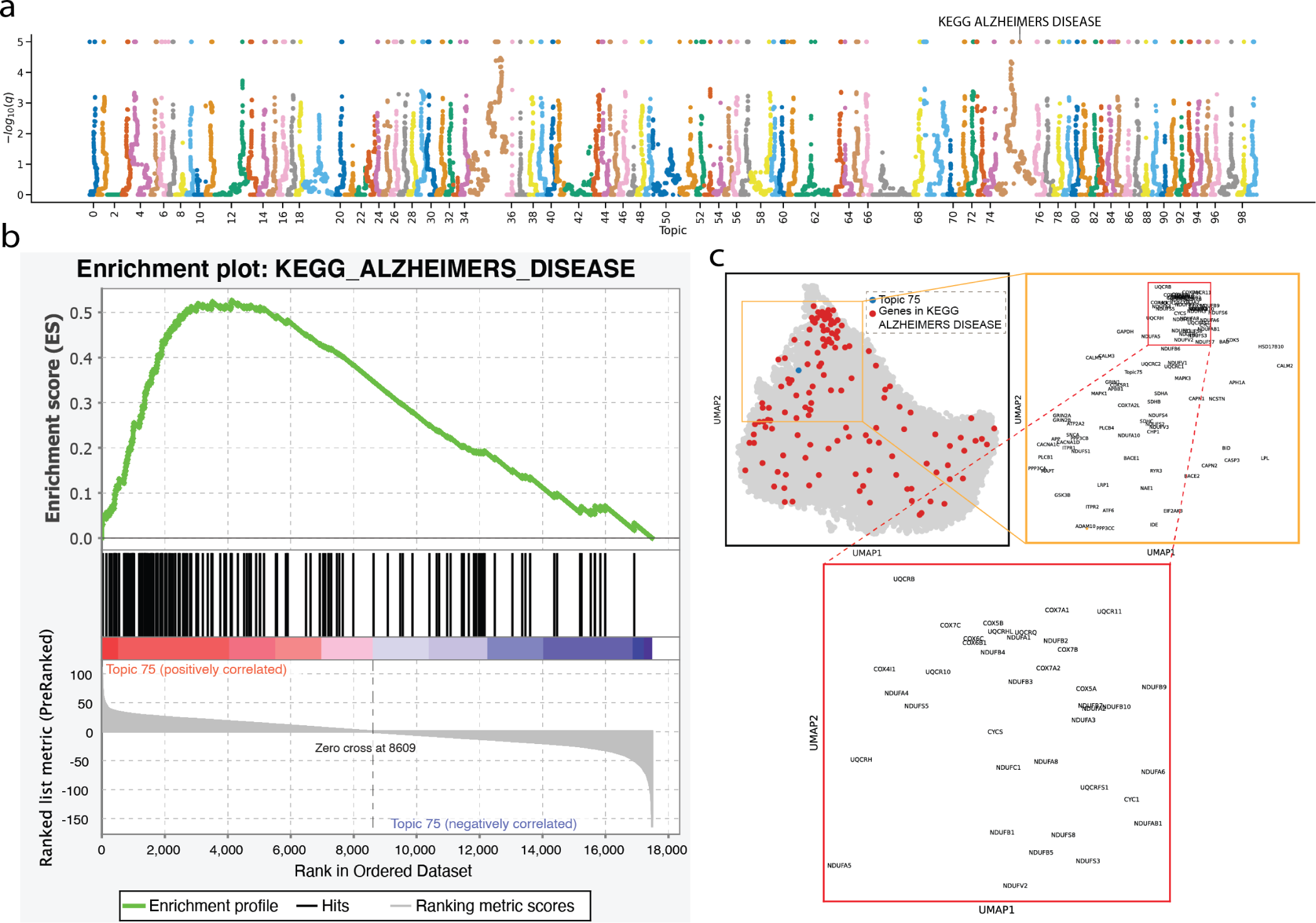
Alzheimer’s Disease dataset gene and topic embedding. **(a)** Manhattan plot of the GSEA results on the 100 scETM topic learned from the AD dataset. The maximum - log q-value is capped at 5. **(b)** Leading edge analysis of KEGG_ALZHEIMERS_DISEASE (KAD) pathway. The genes are ranked by the topic score (i.e., the unnormalized topic mixture beta) under Topic 75. Topic 75 is significantly enriched in KAD pathway (GSEA permutation test q-value=0). **(c)** UMAP visualization of the embeddings of all genes in AD and Topic 75. A magnified view of the cluster shows the KAD pathway genes that are near Topic 75.

**Figure S12:**
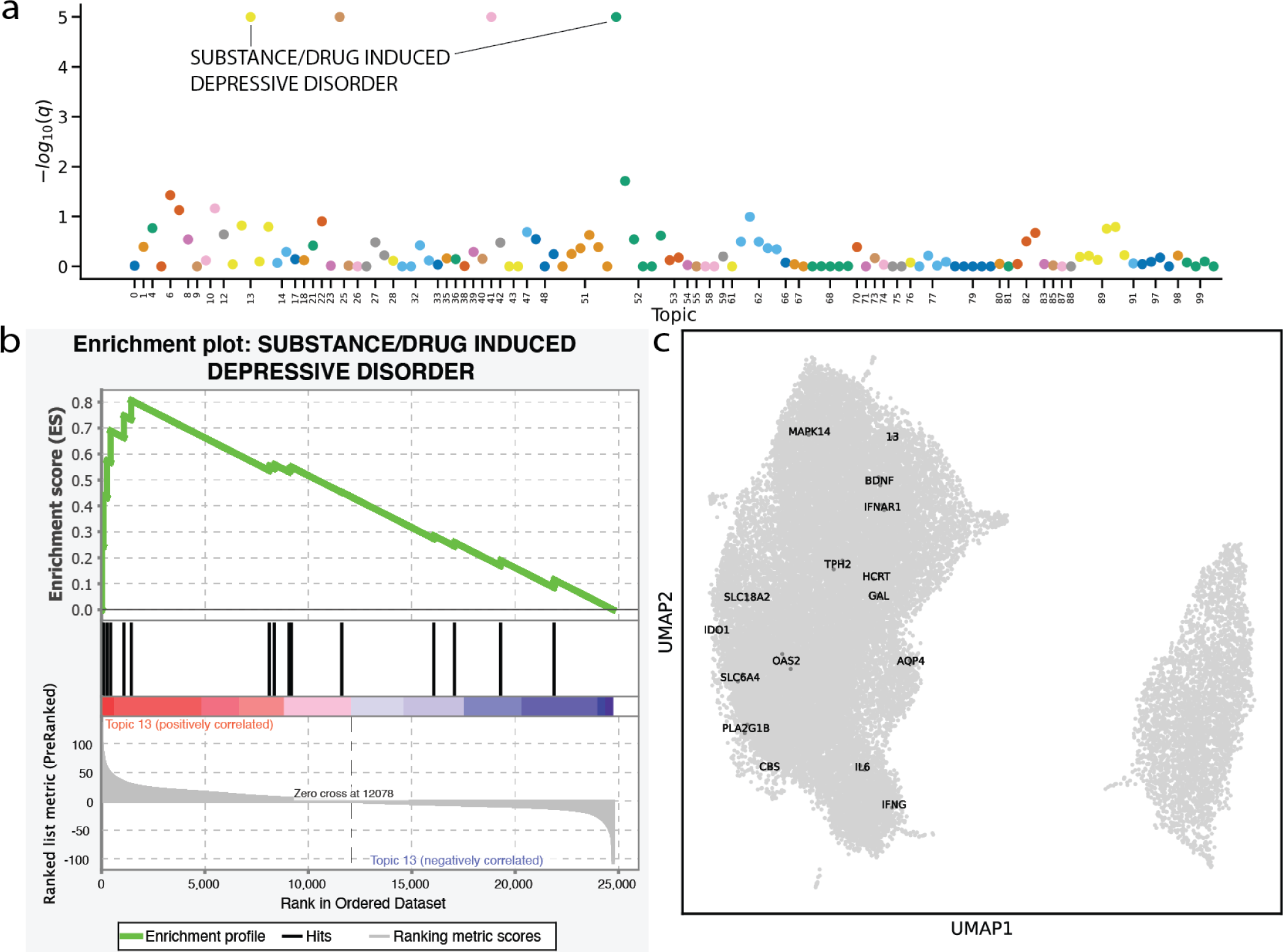
MDD dataset gene and topic embedding. **(a)** Manhattan plot of the GSEA re- sults on the 100 scETM topic learned from the MDD dataset. **(b)** Leading edge analysis of SUBSTANCE/DRUG INDUCED DEPRESSIVE DISORDER (SID) pathway. Topic 13 is signifi- cantly enriched in SID pathway (GSEA permutation test q-value=0). **(c)** UMAP visualization of the embeddings of all genes in the MDD dataset and Topic 13. A magnified view of the cluster shows the SID pathway genes that are near Topic 13.

**Figure S13:**
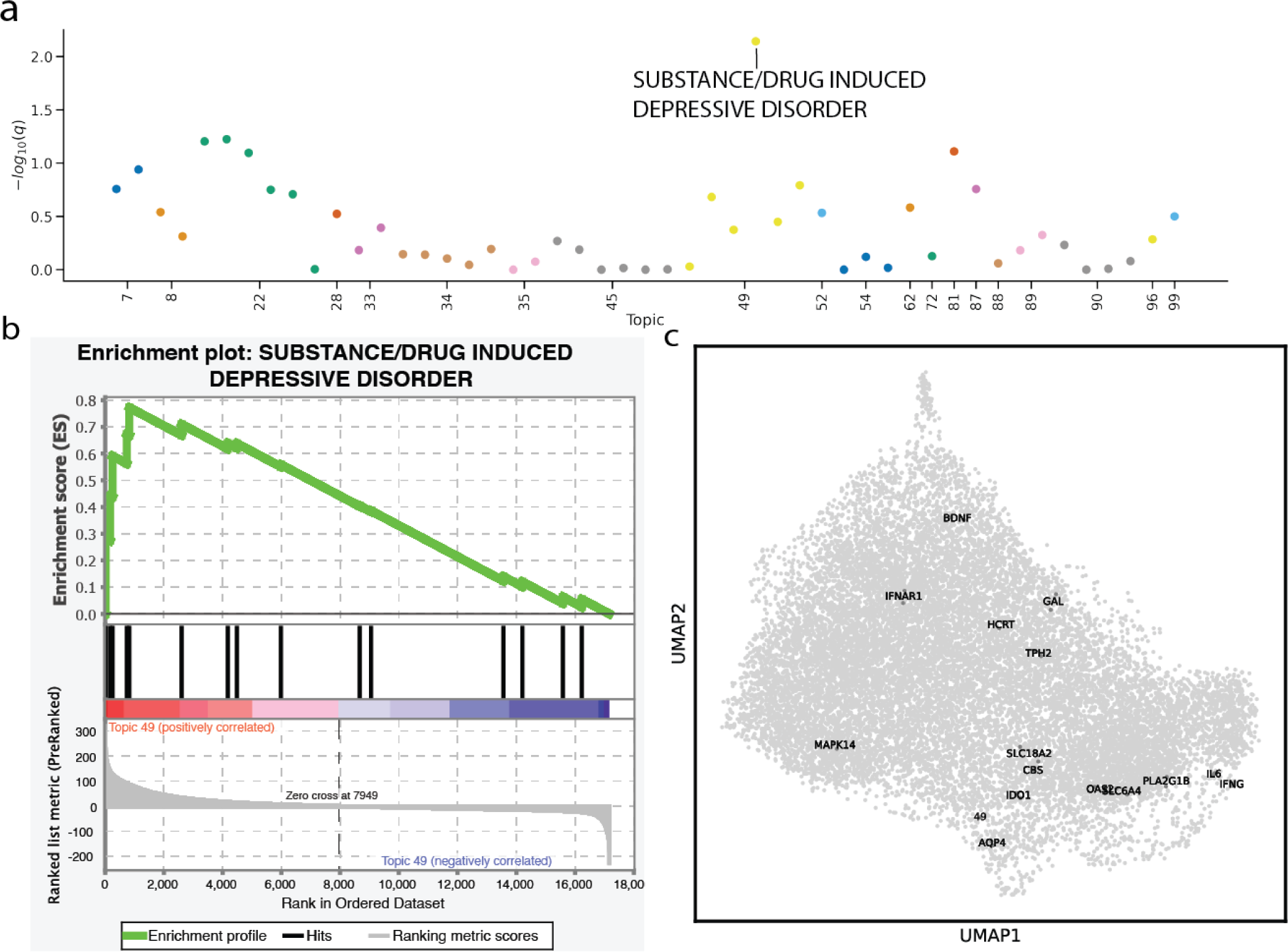
Gene and topic embedding of MDD (coding genes only). **(a)** Manhattan plot of the GSEA results on the 100 scETM topic learned from the MDD dataset. **(b)** Leading edge analysis of SUBSTANCE/DRUG INDUCED DEPRESSIVE DISORDER (SID) pathway. Topic 49 is significantly enriched in SID pathway (GSEA permutation test q-value=0). **(c)** UMAP visualization of the embeddings of all protein coding genes in MDD dataset and Topic 49. A magnified view of the cluster shows the SID pathway genes that are near Topic 49.

**Figure S14:**
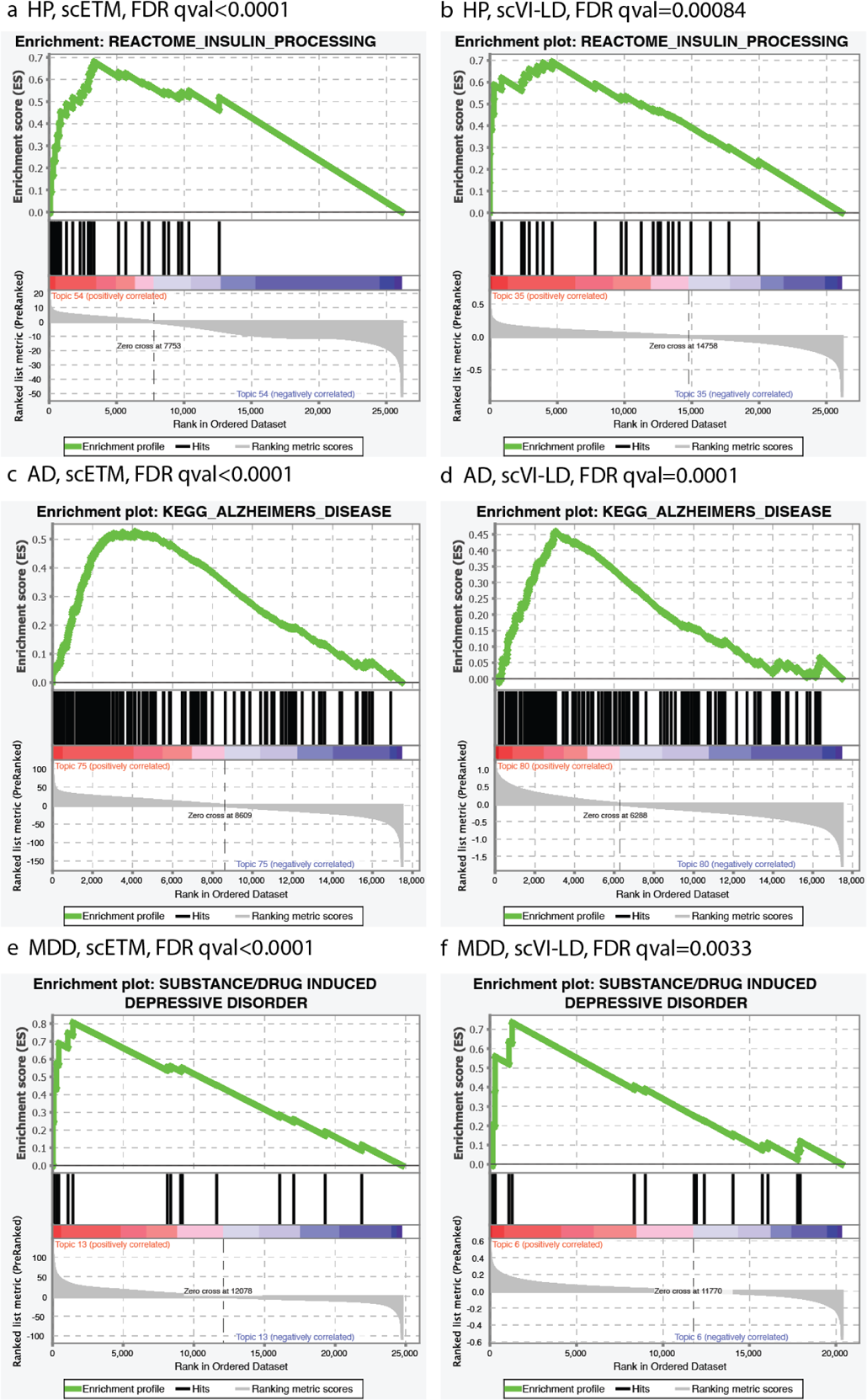
GSEA leading edge plots for scETM (left column) and scVI-LD (right col- umn) on Human Pancreas (HP), Alzheimer’s Disease (AD), and Major Depressive Disor- der (MDD). Due to the disease relevance, we showed Reactome Insulin Processing, KEGG Alzheimer’s Disease, and PsyGeNet substance/drug induced depressive disorder for HP, AD, and MDD, respectively.

**Figure S15:**
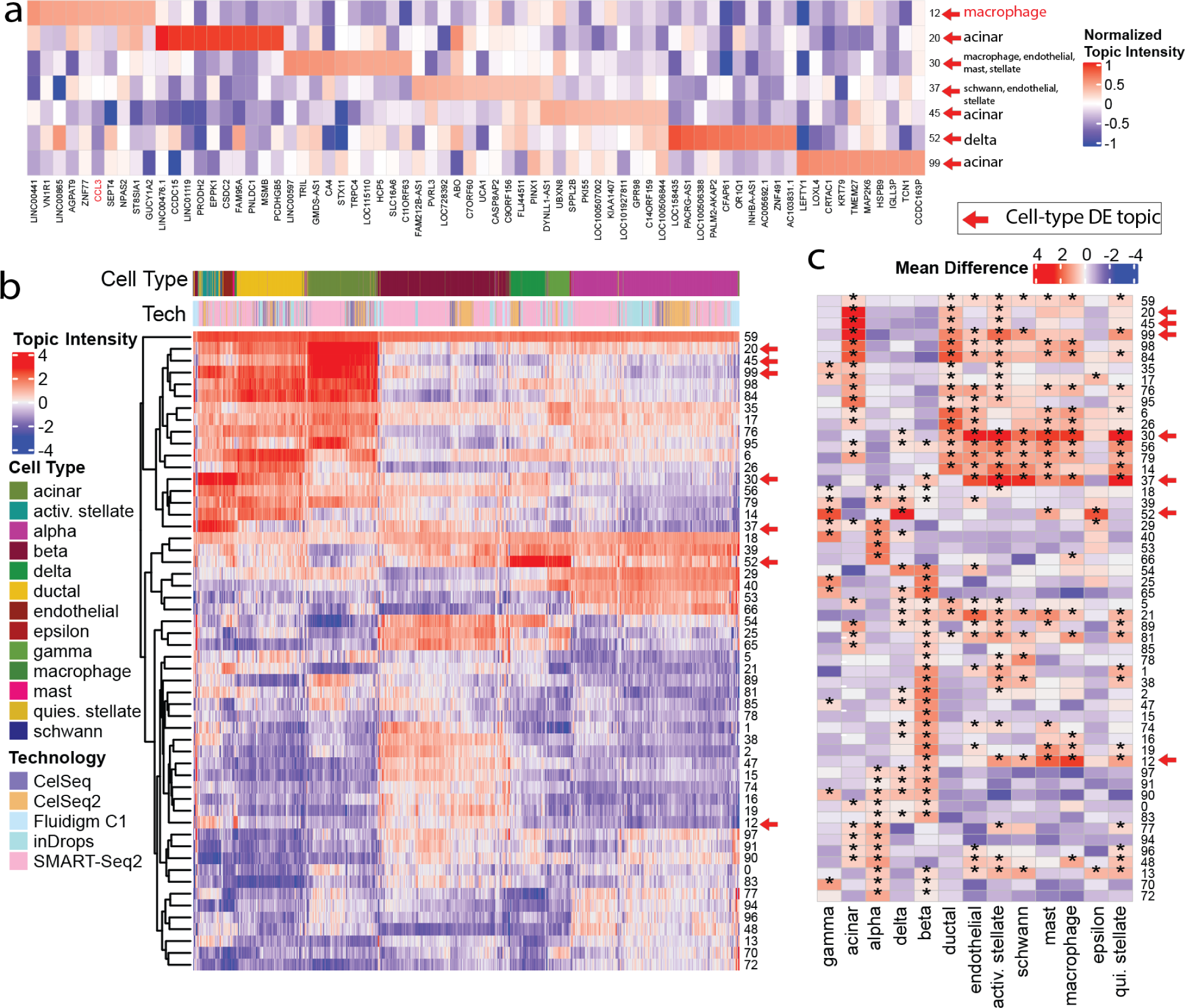
scETM topic embeddings of Human Pancreas scRNA-seq data. **(a)** Gene-topics heatmap of top 10 genes in each topic based on topic intensity. The top genes which are known as cell-type marker genes based on PanglaoDB are highlighted (rows). For visualization purposes, we divided the topic values by the maximum absolute value within the same topic. Only cell-type and disease differntial topics are shown. **(b)** Topic intensity of cells (n=10,000) sub-sampled from the HP dataset. Topic intensities shown here are the Gaussian mean before applying softmax. Only the select topics with the sum of absolute values greater than 1500 across all sampled cells are shown. The two color bars show cell types and batch identifiers (i.e., sequencing technologies). **(c)** Differential expression analysis of topics across the 13 cell types. Colors indicate mean differences between cell groups with and without a certain label (cell-type). Asterisks indicate Bonferroni-corrected empirical p-value < 0.05 for permutation tests of up-regulated topics in each cell-type labels. The number of permutations in each test is 100,000.

**Figure S16:**
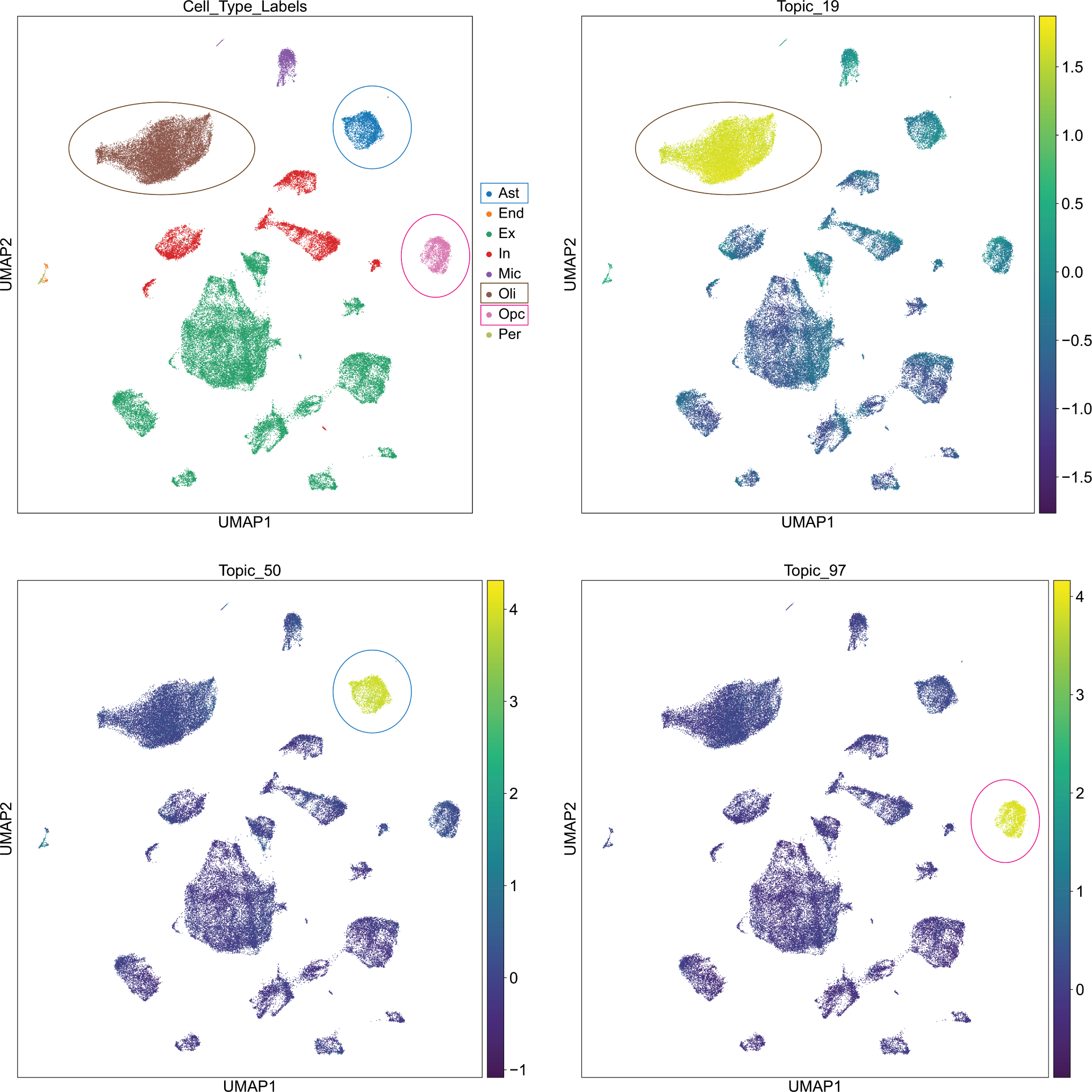
UMAP cell embedding visualization on the AD dataset, colored by differentially expressed topics (or *metagenes*) and ground truth labels for the cell types. Circled cell clusters has been discussed in the main text (see **Differential scETM topics in disease conditions and cell types** section).

**Figure S17:**
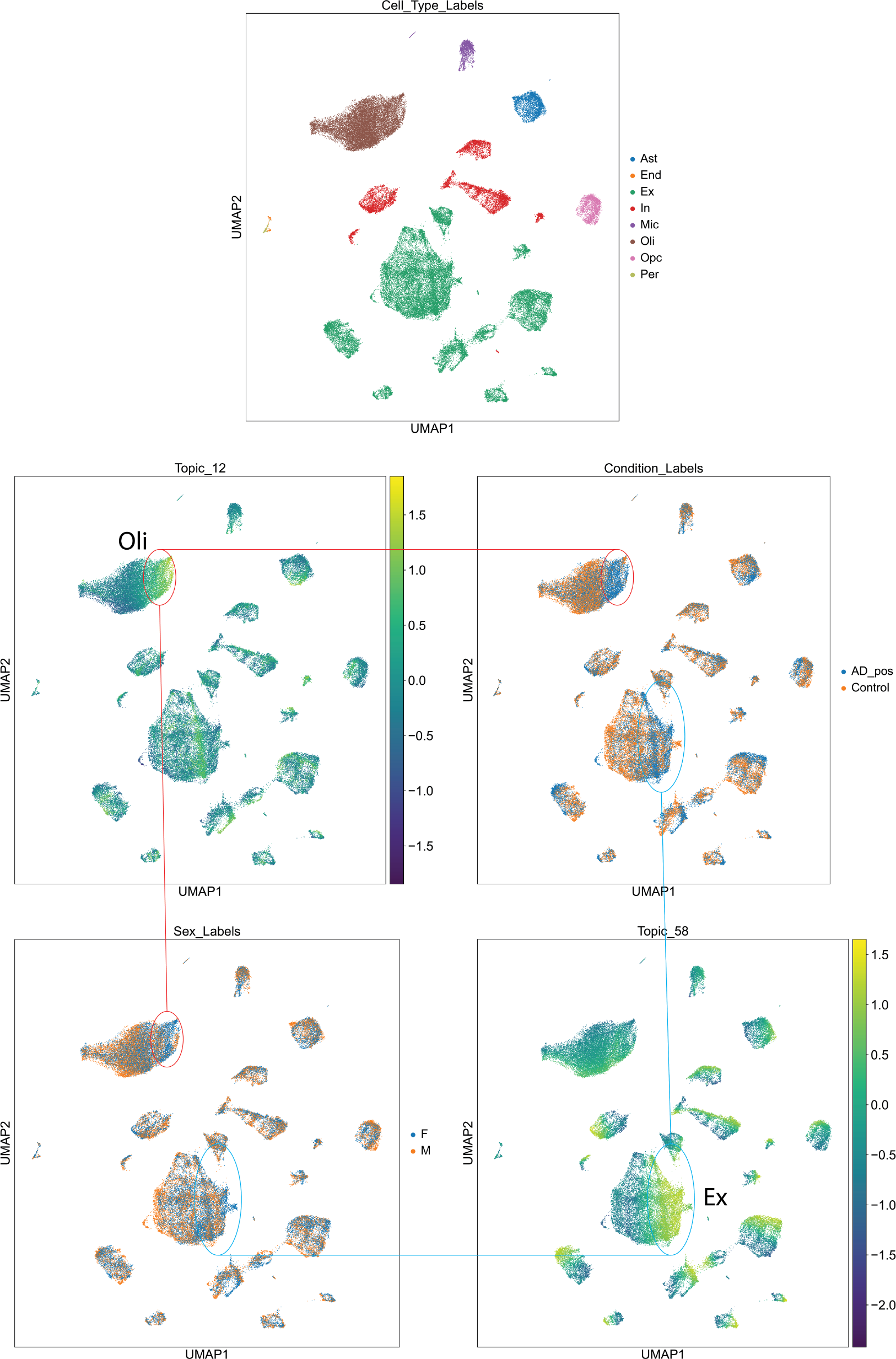
UMAP cell embedding visualization on the AD dataset, colored by differentially expressed topics (or *metagenes*) and AD/control or Male/Female labels. Circled cell clusters were discussed in the main text (see **Differential scETM topics in disease conditions and cell types** section).

**Figure S18:**
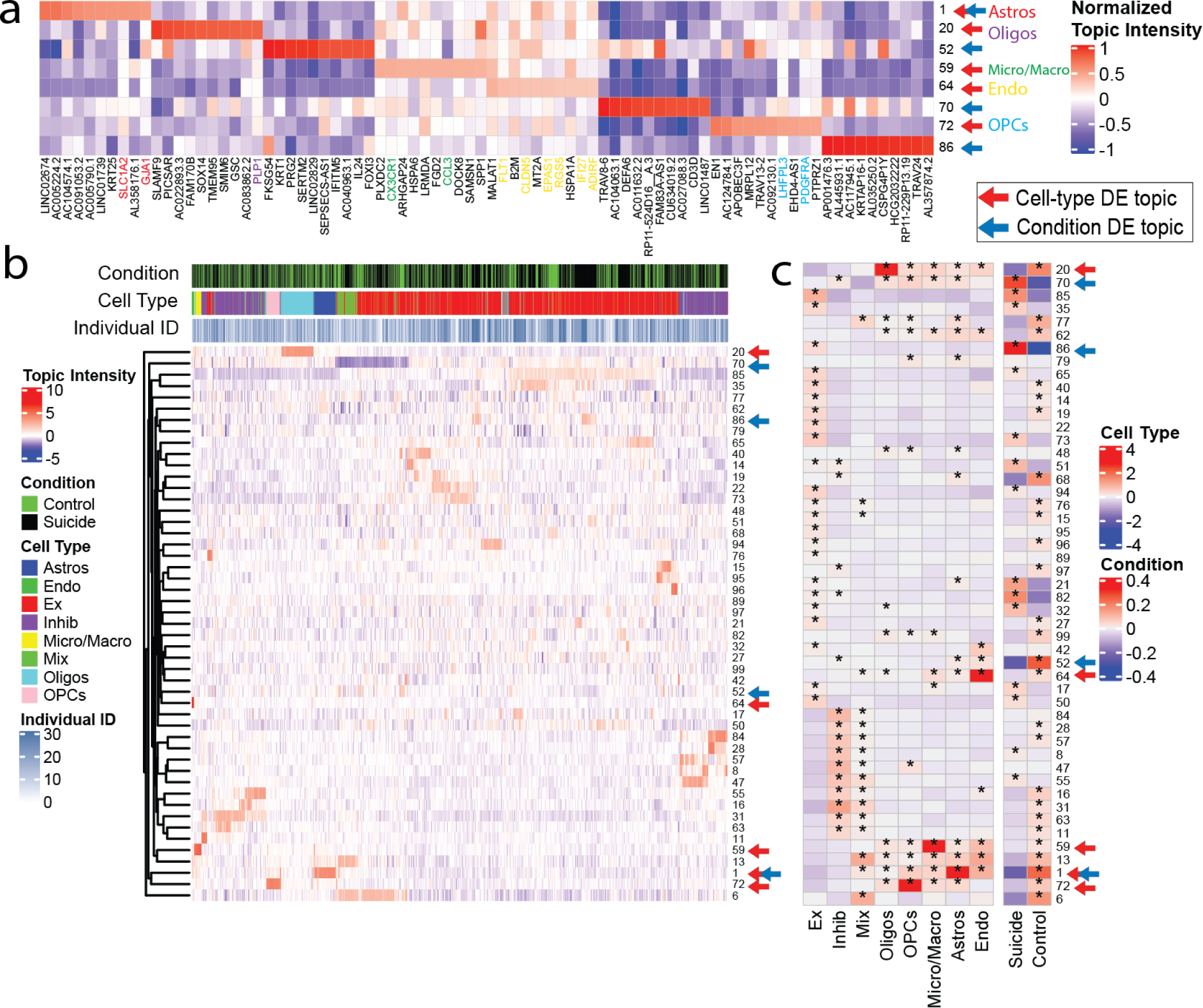
scETM-topic embeddings learned from the Major Depressive Disorder scRNA-seq data. **(a)** Gene-topics heatmap of top 10 genes in each topic based on topic in- tensity. The top genes which are known as cell-type marker genes based on PanglaoDB are highlighted (rows). For visualization purposes, we divided the topic values by the maximum absolute value within the same topic. Only cell-type and disease differential topics are shown. **(b)** Topics intensity of cells (n=10,000) sub-sampled from the MDD dataset. Topic intensities shown here are the Gaussian mean before applying softmax. Only the select topics with the sum of absolute values greater than 1500 across all sampled cells are shown. The three color bars show disease conditions, cell types, and batch identifiers (i.e., subject IDs). **(c)** Differential expression analysis of topics across the 8 cell types and 2 clinical conditions. Colors indicate mean differences between cell groups with and without a certain label (cell-type or condition). Asterisks indicate Bonferroni-corrected empirical p-value *<* 0.05 for permutation tests of up- regulated topics in each cell-type and disease labels. The number of permutations in each test is 100,000.

**Figure S19:**
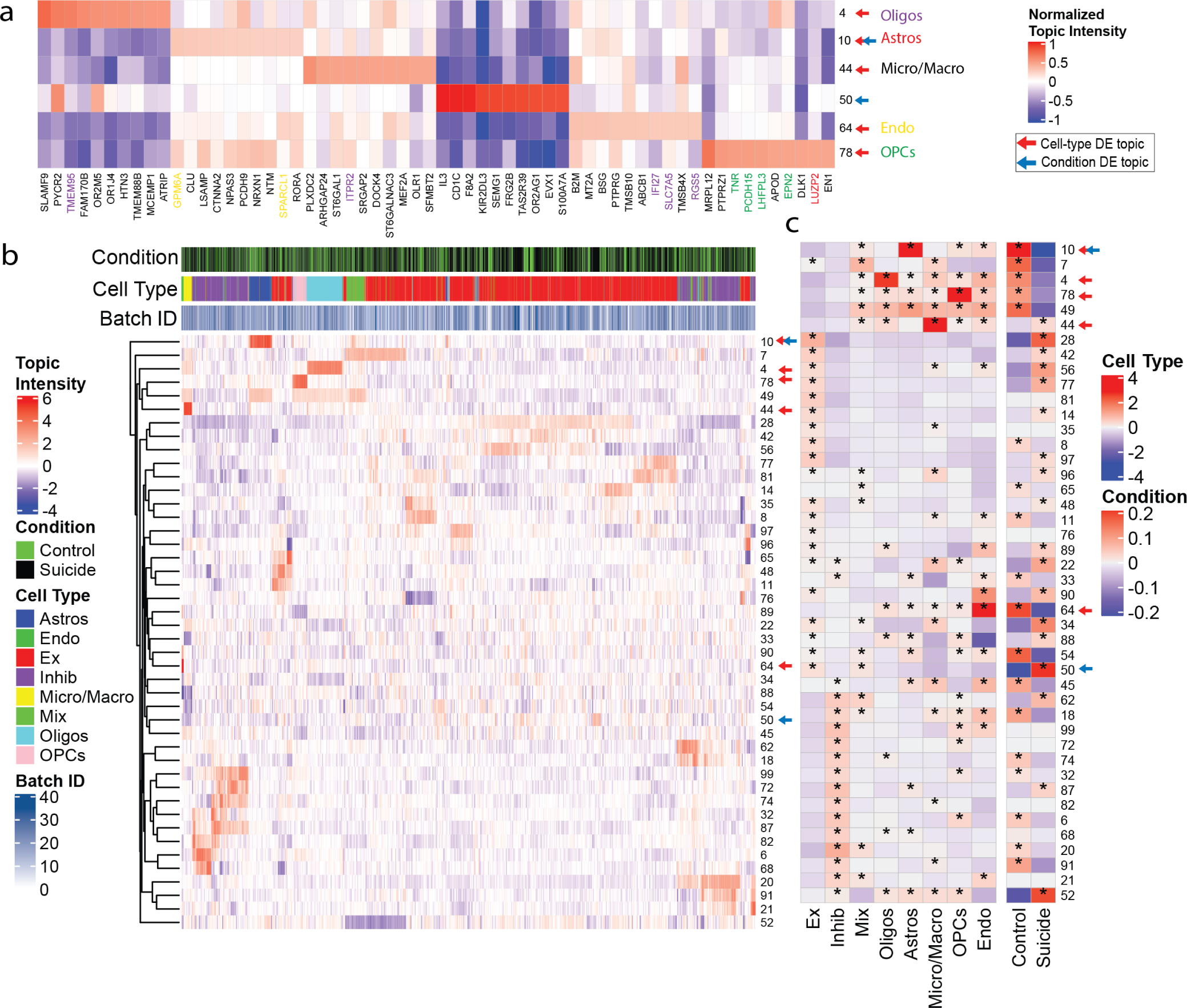
scETM-topic embeddings learned from the Major Depressive Disorder scRNA-seq data with genes restricted to protein coding genes. **(a)** Gene-topics heatmap of top 10 genes in each topic based on topic intensity. **(b)** Topics intensity of cells (n=10,000) sub-sampled from the MDD - coding genes only dataset. **(c)** Differential expression analysis of topics across the 8 cell types and 2 clinical conditions.

**Figure S20:**
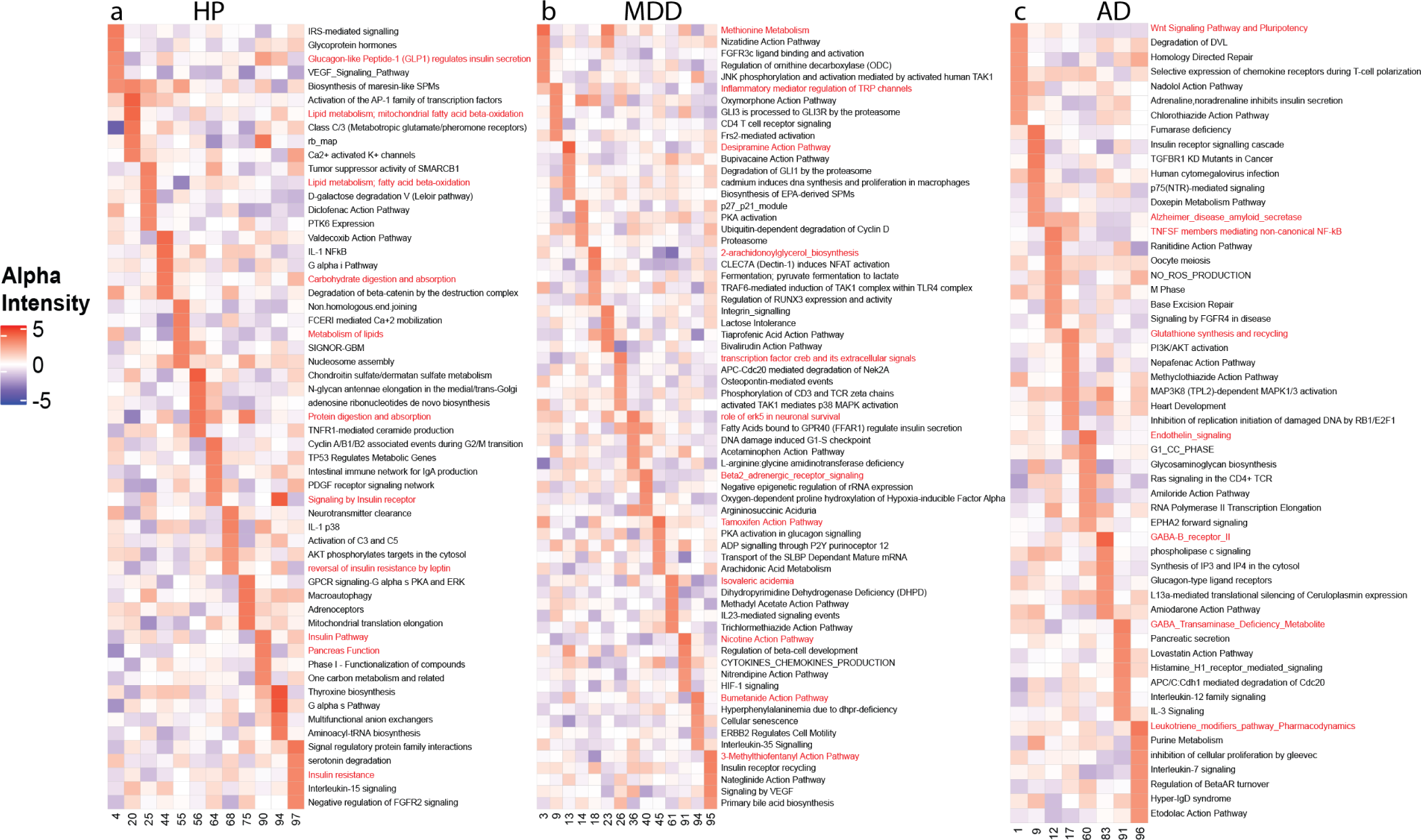
p-scETM pathway-topics embeddings by fixing pathDIP gene set database as *ρ*. **(a)** The pathway-topics heatmap of top 5 pathways in selected topics, inferred by a p-scETM model trained on HP. Pathways potentially related to pancreas function, insulin signalling and digestion are highlighted. **(b)** The pathway-topics heatmap of top 5 pathways in selected topics, inferred by a p-scETM model trained on MDD. Pathways potentially related to MDD pathogene- sis and therapeutic targets are highlighted. **(c)** The pathway-topics heatmap of top 7 pathways in selected topics, inferred by a p-scETM model trained on AD. Pathways potentially related to AD pathogenesis and therapeutic targets are highlighted.

## Supplementary Tables

**Table S1:** Normalized Mutual Information (NMI) between ground truth cell types and leiden clusters on 6 benchmark scRNA-seq datasets. NA is reported for models that did not converge. scETM performances with or without the linear batch correction (*scETM, scETM λ*) are both reported. scETM+adv is scETM plus adversarial network loss to further correct batch effects. Batch variables include strain, sequencing technologies (“Tech.”) and individuals (“Ind.”). NA is reported for models that did not converge. More details are described in Table 1 caption and **Methods**.

|  | MP | HP | TM | AD | MDD | MR |
| --- | --- | --- | --- | --- | --- | --- |
| Harmony | 0.928 | 0.927 | 0.827 | 0.983 | 0.705 | 0.726 |
| Scanorama | 0.845 | 0.816 | 0.766 | 0.988 | 0.561 | 0.747 |
| Seurat | 0.913 | 0.947 | 0.825 | 0.967 | 0.540 | 0.783 |
| scVAE-GM | 0.764 | NA | NA | 0.989 | 0.526 | 0.781 |
| scVI | 0.876 | 0.783 | 0.835 | 0.977 | 0.534 | 0.763 |
| LIGER | 0.875 | 0.876 | 0.757 | 0.861 | 0.618 | 0.742 |
| scVI-LD | 0.831 | 0.713 | 0.797 | 0.975 | 0.614 | 0.765 |
| scETM+adv | 0.852 | 0.908 | 0.838 | 0.985 | 0.598 | 0.747 |
| scETM | 0.902 | 0.902 | 0.856 | 0.987 | 0.583 | 0.809 |
| scETM$-\lambda$ | 0.819 | 0.644 | 0.819 | 0.987 | 0.604 | 0.764 |
| Batch Effect | Strain | Tech. | Tech. | Ind. | Ind. | Studies |

**Table S2:** Robustness analysis of the scETM model. Changing the encoder architecture, gene embedding dimensions and number of topics has limited impact on model performance. We report the average ARI and the held-out negative log-likehood (NLL) of three runs with differ- ent random seeds. The current model has an encoder with one 128-dim hidden layer, a gene embedding dimension of 400 and 50 topics.

|  | MP |  | HP |  |
| --- | --- | --- | --- | --- |
|  | ARI | NLL | ARI | NLL |
| Current model | 0.951 | 6.721 | 0.943 | 7.155 |
| Encoder arch (256, 128) | 0.897 | 6.741 | 0.940 | 7.151 |
| Encoder arch (512, 256, 128) | 0.898 | 6.722 | 0.936 | 7.159 |
| Gene emb. dim. 25 | 0.920 | 6.716 | 0.945 | 7.166 |
| Gene emb. dim. 100 | 0.916 | 6.734 | 0.944 | 7.155 |
| 10 topics | 0.941 | 6.723 | 0.940 | 7.168 |
| 200 topics | 0.915 | 6.771 | 0.944 | 7.152 |

**Table S3:** Ablation study of the scETM model. We report the average ARI of three repeated trails.

|  | MP |  | HP |  |
| --- | --- | --- | --- | --- |
|  | ARI | NLL | ARI | NLL |
| Current model | 0.951 | 6.721 | 0.943 | 7.155 |
| - BatchNorm | 0.844 | 6.724 | 0.834 | 7.149 |
| - Total count Normalization | 0.726 | $6.815 \times 10^4$ | 0.388 | $2.041 \times 10^6$ |
| - BatchNorm - Total count Normalization | 0.000 | $6.420 \times 10^4$ | 0.000 | $1.909 \times 10^6$ |
| - Batch correction module | 0.851 | 6.724 | 0.471 | 7.163 |

**Table S4:**
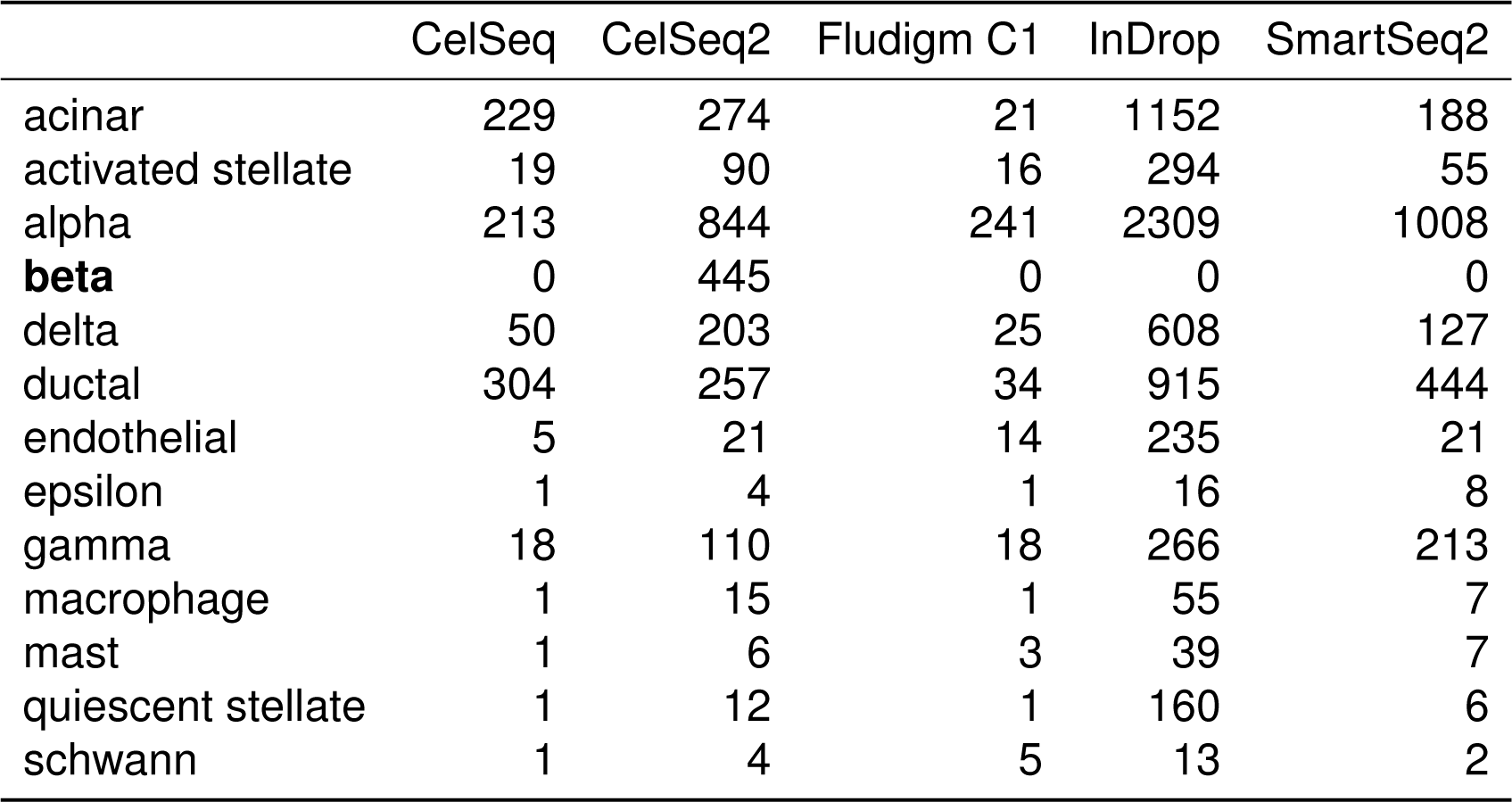
Cell type distribution of HP–beta. Beta cells are removed from all batches except CelSeq2 of the HP dataset.

|  | CelSeq | CelSeq2 | Fludigm C1 | InDrop | SmartSeq2 |
| --- | --- | --- | --- | --- | --- |
| acinar | 229 | 274 | 21 | 1152 | 188 |
| activated stellate | 19 | 90 | 16 | 294 | 55 |
| alpha | 213 | 844 | 241 | 2309 | 1008 |
| **beta** | 0 | 445 | 0 | 0 | 0 |
| delta | 50 | 203 | 25 | 608 | 127 |
| ductal | 304 | 257 | 34 | 915 | 444 |
| endothelial | 5 | 21 | 14 | 235 | 21 |
| epsilon | 1 | 4 | 1 | 16 | 8 |
| gamma | 18 | 110 | 18 | 266 | 213 |
| macrophage | 1 | 15 | 1 | 55 | 7 |
| mast | 1 | 6 | 3 | 39 | 7 |
| quiescent stellate | 1 | 12 | 1 | 160 | 6 |
| schwann | 1 | 4 | 5 | 13 | 2 |

**Table S5:** Batch overcorrection analysis on HP–beta. We report ARI, ASW, ASW-beta (SWs averaged over all beta cells) and kBET scores on the HP–beta dataset.

|  | ARI | ASW | ASW-beta | kBET |
| --- | --- | --- | --- | --- |
| Seurat v3 | 0.9225 | 0.2732 | 0.1490 | 0.3337 |
| Harmony | 0.9024 | 0.2323 | 0.1558 | 0.2637 |
| Scanorama | 0.8989 | 0.3832 | 0.3203 | 0.1067 |
| LIGER | 0.8476 | 0.1946 | 0.0912 | 0.5978 |
| scVI-LD | 0.5077 | 0.1397 | 0.6135 | 0.0031 |
| scVI | 0.6975 | 0.1579 | 0.4044 | 0.0567 |
| scETM+adv | 0.9265 | 0.3026 | 0.0045 | 0.3445 |
| scETM | 0.9298 | 0.3525 | 0.5370 | 0.1247 |

**Table S6:** Cell type distribution of MR.

|  | Shekhar *et al.* | Macosko *et al.* |
| --- | --- | --- |
| amacrine | 252 | 4426 |
| astrocytes | 0 | 54 |
| bipolar | 23494 | 6285 |
| cones | 48 | 1868 |
| fibroblasts | 0 | 85 |
| ganglion | 0 | 432 |
| horizontal | 0 | 252 |
| microglia | 0 | 67 |
| muller | 2945 | 1624 |
| pericytes | 0 | 63 |
| rods | 91 | 29400 |
| vascular endothelium | 0 | 252 |

**Table S7:** Batch overcorrection analysis on MR. We report ARI, ASW and kBET scores on the MR dataset.

|  | ARI | ASW | kBET |
| --- | --- | --- | --- |
| Harmony | 0.7632 | 0.1366 | 0.0631 |
| Scanorama | 0.7795 | 0.0934 | 0.0625 |
| Seurat v3 | 0.7813 | 0.3464 | 0.0534 |
| LIGER | 0.7135 | 0.2215 | 0.1761 |
| scVI | 0.7827 | 0.1331 | 0.0732 |
| scVI-LD | 0.7182 | 0.1964 | 0.0217 |
| scETM+adv | 0.7720 | 0.2669 | 0.1410 |
| scETM | 0.8593 | 0.2873 | 0.0656 |

**Table S8:** The impacts of doublets/contaminants on the model performance. ARI and kBET scores of each method, with and without filtering of the 669 doublets/contaminants in the Mouse Retina (MR) dataset, are reported.

| Dataset | w/ filtering |  | w/o filtering |  |
| --- | --- | --- | --- | --- |
| Metric | ARI | kBET | ARI | kBET |
| Harmony | 0.763 | 0.063 | 0.763 | 0.044 |
| Scanorama | 0.780 | 0.063 | 0.770 | 0.052 |
| Seurat v3 | 0.781 | 0.053 | 0.638 | 0.007 |
| scVI | 0.783 | 0.073 | 0.774 | 0.060 |
| LIGER | 0.714 | 0.176 | 0.463 | 0.196 |
| scVI-LD | 0.718 | 0.022 | 0.701 | 0.013 |
| scETM+adv | 0.772 | 0.141 | 0.767 | 0.129 |
| scETM | 0.859 | 0.066 | 0.854 | 0.057 |

**Table S9:**
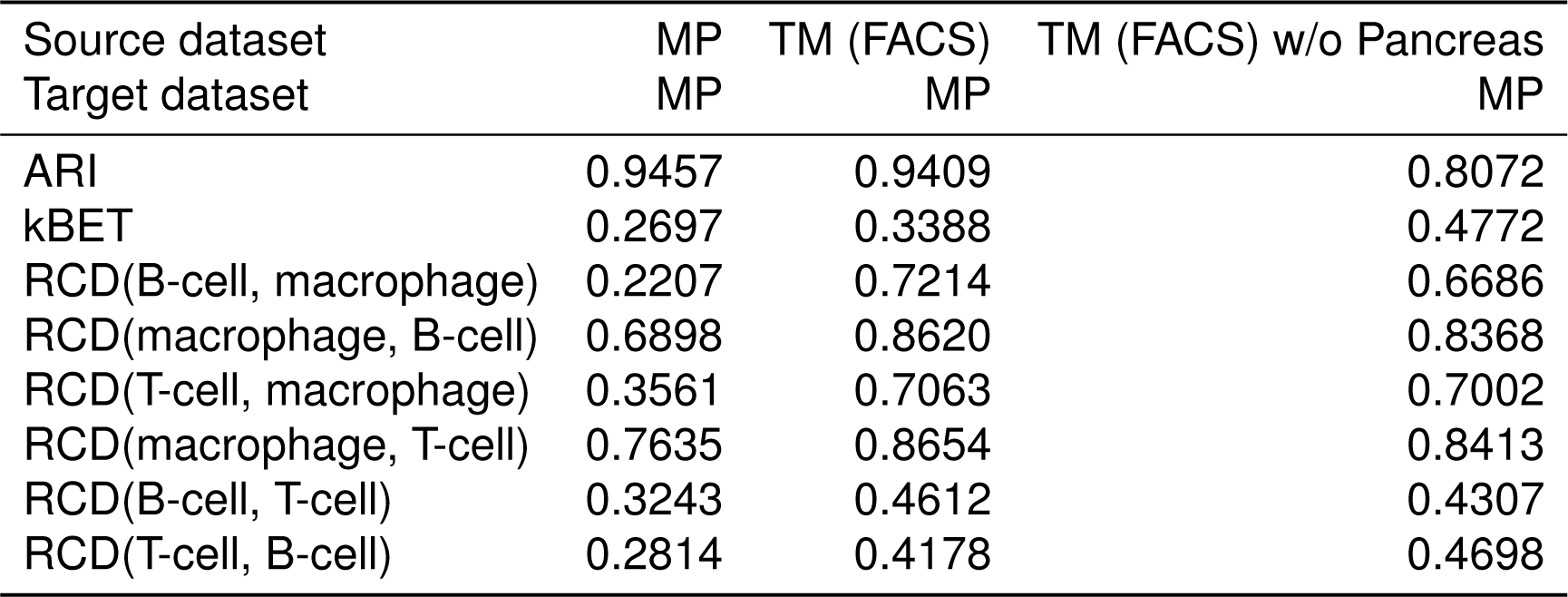
Separation of immune cell types in MP with transfer learning on TM (FACS). We separately trained scETM on MP, TM (FACS) without Pancreas and TM (FACS), and then applied them to the MP dataset to cluster MP cells. We reported the Relative Clus- ter Distance (RCD) between different immune cell types (T-cell and B-cell) in MP, as well as the overall ARI and kBET scores. RCD is inspired by silhouette width and is defined by 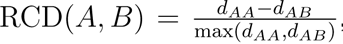, where A, B are two cell types and *d* is the cosine distance. Higher values indicate cell types A and B are more separated in the embedding. All results are averages over three runs with different random seeds.

**Table S10:** Clustering performance on target datasets in the 6 cross-tissue and cross- species transfer learning tasks. The clustering performance is measured by Adjusted Rand Index (ARI) between ground truth cell types and Leiden [75] clusters. The cell clusters based on the cell embedding produced by each method are also depicted in the UMAP visualization in **Fig. 4**. Abbreviations: MP: Mouse Pancreas; HP (InDrop): Human Pancreas sequenced with InDrop technology; TM (FACS) : A subset of Tabula Muris sequenced with Fluorescence- Activated Single Cell Sorting; TM (FACS) w/o Panc: TM-FACS without Pancreas; HumM1C: human primary motor cortex (from Allen Brain map); MusMOp: Mouse primary motor area (from Allen Brain map).

| Type of transfer | Cross-tissue |  |  |  | Cross-species |  |  |
| --- | --- | --- | --- | --- | --- | --- | --- |
| Source dataset | MP | TM (FACS) | TM (FACS) w/o Panc | MP | HP (InDrop) | MusMOp | HumM1C |
| Target dataset | TM (FACS) | MP | MP | HP (InDrop) | MP | HumM1C | MusMOp |
| scVI | 0.5075 | 0.4844 | 0.4197 | 0.5236 | 0.4251 | 0.0900 | 0.0252 |
| scVI-LD | 0.5159 | 0.3985 | 0.4317 | 0.6902 | 0.4757 | 0.0895 | 0.0375 |
| scETM | 0.5659 | 0.9409 | 0.8072 | 0.8680 | 0.7998 | 0.7105 | 0.3515 |

**Table S11:** kBET scores on target datasets during cross-tissue and cross-species trans- fer learning. Abbreviations are the same as in **Supp.** Table S10

| Type of transfer | Cross-tissue |  |  |  | Cross-species |  |  |
| --- | --- | --- | --- | --- | --- | --- | --- |
| Source dataset | MP | TM (FACS) | TM (FACS) w/o Panc | MP | HP (InDrop) | MusMOp | HumM1C |
| Target dataset | TM (FACS) | MP | MP | HP (InDrop) | MP | HumM1C | MusMOp |
| scVI | 0.0874 | 0.2570 | 0.2672 | 0.1707 | 0.2276 | 0.9002 | 0.8478 |
| scVI-LD | 0.0559 | 0.2564 | 0.2948 | 0.1425 | 0.2204 | 0.9181 | 0.8524 |
| scETM | 0.0585 | 0.3388 | 0.4772 | 0.1782 | 0.2930 | 0.7391 | 0.7775 |

**Table S12:** Pathway enrichment statistics for Human Pancreas dataset. Pathways whose names include the keywords “insulin” or “pancreatic” are shown. For each pathway, the number of topics with significant enrichment (BH-corrected q-value < 0.01) are counted. Both scETM and scVI-LD use latent dimensions of 100 for fair comparison.

| Pathway Name | scETM | scVI-LD |
| --- | --- | --- |
| REACTOME_INSULIN_PROCESSING | 15 | 2 |
| REACTOME_INSULIN_RECEPTOR_RECYCLING | 6 | 0 |
| PID_INSULIN_GLUCOSE_PATHWAY | 4 | 0 |
| WP_PANCREATIC_ADENOCARCINOMA_PATHWAY | 2 | 0 |
| KEGG_PANCREATIC_CANCER | 2 | 0 |
| PID_INSULIN_PATHWAY | 2 | 0 |
| SIG_INSULIN_RECEPTOR_PATHWAY_IN_CARDIAC_MYOCYTES | 1 | 0 |
| REACTOME_REGULATION_OF_GENE_EXPRESSION_IN_LATE_STAGE... | 1 | 0 |
| REACTOME_REGULATION_OF_INSULIN_SECRETION | 1 | 1 |
| REACTOME_REGULATION_OF_INSULIN_LIKE_GROWTH_FACTOR... | 1 | 3 |
| WP_INSULIN_SIGNALING | 1 | 0 |
| BIOCARTA_INSULIN_PATHWAY | 1 | 0 |
| REACTOME_INSULIN_RECEPTOR_SIGNALLING_CASCADE | 0 | 1 |
| REACTOME_SYNTHESIS_SECRETION_AND_INACTIVATION_OF_GLUCOSE... | 0 | 1 |
| REACTOME_SIGNALING_BY_TYPE_1_INSULIN_LIKE_GROWTH_FACTOR... | 0 | 1 |
| WP_FACTORS_AND_PATHWAYS_AFFECTING_INSULINLIKE_GROWTH_FACTOR... | 0 | 1 |
| **Total number of pathways** | **12** | **7** |

**Table S13:** Pathway enrichment statistics for AD dataset. Pathways whose names include the keywords “amyloid” or “alzheimer” are shown. For each pathway, the number of topics with significant enrichment (BH-corrected q-value < 0.01) are counted. Both scETM and scVI-LD use latent dimensions of 100 for fair comparison.

| Pathway Name | scETM | scVI-LD |
| --- | --- | --- |
| REACTOME_AMYLOID_FIBER_FORMATION | 1 | 2 |
| KEGG_ALZHEIMERS_DISEASE | 1 | 2 |
| REACTOME_DEREGULATED_CDK5_TRIGGERS_MULTIPLE_NEURODEGENERATIVE_PATHWAYS_IN_ALZHEIMER_S_DISEASE_MODELS | 1 | 0 |
| **Total number of pathways** | **3** | **2** |

**Table S14:**
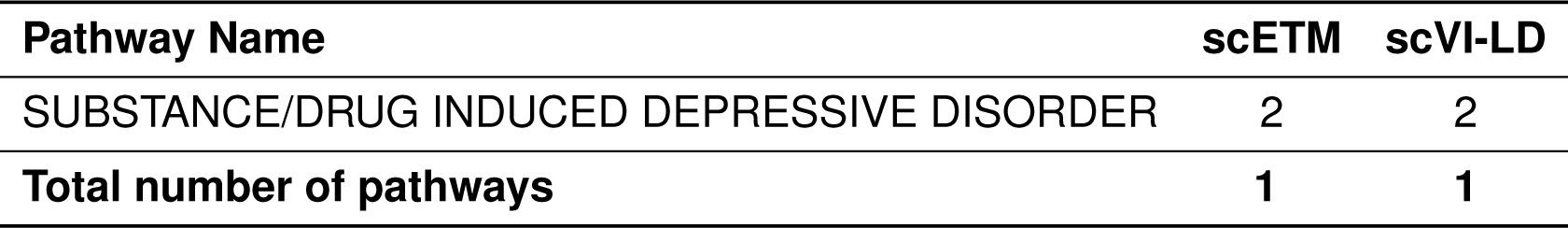
Pathway enrichment statistics for MDD dataset. Pathways whose names include the keyword “depressive” are shown. For each pathway, the number of topics with significant enrichment (BH-corrected q-value < 0.01) are counted. Both scETM and scVI-LD use latent dimensions of 100 for fair comparison.

| Pathway Name | scETM | scVI-LD |
| --- | --- | --- |
| SUBSTANCE/DRUG INDUCED DEPRESSIVE DISORDER | 2 | 2 |
| **Total number of pathways** | **1** | **1** |

**Table S15:** Adjusted Rand Index (ARI) comparison of scETMs and p-scETMs in three human single cell transcriptomics datasets. Refer to **Incorporation of pathway knowledge into the gene embeddings in p-scETM** section for experimental details.

|  | HP | MDD | AD |
| --- | --- | --- | --- |
| scETM | 0.937 | 0.734 | 0.976 |
| p-scETM | 0.926 | 0.753 | 0.996 |

**Table S16:** MDD-relevant pathways from the pathway-topic embedding inferred by p-scETM trained on the MDD dataset.

| Pathway | Relevance | Reference |
| --- | --- | --- |
| Methionine Metabolism | MDD treatment | [80] |
| Inflammatory mediator regulation of TRP channels | MDD treatment | [81] |
| Desipramine Action Pathway | MDD treatment | [82, 83] |
| 2-arachidonoylglycerol_biosynthesis | MDD pathogenesis | [84] |
| transcription factor creb and its extracellular signals | MDD treatment | [85, 86] |
| role of erk5 in neuronal survival | MDD pathogenesis and treatment | [87] |
| Beta2_adrenergic_receptor_signaling | same pathway enrichment found in a MDD study | [50] |
| Tamoxifen Action Pathway | correlation with MDD onset | [88] |
| Isovaleric acidemia | correlation with MDD | [89] |
| Nicotine Action Pathway | correlation with MDD onset | [90] |
| Bumetanide Action Pathway | MDD treatment | [91] |
| 3-Methylthiofentanyl Action Pathway | MDD treatment | [92] |

**Table S17:** AD-relevant pathways from the pathway-topic embedding inferred by p-scETM trained on the AD dataset.

| Pathway | Relevance | Reference |
| --- | --- | --- |
| Wnt Signaling Pathway and Pluripotency | AD treatment | [93] |
| Alzheimer_disease_amyloid_secretase | AD pathogenesis | [94] |
| TNF receptor superfamily (TNFS) members mediating non-canonical NF-kB | AD treatment | [95] |
| Glutathione synthesis and recycling | AD pathogenesis | [96] |
| Endothelin_signaling | AD treatment | [97] |
| GABA-B_receptor_II | AD treatment | [98, 99] |
| GABA_Transaminase_Deficiency_Metabolite | AD treatment | [98, 99] |
| Leukotriene_modifiers_pathway_Pharmacodynamics | AD treatment | [100] |

Table S18: Pathway enrichment statistics breakdown of scETM for four datasets including HP, AD, MDD (all genes) and MDD (coding genes only), and scVI-LD for three datasets including HP, AD, MDD (all genes). The table is saved in Additional file 1.xlsx.

Table S19: Top 5 pathways of each topic (row) based on alpha intensities for four datasets including HP, AD MDD (all genes), MDD (coding genes only). The table is saved in Additional file 2.xlsx.

**Table S20:** Differential expression (DE) analysis summary of topics in AD and MDD data (up- regulation only).

|  | AD | MDD |
| --- | --- | --- |
| DE cell-type and topics pairs | 58 | 110 |
| Cell types identified by DE topics | 8 | 8 |
| DE topics w.r.t. disease | 8 | 15 |
| Topics associated with cell types and disease | 8 | 15 |

**Table S21:** Performance of scVI(-LD) with 10 and 100 latent dimensions on the five benchmark scRNA-seq datasets. The results showed that scVI and scVI-LD with 10 dimensions performed better than the same models with 100 dimensions, justifying our choice of 10 latent dimensions when comparing these models with other models.

| Metric | Method | MP | HP | MDD | AD | TM |
| --- | --- | --- | --- | --- | --- | --- |
| ARI | scVI-LD (10) | **0.8753** | **0.6563** | **0.6401** | **0.9889** | 0.6076 |
|  | scVI-LD (100) | 0.4454 | 0.4431 | 0.6145 | 0.9848 | **0.6705** |
|  | scVI (10) | **0.9325** | **0.7590** | 0.5411 | **0.9915** | **0.6699** |
|  | scVI (100) | 0.8215 | 0.7051 | **0.5813** | 0.9921 | 0.6284 |
| kBET | scVI-LD (10) | 0.1483 | **0.0335** | **0.2759** | **0.3967** | **0.0687** |
|  | scVI-LD (100) | **0.4601** | 0.0045 | 0.2745 | 0.3570 | 0.0331 |
|  | scVI (10) | 0.5159 | **0.1403** | 0.3132 | 0.4338 | **0.0557** |
|  | scVI (100) | **0.6856** | 0.0876 | **0.3402** | **0.4371** | 0.0301 |

## References

1. Byungjin Hwang, Ji Hyun Lee, and Duhee Bang. Single-cell rna sequencing technologies and bioinformatics pipelines. Experimental & molecular medicine, 50(8):1–14, 2018.

2. Xiaoping Han, Ziming Zhou, Lijiang Fei, Huiyu Sun, Renying Wang, Yao Chen, Haide Chen, Jingjing Wang, Huanna Tang, Wenhao Ge, et al. Construction of a human cell landscape at single-cell level. Nature, 581(7808):303–309, 2020.

3. Tabula Muris Consortium et al. Single-cell transcriptomics of 20 mouse organs creates a tabula muris. Nature, 562(7727):367, 2018.

4. Aviv Regev, Sarah A Teichmann, Eric S Lander, Ido Amit, Christophe Benoist, Ewan Birney, Bernd Bodenmiller, Peter Campbell, Piero Carninci, Menna Clatworthy, et al. Science forum: the human cell atlas. Elife, 6:e27041, 2017.

5. Orit Rozenblatt-Rosen, Michael JT Stubbington, Aviv Regev, and Sarah A Teichmann. The human cell atlas: from vision to reality. Nature News, 550(7677):451, 2017.

6. Romain Lopez, Jeffrey Regier, Michael B. Cole, Michael I. Jordan, and Nir Yosef. Deep generative modeling for single-cell transcriptomics. Nature Methods, 15(12):1053–1058, 2018.

7. Tim Stuart, Andrew Butler, Paul Hoffman, Christoph Hafemeister, Efthymia Papalexi, William M Mauck III, Yuhan Hao, Marlon Stoeckius, Peter Smibert, and Rahul Satija. Comprehensive integration of single-cell data. Cell, 177(7):1888–1902, 2019.

8. Andrew Butler, Paul Hoffman, Peter Smibert, Efthymia Papalexi, and Rahul Satija. Integrating single-cell transcriptomic data across different conditions, technologies, and species. Nature biotechnology, 36(5):411–420, 2018.

9. Joshua D Welch, Velina Kozareva, Ashley Ferreira, Charles Vanderburg, Carly Martin, and Evan Z Macosko. Single-cell multi-omic integration compares and contrasts features of brain cell identity. Cell, 177(7):1873–1887, 2019.

10. Brian Hie, Bryan Bryson, and Bonnie Berger. Efficient integration of heterogeneous single-cell transcriptomes using scanorama. Nature biotechnology, 37(6):685–691, 2019.

11. Christopher H Grønbech, Maximillian F Vording, Pascal N Timshel, Capser K Sønderby, Tune H Pers, and Ole Winther. scvae: Variational auto-encoders for single-cell gene expression datas. bioRxiv, page 318295, 2018.

12. Laleh Haghverdi, Aaron TL Lun, Michael D Morgan, and John C Marioni. Batch effects in single-cell rna-sequencing data are corrected by matching mutual nearest neighbors. Nature biotechnology, 36(5):421–427, 2018.

13. Zhe Sun, Li Chen, Hongyi Xin, Yale Jiang, Qianhui Huang, Anthony R Cillo, Tracy Tabib, Jay K Kolls, Tullia C Bruno, Robert Lafyatis, et al. A bayesian mixture model for clustering droplet-based single-cell transcriptomic data from population studies. Nature communications, 10(1):1–10, 2019.

14. Valentine Svensson, Adam Gayoso, Nir Yosef, and Lior Pachter. Interpretable factor models of single-cell rna-seq via variational autoencoders. Bioinformatics, 36(11):3418– 3421, 2020.

15. Nelson Johansen and Gerald Quon. scalign: a tool for alignment, integration, and rare cell identification from scrna-seq data. Genome biology, 20(1):1–21, 2019.

16. F Alexander Wolf, Philipp Angerer, and Fabian J Theis. Scanpy: large-scale single-cell gene expression data analysis. Genome biology, 19(1):15, 2018.

17. Peng Qiu. Embracing the dropouts in single-cell rna-seq analysis. Nature communications, 11(1):1–9, 2020.

18. Maren Büttner, Zhichao Miao, F Alexander Wolf, Sarah A Teichmann, and Fabian J Theis. A test metric for assessing single-cell rna-seq batch correction. Nature methods, 16(1):43–49, 2019.

19. Po-Yuan Tung, John D Blischak, Chiaowen Joyce Hsiao, David A Knowles, Jonathan E Burnett, Jonathan K Pritchard, and Yoav Gilad. Batch effects and the effective design of single-cell gene expression studies. Scientific reports, 7:39921, 2017.

20. Vladimir Yu Kiselev, Tallulah S Andrews, and Martin Hemberg. Challenges in unsupervised clustering of single-cell rna-seq data. Nature Reviews Genetics, 20(5):273–282, 2019.

21. Daniel Backenroth, Zihuai He, Krzysztof Kiryluk, Valentina Boeva, Lynn Pethukova, Ekta Khurana, Angela Christiano, Joseph D Buxbaum, and Iuliana Ionita-Laza. FUN-LDA: A Latent Dirichlet Allocation Model for Predicting Tissue-Specific Functional Effects of Noncoding Variation: Methods and Applications. The American Journal of Human Genetics, 102(5):920–942, 2018.

22. Carmen Bravo González-Blas, Liesbeth Minnoye, Dafni Papasokrati, Sara Aibar, Gert Hulselmans, Valerie Christiaens, Kristofer Davie, Jasper Wouters, and Stein Aerts. cisTopic: cis-regulatory topic modeling on single-cell ATAC-seq data. Nature Methods, 16(5):1–14, April 2019.

23. Yue Li, Pratheeksha Nair, Xing Han Lu, Zhi Wen, Yuening Wang, Amir Ardalan Kalantari Dehaghi, Yan Miao, Weiqi Liu, Tamas Ordog, Joanna M Biernacka, Euijung Ryu, Janet E Olson, Mark A Frye, Aihua Liu, Liming Guo, Ariane Marelli, Yuri Ahuja, Jose Davila-Velderrain, and Manolis Kellis. Inferring multimodal latent topics from electronic health records. Nature Communications, 11(1):1 – 17, 05 2020.

24. Zhe Wang, Shiyi Yang, Yusuke Koga, Sean E. Corbett, W. Evan Johnson, Masanao Yajima, and Joshua D. Campbell. Celda: A Bayesian model to perform co-clustering of genes into modules and cells into subpopulations using single-cell RNA-seq data. bioRxiv, page 2020.11.16.373274, 2021.

25. D D Lee and H S Seung. Learning the parts of objects by non-negative matrix factoriza- tion. Nature, 401(6755):788–791, 10 1999.

26. Mohammad Lotfollahi, F. Alexander Wolf, and Fabian J. Theis. scgen predicts single-cell perturbation responses. Nature Methods, 16(8):715–721, Aug 2019.

27. Mohammad Lotfollahi, Mohsen Naghipourfar, Malte D. Luecken, Matin Khajavi, Maren Büttner, Ziga Avsec, Alexander V. Misharin, and Fabian J. Theis. Query to reference single-cell integration with transfer learning. bioRxiv, 2020.

28. Ilya Korsunsky, Nghia Millard, Jean Fan, Kamil Slowikowski, Fan Zhang, Kevin Wei, Yuriy Baglaenko, Michael Brenner, Po-ru Loh, and Soumya Raychaudhuri. Fast, sensitive and accurate integration of single-cell data with harmony. Nature methods, pages 1–8, 2019.

29. Corina Nagy, Malosree Maitra, Arnaud Tanti, Matthew Suderman, Jean-Francois Théroux, Maria Antonietta Davoli, Kelly Perlman, Volodymyr Yerko, Yu Chang Wang, Shreejoy J Tripathy, et al. Single-nucleus transcriptomics of the prefrontal cortex in major depressive disorder implicates oligodendrocyte precursor cells and excitatory neurons. Nature Neuroscience, pages 1–11, 2020.

30. Sumit Mukherjee, Yue Zhang, Joshua Fan, Georg Seelig, and Sreeram Kannan. Scalable preprocessing for sparse scRNA-seq data exploiting prior knowledge. Bioinformatics, 34(13):i124–i132, 2018.

31. Maria Brbic, Marinka Zitnik, Sheng Wang, Angela O Pisco, Russ B Altman, Spyros Darmanis, and Jure Leskovec. Mars: discovering novel cell types across heterogeneous single-cell experiments. Nature Methods, pages 1–7, 2020.

32. Diederik P Kingma and Max Welling. Auto-Encoding Variational Bayes. arXiv.org, December 2013.

33. Adji B. Dieng, Francisco J. R. Ruiz, and David M. Blei. Topic modeling in embedding spaces. Transactions of the Association for Computational Linguistics, 8:439–453, 2020.

34. Maayan Baron, Adrian Veres, Samuel L Wolock, Aubrey L Faust, Renaud Gaujoux, Amedeo Vetere, Jennifer Hyoje Ryu, Bridget K Wagner, Shai S Shen-Orr, Allon M Klein, et al. A single-cell transcriptomic map of the human and mouse pancreas reveals inter- and intra-cell population structure. Cell systems, 3(4):346–360, 2016.

35. Hansruedi Mathys, Jose Davila-Velderrain, Zhuyu Peng, Fan Gao, Shahin Mohammadi, Jennie Z Young, Madhvi Menon, Liang He, Fatema Abdurrob, Xueqiao Jiang, et al. Single-cell transcriptomic analysis of alzheimer’s disease. Nature, 570(7761):332–337, 2019.

36. Evan Z. Macosko, Anindita Basu, Rahul Satija, James Nemesh, Karthik Shekhar, Melissa Goldman, Itay Tirosh, Allison R. Bialas, Nolan Kamitaki, Emily M. Martersteck, John J. Trombetta, David A. Weitz, Joshua R. Sanes, Alex K. Shalek, Aviv Regev, and Steven A. McCarroll. Highly Parallel Genome-wide Expression Profiling of Individual Cells Using Nanoliter Droplets. Cell, 161(5):1202–1214, 2015.

37. Karthik Shekhar, Sylvain W. Lapan, Irene E. Whitney, Nicholas M. Tran, Evan Z. Macosko, Monika Kowalczyk, Xian Adiconis, Joshua Z. Levin, James Nemesh, Melissa Goldman, Steven A. McCarroll, Constance L. Cepko, Aviv Regev, and Joshua R. Sanes. Comprehensive Classification of Retinal Bipolar Neurons by Single-Cell Transcrip- tomics. Cell, 166(5):1308–1323.e30, 2016.

38. Mojtaba Bahrami, Malosree Maitra, Corina Nagy, Gustavo Turecki, Hamid R Rabiee, and Yue Li. Deep feature extraction of single-cell transcriptomes by generative adver- sarial network. *Bioinformatics (Oxford*, England), 3:346, 2020. btaa976.

39. L. McInnes, J. Healy, and J. Melville. UMAP: Uniform Manifold Approximation and Projection for Dimension Reduction. *ArXiv e-prints*, February 2018.

40. Fatima Batool and Christian Hennig. Clustering with the Average Silhouette Width. Computational Statistics & Data Analysis, 158:107190, 2021.

41. R Ranganath, S Gerrish, D Blei Artificial Intelligence Statistics,, and 2014. Black box variational inference. jmlr.org.

42. M D Hoffman, D M Blei, C Wang, and J W Paisley. Stochastic variational inference. Journal of Machine Learning Research (JMLR), 2013.

43. Zizhen Yao, Thuc Nghi Nguyen, Cindy T. J. van Velthoven, Jeff Goldy, Adriana E. Sedeno-Cortes, Fahimeh Baftizadeh, Darren Bertagnolli, Tamara Casper, Kirsten Crich- ton, Song-Lin Ding, Olivia Fong, Emma Garren, Alexandra Glandon, James Gray, Lucas T. Graybuck, Daniel Hirschstein, Matthew Kroll, Kanan Lathia, Boaz Levi, Delissa McMillen, Stephanie Mok, Thanh Pham, Qingzhong Ren, Christine Rimorin, Nadiya Shapovalova, Josef Sulc, Susan M. Sunkin, Michael Tieu, Amy Torkelson, Herman Tung, Katelyn Ward, Nick Dee, Kimberly A. Smith, Bosiljka Tasic, and Hongkui Zeng. A taxonomy of transcriptomic cell types across the isocortex and hippocampal formation. bioRxiv, page 2020.03.30.015214, 03 2020.

44. Aravind Subramanian, Pablo Tamayo, Vamsi K Mootha, Sayan Mukherjee, Benjamin L Ebert, Michael A Gillette, Amanda Paulovich, Scott L Pomeroy, Todd R Golub, Eric S Lander, and Jill P Mesirov. Gene set enrichment analysis: a knowledge-based approach for interpreting genome-wide expression profiles. Proceedings of the National Academy of Sciences of the United States of America, 102(43):15545 – 15550, 10 2005.

45. Xi Chen and Shi Du Yan. Mitochondrial a*β* a potential cause of metabolic dysfunction in alzheimer’s disease. IUBMB life, 58(12):686–694, 2006.

46. Moran N Cabili, Cole Trapnell, Loyal Goff, Magdalena Koziol, Barbara Tazon-Vega, Aviv Regev, and John L Rinn. Integrative annotation of human large intergenic noncod- ing rnas reveals global properties and specific subclasses. Genes & development, 25(18):1915–1927, 2011.

47. Elena Perenthaler, Soheil Yousefi, Eva Niggl, and Stefan Barakat. Beyond the exome: the non-coding genome and enhancers in malformations of cortical development. Fron- tiers in cellular neuroscience, 13:352, 2019.

48. Sara Rahmati, Mark Abovsky, Chiara Pastrello, Max Kotlyar, Richard Lu, Christian A Cumbaa, Proton Rahman, Vinod Chandran, and Igor Jurisica. pathdip 4: an extended pathway annotations and enrichment analysis resource for human, model organisms and domesticated species. Nucleic acids research, 48(D1):D479–D488, 2020.

49. pathdip 4.0 database. http://ophid.utoronto.ca/pathDIP/Download.jsp. accessed 23 oct 2020.

50. Anqi Qiu, Mojun Shen, Claudia Buss, Yap-Seng Chong, Kenneth Kwek, Seang-Mei Saw, Peter D Gluckman, Pathik D Wadhwa, Sonja Entringer, Martin Styner, et al. Effects of antenatal maternal depressive symptoms and socio-economic status on neonatal brain development are modulated by genetic risk. Cerebral Cortex, 27(5):3080–3092, 2017.

51. Michael Ashburner, Catherine A Ball, Judith A Blake, David Botstein, Heather Butler, J Michael Cherry, Allan P Davis, Kara Dolinski, Selina S Dwight, Janan T Eppig, et al. Gene ontology: tool for the unification of biology. Nature genetics, 25(1):25–29, 2000.

52. The gene ontology resource: enriching a gold mine. Nucleic Acids Research, 49(D1):D325–D334, 2021.

53. Ioannis Mantas, Marcus Saarinen, Zhi-Qing David Xu, and Per Svenningsson. Update on gpcr-based targets for the development of novel antidepressants. Molecular Psychiatry, pages 1–25, 2021.

54. Hanna Mendes Levitin, Jinzhou Yuan, Yim Ling Cheng, Francisco JR Ruiz, Erin C Bush, Jeffrey N Bruce, Peter Canoll, Antonio Iavarone, Anna Lasorella, David M Blei, et al. De novo gene signature identification from single-cell rna-seq with hierarchical poisson factorization. Molecular systems biology, 15(2):e8557, 2019.

55. Adam Gayoso, Zoë Steier, Romain Lopez, Jeffrey Regier, Kristopher L Nazor, Aaron Streets, and Nir Yosef. Joint probabilistic modeling of single-cell multi-omic data with totalVI. Nature Methods, pages 1 – 31, 03 2021.

56. Quoc Le and Tomas Mikolov. Distributed representations of sentences and documents. In International conference on machine learning, pages 1188–1196. PMLR, 2014.

57. Jingcheng Du, Peilin Jia, Yulin Dai, Cui Tao, Zhongming Zhao, and Degui Zhi. Gene2vec: distributed representation of genes based on co-expression. BMC Genomics, 20(Suppl 1):82, 02 2019.

58. Jian Tang, Meng Qu, Mingzhe Wang, Ming Zhang, Jun Yan, and Qiaozhu Mei. Line: Large-scale information network embedding. In Proceedings of the 24th international conference on world wide web, pages 1067–1077, 2015.

59. Aditya Grover and Jure Leskovec. node2vec: Scalable feature learning for networks. In Proceedings of the 22nd ACM SIGKDD international conference on Knowledge discovery and data mining, pages 855–864, 2016.

60. David M Blei, Andrew Y Ng, and Michael I Jordan. Latent dirichlet allocation. The Journal of Machine Learning Research, 3:993–1022, March 2003.

61. Xavier Glorot, Antoine Bordes, and Yoshua Bengio. Deep sparse rectifier neural networks. volume 15 of *Proceedings of Machine Learning Research*, pages 315–323, Fort Lauderdale, FL, USA, 11–13 Apr 2011. JMLR Workshop and Conference Proceedings.

62. Sergey Ioffe and Christian Szegedy. Batch normalization: Accelerating deep network training by reducing internal covariate shift. In Proceedings of the 32nd International Conference on International Conference on Machine Learning - Volume 37, ICML’15, pages 448–456. JMLR.org, 2015.

63. Adam Paszke, Sam Gross, Francisco Massa, Adam Lerer, James Bradbury, Gregory Chanan, Trevor Killeen, Zeming Lin, Natalia Gimelshein, Luca Antiga, Alban Desmai- son, Andreas Kopf, Edward Yang, Zachary DeVito, Martin Raison, Alykhan Tejani, Sasank Chilamkurthy, Benoit Steiner, Lu Fang, Junjie Bai, and Soumith Chintala. Pytorch: An imperative style, high-performance deep learning library. In H. Wallach, H. Larochelle, A. Beygelzimer, F. d’Alché-Buc, E. Fox, and R. Garnett, editors, Advances in Neural Information Processing Systems 32, pages 8024–8035. Curran Associates, Inc., 2019.

64. Martín Abadi, Ashish Agarwal, Paul Barham, Eugene Brevdo, Zhifeng Chen, Craig Citro, Greg S. Corrado, Andy Davis, Jeffrey Dean, Matthieu Devin, Sanjay Ghemawat, Ian Goodfellow, Andrew Harp, Geoffrey Irving, Michael Isard, Yangqing Jia, Rafal Joze- fowicz, Lukasz Kaiser, Manjunath Kudlur, Josh Levenberg, Dandelion Mané, Rajat Monga, Sherry Moore, Derek Murray, Chris Olah, Mike Schuster, Jonathon Shlens, Benoit Steiner, Ilya Sutskever, Kunal Talwar, Paul Tucker, Vincent Vanhoucke, Vijay Vasudevan, Fernanda Viégas, Oriol Vinyals, Pete Warden, Martin Wattenberg, Martin Wicke, Yuan Yu, and Xiaoqiang Zheng. TensorFlow: Large-scale machine learning on heterogeneous systems, 2015. Software available from tensorflow.org.

65. Cynthia L Smith and Janan T Eppig. The mammalian phenotype ontology: enabling robust annotation and comparative analysis. Wiley Interdisciplinary Reviews: Systems Biology and Medicine, 1(3):390–399, 2009.

66. Mouse genome informatics database. http://www.informatics.jax.org/downloads/reports/HOM_MouseHumanSequence.rpt. accessed 30 nov 2020.

67. Elizabeth I Boyle, Shuai Weng, Jeremy Gollub, Heng Jin, David Botstein, J Michael Cherry, and Gavin Sherlock. Go:: Termfinder—open source software for accessing gene ontology information and finding significantly enriched gene ontology terms associated with a list of genes. Bioinformatics, 20(18):3710–3715, 2004.

68. Yoav Benjamini and Yosef Hochberg. Controlling the false discovery rate: a practical and powerful approach to multiple testing. Journal of the Royal Statistical Society. Series B (Methodological), pages 289–300, 1995.

69. Arthur Liberzon, Aravind Subramanian, Reid Pinchback, Helga Thorvaldsdóttir, Pablo Tamayo, and Jill P Mesirov. Molecular signatures database (msigdb) 3.0. Bioinformatics, 27(12):1739–1740, 2011.

70. Alba Gutiérrez-Sacristán, Solène Grosdidier, Olga Valverde, Marta Torrens, Àlex Bravo, Janet Piñero, Ferran Sanz, and Laura I Furlong. Psygenet: a knowledge platform on psychiatric disorders and their genes. Bioinformatics, 31(18):3075–3077, 2015.

71. Oscar Franzén, Li-Ming Gan, and Johan LM Björkegren. Panglaodb: a web server for exploration of mouse and human single-cell rna sequencing data. Database, 2019, 2019.

72. Lawrence Hubert and Phipps Arabie. Comparing partitions. Journal of Classification, 2(1):193–218, 1985.

73. F. Pedregosa, G. Varoquaux, A. Gramfort, V. Michel, B. Thirion, O. Grisel, M. Blon- del, P. Prettenhofer, R. Weiss, V. Dubourg, J. Vanderplas, A. Passos, D. Cournapeau, M. Brucher, M. Perrot, and E. Duchesnay. Scikit-learn: Machine learning in Python. Journal of Machine Learning Research, 12:2825–2830, 2011.

74. Bo Li, Joshua Gould, Yiming Yang, Siranush Sarkizova, Marcin Tabaka, Orr Ashenberg, Yanay Rosen, Michal Slyper, Monika S. Kowalczyk, Alexandra-Chloé Villani, Timothy Tickle, Nir Hacohen, Orit Rozenblatt-Rosen, and Aviv Regev. Cumulus provides cloud- based data analysis for large-scale single-cell and single-nucleus rna-seq. Nature Methods, 17(8):793–798, Aug 2020.

75. Vincent Traag, L. Waltman, and Nees Jan van Eck. From louvain to leiden: guaranteeing well-connected communities. Scientific Reports, 9:5233, 03 2019.

76. Kevin Ushey, JJ Allaire, and Yuan Tang. reticulate: Interface to ’Python’, 2020. R package version 1.18.

77. Giampaolo Rodola. psutil: Cross-platform lib for process and system monitoring in Python., 2020. psutil 5.8.0.

78. Susan M Sunkin, Lydia Ng, Chris Lau, Tim Dolbeare, Terri L Gilbert, Carol L Thompson, Michael Hawrylycz, and Chinh Dang. Allen brain atlas: an integrated spatio-temporal portal for exploring the central nervous system. Nucleic acids research, 41(D1):D996– D1008, 2012.

79. Seurat3.0 finding integration vectors: long vectors not supported yet number 1029. https://github.com/satijalab/seurat/issues/1029. accessed 5 jan 2021.

80. Domenico De Berardis, Laura Orsolini, Nicola Serroni, Gabriella Girinelli, Felice Iasevoli, Carmine Tomasetti, Andrea de Bartolomeis, Monica Mazza, Alessandro Valchera, Michele Fornaro, et al. A comprehensive review on the efficacy of s-adenosyl- l-methionine in major depressive disorder. CNS & Neurological Disorders-Drug Targets (Formerly Current Drug Targets-CNS & Neurological Disorders), 15(1):35–44, 2016.

81. Loris A Chahl. Trp channels and psychiatric disorders. pages 987–1009. Springer, 2011.

82. J Craig Nelson, Carolyn M Mazure, and Peter I Jatlow. Desipramine treatment of major depression in patients over 75 years of age. Journal of clinical psychopharmacology, 15(2):99–105, 1995.

83. J Craig Nelson. Use of desipramine in depressed inpatients. The Journal of Clinical Psychiatry, 1984.

84. Matthew N Hill, Gregory E Miller, W-S Vanessa Ho, Boris B Gorzalka, and Cecilia J Hillard. Serum endocannabinoid content is altered in females with depressive disorders: a preliminary report. Pharmacopsychiatry, 41(2):48, 2008.

85. Julie A Blendy. The role of creb in depression and antidepressant treatment. Biological psychiatry, 59(12):1144–1150, 2006.

86. Jakob M Koch, Susanne Kell, Dunja Hinze-Selch, and Josef B Aldenhoff. Changes in creb-phosphorylation during recovery from major depression. Journal of psychiatric research, 36(6):369–375, 2002.

87. John Q Wang and Limin Mao. The erk pathway: molecular mechanisms and treatment of depression. Molecular neurobiology, 56(9):6197–6205, 2019.

88. Richard Day, Patricia A Ganz, and Joseph P Costantino. Tamoxifen and depression: more evidence from the national surgical adjuvant breast and bowel project’s breast cancer prevention (p-1) randomized study. Journal of the National Cancer Institute, 93(21):1615–1623, 2001.

89. Olga Szczesniak, Knut A Hestad, Jon Fredrik Hanssen, and Knut Rudi. Isovaleric acid in stool correlates with human depression. Nutritional neuroscience, 19(7):279–283, 2016.

90. Naomi Breslau, M Marlyne Kilbey, and Patricia Andreski. Nicotine dependence, major depression, and anxiety in young adults. Archives of general psychiatry, 48(12):1069– 1074, 1991.

91. Emmanuelle Goubert, Marc Altvater, Marie-Noelle Rovira, Ilgam Khalilov, Morgane Mazzarino, Anne Sebastiani, Michael KE Schaefer, Claudio Rivera, and Christophe Pel- legrino. Bumetanide prevents brain trauma-induced depressive-like behavior. Frontiers in molecular neuroscience, 12:12, 2019.

92. Elliot Ehrich, Ryan Turncliff, Yangchun Du, Richard Leigh-Pemberton, Emilio Fernan- dez, Reese Jones, and Maurizio Fava. Evaluation of opioid modulation in major depres- sive disorder. Neuropsychopharmacology, 40(6):1448–1455, 2015.

93. Giancarlo V De Ferrari and Nibaldo C Inestrosa. Wnt signaling function in alzheimer’s disease. Brain research reviews, 33(1):1–12, 2000.

94. RM Damian Holsinger, Catriona A McLean, Konrad Beyreuther, Colin L Masters, and Geneviève Evin. Increased expression of the amyloid precursor *β*-secretase in alzheimer’s disease. Annals of Neurology: Official Journal of the American Neurological Association and the Child Neurology Society, 51(6):783–786, 2002.

95. Boris Decourt, Debomoy K Lahiri, and Marwan N Sabbagh. Targeting tumor necrosis factor alpha for alzheimer’s disease. Current Alzheimer Research, 14(4):412–425, 2017.

96. Sumiti Saharan and Pravat K Mandal. The emerging role of glutathione in alzheimer’s disease. Journal of Alzheimer’s Disease, 40(3):519–529, 2014.

97. Jennifer C Palmer, Rachel Barker, Patrick G Kehoe, and Seth Love. Endothelin-1 is elevated in alzheimer’s disease and upregulated by amyloid-*β*. Journal of Alzheimer’s Disease, 29(4):853–861, 2012.

98. Yanfang Li, Hao Sun, Zhicai Chen, Huaxi Xu, Guojun Bu, and Hui Zheng. Implications of gabaergic neurotransmission in alzheimer’s disease. Frontiers in aging neuroscience, 8:31, 2016.

99. Karan Govindpani, Beatriz Calvo-Flores Guzmán, Chitra Vinnakota, Henry J Waldvogel, Richard L Faull, and Andrea Kwakowsky. Towards a better understanding of gabaergic re- modeling in alzheimer’s disease. International journal of molecular sciences, 18(8):1813, 2017.

100. Johanna Michael, Julia Marschallinger, and Ludwig Aigner. The leukotriene signaling pathway: a druggable target in alzheimer’s disease. Drug discovery today, 24(2):505– 516, 2019.

